# A replication-centered phylogeny illuminates the evolutionary landscape of bacterial plasmids

**DOI:** 10.1101/2024.09.03.610885

**Authors:** Yosuke Nishimura, Kensei Kaneko, Tatsuya Kamijo, Nanako Isogai, Yoshiki Morishita, Maho Tokuda, Hui Xie, Yasuhiro Tanizawa, Yusuke Tsuda, Aki Hirabayashi, Ryota Moriuchi, Hideo Dohra, Kazuhide Kimbara, Chiho Suzuki-Minakuchi, Hideaki Nojiri, Haruo Suzuki, Masato Suzuki, Masaki Shintani

## Abstract

Plasmids are amongst the most influential engines of bacterial evolution and horizontal gene transfer, fueling the global spread of traits such as antimicrobial resistance. Their deep evolutionary relationships, however, remain difficult to resolve because current classification schemes are constrained by host range and nucleotide similarity. Replication initiation proteins (RIPs), which govern plasmid persistence and diversification, also remain poorly annotated across public databases. Here we establish PInc, a curated and experimentally grounded replicon classification framework anchored in historically defined incompatibility groups of *Pseudomonas* plasmids. Homology searches beyond PInc revealed that most replication initiators analyzed here share a conserved winged-helix (WH) domain, defining a broad WH RIP superfamily. Using the conserved WH region, we reconstructed a large-scale phylogeny that linked WH RIPs to over 100,000 plasmids, representing approximately half of those analyzed across public databases. This phylogeny resolved eight major clades and the deep split between the single– and double-winged-helix superclades, while revealing clade-specific host and environmental distributions and substantial RIP diversity not captured by current typing tools or annotation schemes. Together, these results overcome the historical host bias of plasmid typing and provide a replication-centered view of plasmid diversification across bacterial lineages and environments.

## INTRODUCTION

Plasmids are among the most influential mobile genetic elements in bacteria, driving rapid evolutionary innovation through horizontal gene transfer^1,2^. They are also major vehicles for the global spread of antimicrobial resistance genes (ARGs), representing a critical threat to public health^3–5^. Recent large-scale sequencing efforts have revealed an unexpectedly vast and heterogeneous collection of plasmids^6,7^,underscoring the need for a systematic framework to organize this diversity and interpret plasmid evolution coherently.

A historical scheme known as incompatibility (Inc) groups provided the conceptual foundation for plasmid classification by exploiting phenotypic traits, defined by the inability of plasmids with similar replication systems to stably coexist in the same host^8^. More recently, sequence-based approaches have incorporated Inc group names into their classification schemes, improving classification coverage, but still falling short of delivering a macroevolutionary perspective. Replicon typing^9^ and plasmid taxonomic units (PTUs)^10,11^ rely on nucleotide similarity of rep genes or whole plasmid sequences, respectively, which obscures remote homology among deeply divergent plasmids. Relaxase-based approaches^10,12–14^ have expanded classification based on detectable relaxase homology, but exclude more than half of plasmids that lack relaxase genes^8^, and therefore cannot provide a universal backbone for plasmid classification. Consequently, current schemes do not fully resolve how plasmids are related across deep evolutionary time. By contrast, in virology, higher-order classification has increasingly been anchored by conserved hallmark genes, enabling molecular evolutionary analysis to be incorporated directly into large-scale taxonomic frameworks^15,16^. This contrast highlights the absence of an equivalent organizing principle for plasmid classification.

*Pseudomonas* plasmids provide an ideal starting point because this genus spans an exceptionally wide ecological and clinical range from soils and aquatic environments to hospital settings, and its plasmids serve as major vectors connecting environmental and clinical antimicrobial-resistance (AMR) reservoirs^17^. Phenotype-based 14 Inc groups (IncP-1 through IncP-14) have been designated for *Pseudomonas* plasmids and include some of the most clinically significant resistance plasmids identified in *P. aeruginosa*^18,19^. However, despite their importance, some are not only under-characterized but also lack nucleotide sequence information, relying solely on their original incompatibility phenotypes.

Here, we establish a curated and experimentally supported replicon classification system, termed PInc (*<u>P</u>seudomonas* Incompatibility-informed replico<u>n</u> <u>c</u>lassification), which provides a sequence-based framework anchored in historical Inc groups. This framework is grounded in the experimental identification of RIPs from representative *Pseudomonas* plasmids. Focusing on RIPs, we identified a conserved but distantly homologous winged-helix (WH) domain in most PInc groups and many additional plasmids, pointing to a shared evolutionary origin. Building on this observation, we reconstructed an extended phylogeny of WH-containing RIPs and used it to place many previously unclassified plasmids into a unified evolutionary framework. In this way, our study provides a replication-centered basis for higher-order plasmid classification and reveals the diversification and macroevolutionary trajectories of plasmids through their replication systems.

## Results and Discussion

### Sequencing and molecular characterization of *Pseudomonas* plasmids

We determined the complete nucleotide sequences of six *Pseudomonas* plasmids: Rms139 (IncP-2)^20^, Rms163 (IncP-5)^21,22^, Rsu2 (IncP-9)^23,24^, RP1-1 (IncP-11)^25,26^, R716 (IncP-12)^27^, and pMG26 (IncP-13)^28,29^. Although these plasmids represent the archetype members of their respective Inc groups in *Pseudomonas*, no full-length reference sequences had been available, leaving the replication systems of historically important plasmids unresolved (Table 1). For IncP-10 plasmids, because only a partial plasmid sequence of the representative plasmid R91-5^30,31^ is available, we selected pPAB546 as a representative (Supplemental Text S1). Sequencing of pMG26, previously assigned as IncP-13 plasmid^32^, yielded no circular contigs after removal of reads mapping to the *P. aeruginosa* PAO1 chromosome. Detailed molecular characterization indicated that pMG26 is not a plasmid but an integrative and conjugative element (ICE), and therefore IncP-13 plasmids should be regarded as ICEs (Supplemental Text S1). The other five (Rms139, Rms163, Rsu2, RP1-1, and R716) are shown to be self-transmissible plasmids (Supplemental Text S1). The RIPs and origin of vegetative replication (*oriV*) of these plasmids (Table 1) were identified by sequence– and structure-based searches, and the replication function of the selected predicted RIP– *oriV* regions was experimentally validated using mini-replicon constructs (see Methods, Figure S1, S2, and S3, and Supplemental Text S1).

**Table 1.**
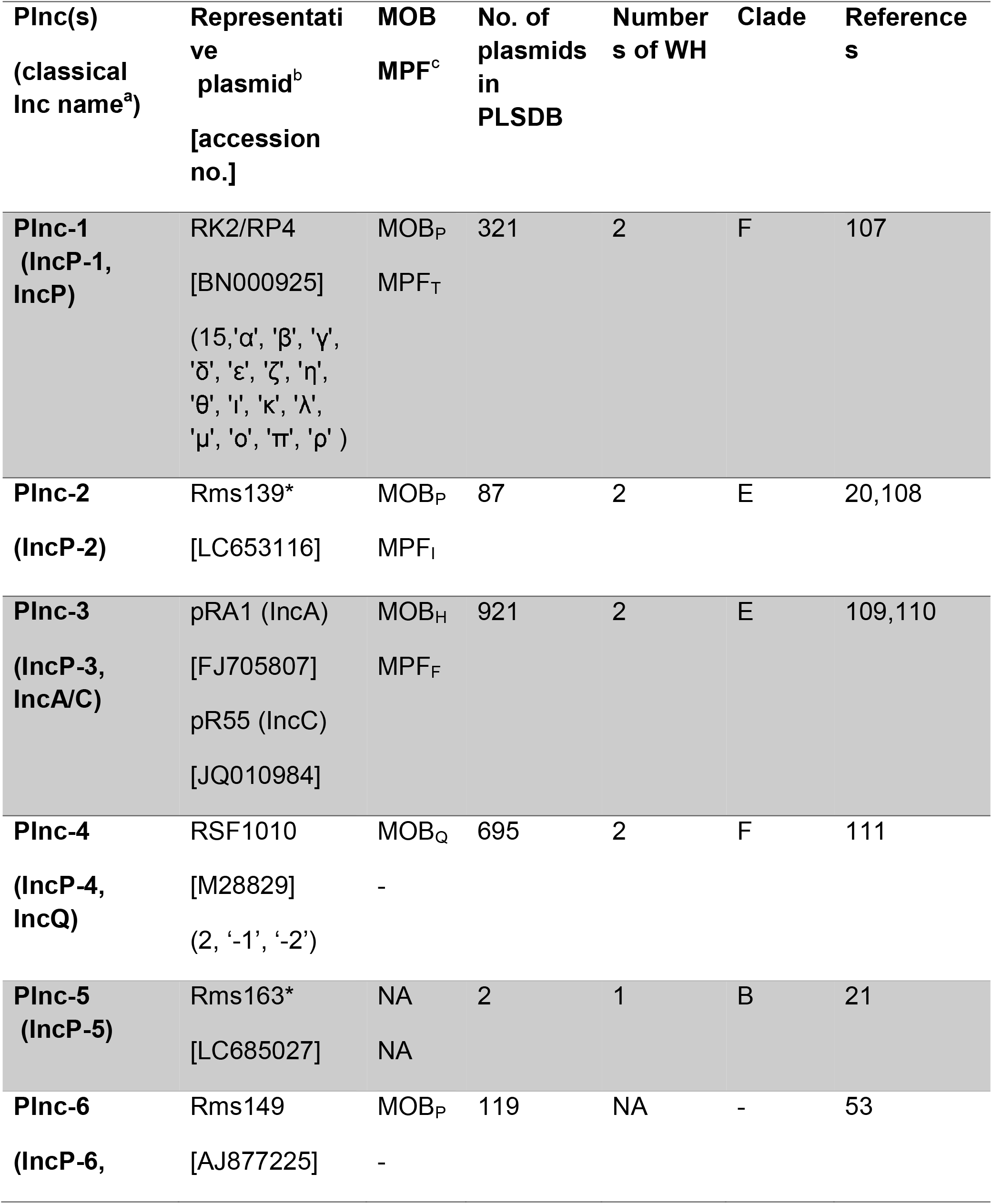
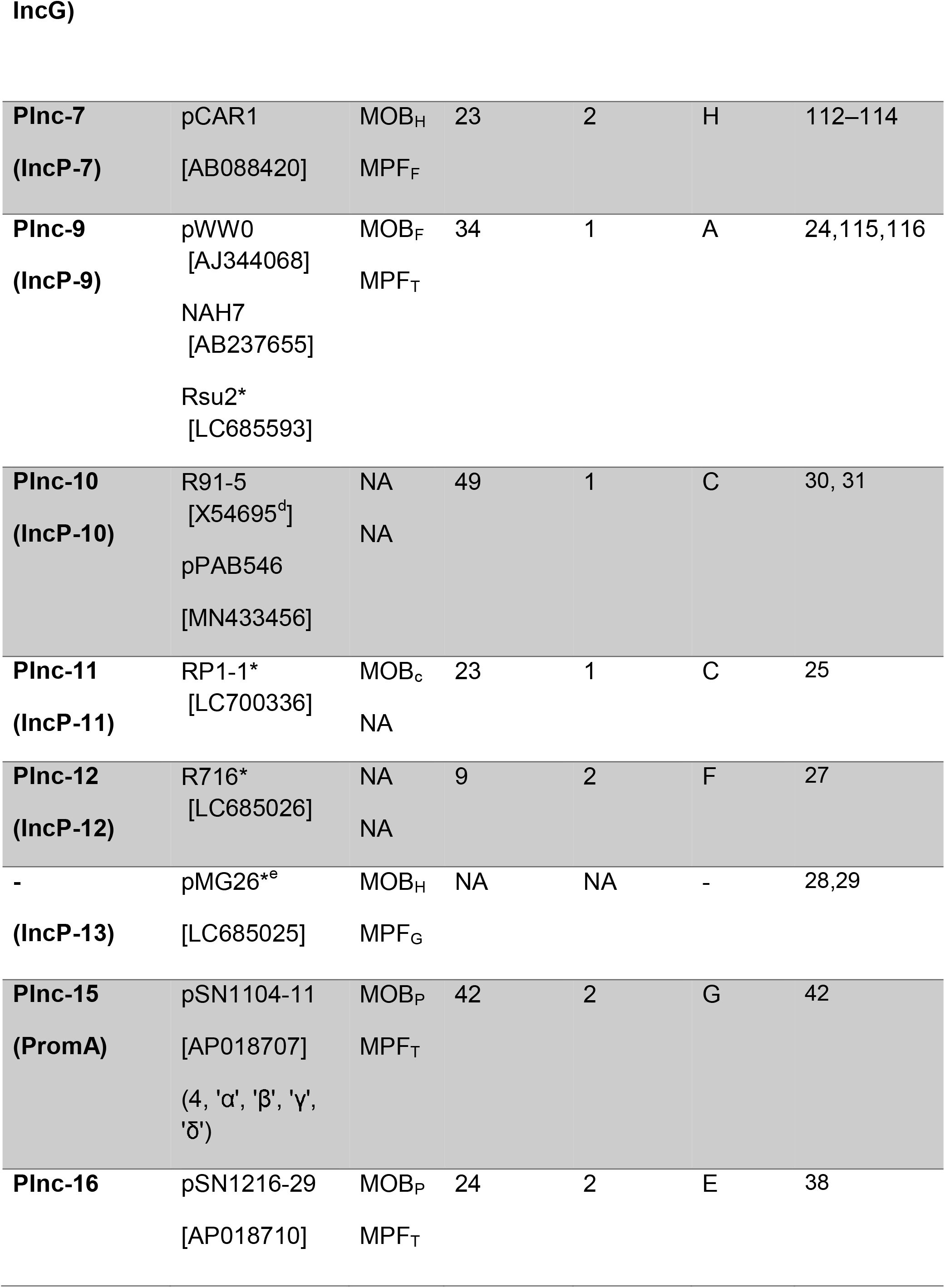
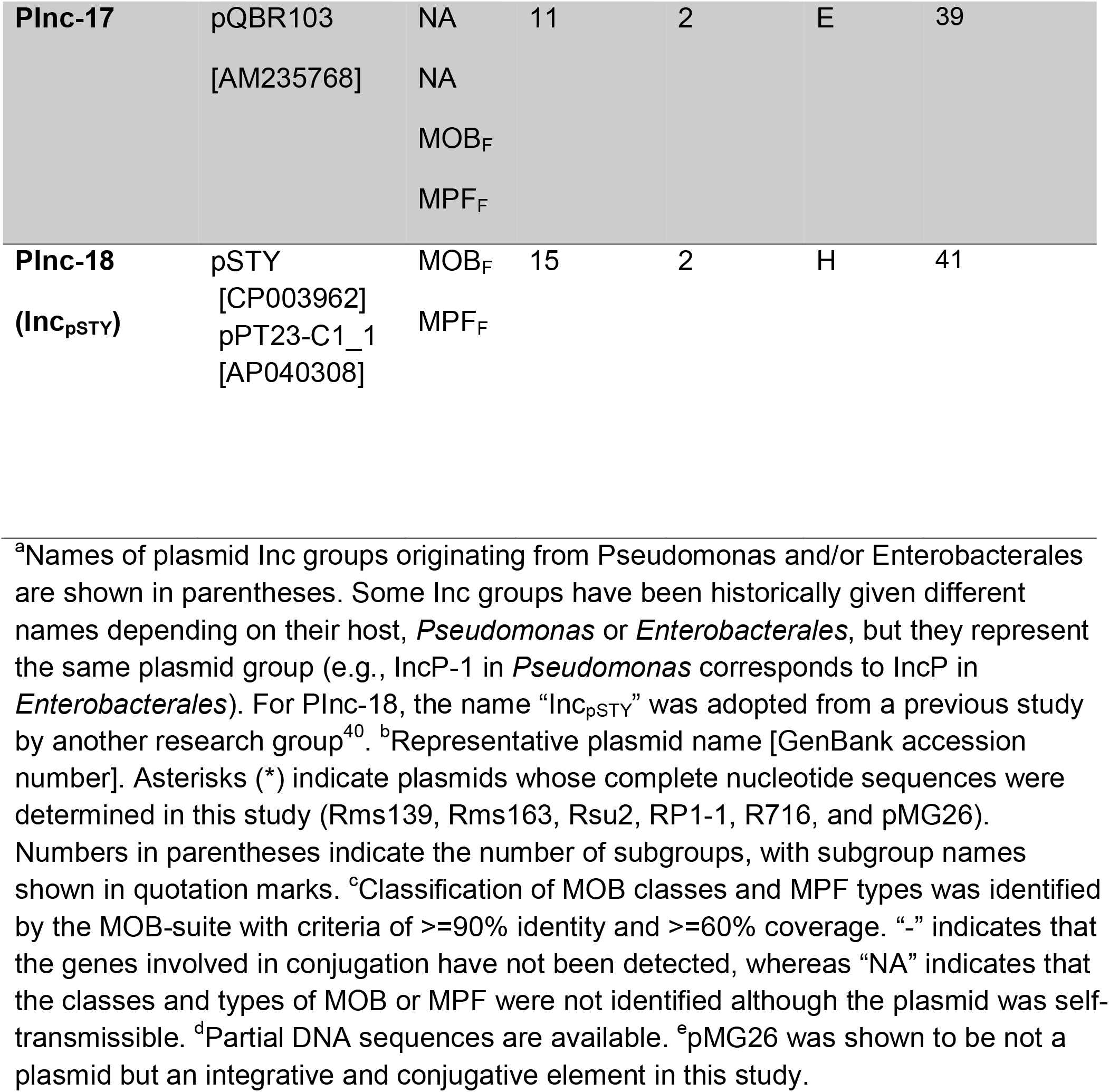
Lists of representative plasmids in the PInc groups.

### PInc classify unclassified plasmids in PLSDB

To provide a biologically grounded and sequence-based framework for *Pseudomonas* plasmids, we constructed a novel replicon-typing scheme named PInc, using experimentally supported RIPs as reference sequences (Table 1). This scheme follows the traditional Inc group numbering and its subgroup lettering (in Greek character) —for instance, PInc-1α corresponds to IncP-1α, and PInc-2 to IncP-2. Because the original Inc were defined on the basis of phenotypic incompatibility in different host lineages such as *Pseudomonas* and Enterobacterales, their names often overlap each other. For example, *Pseudomonas* IncP-1 is equivalent to Enterobacterales IncP (Table 1). Introducing the PInc scheme allows us to retain meaningful correspondence with the original Inc names while eliminating confusion caused by the overlapped or inconsistent Inc labels^33^.

Some Inc groups were excluded from PInc, because IncP-8^34^ and IncP-13 (this study) were identified as ICEs, or due to material unavailability for IncP-14^35^. Additionally, certain *Pseudomonas* plasmid groups that are not included in the traditional Inc grouping were incorporated into the PInc system, as they are widely distributed and known to disseminate antimicrobial resistance. These include PInc-15 (PromA group^36,37^), PInc-16 (pSN1216-29-like group^38^), PInc-17 (pQBR103-like group^39^), and PInc-18 (pSTY-like group^40,41^) (Table 1, Supplemental Text S1). PInc subgroups were defined according to subgroups of the Inc groups or newly defined if a group contains a large nucleotide diversity of the RIP genes (PInc-1, PInc-4, and PInc-15).

The replication regions of PInc-15 and PInc-16 had been experimentally characterized previously^38,42,43^, whereas those of PInc-17 and PInc-18 were characterized in the present study. For PInc-17, two *repA–oriV* regions had previously been predicted in pQBR103^39^, but their replication function had not been experimentally tested. Both regions supported replication in mini-replicon assays (Figure S1 and Supplemental Text S1). We designated one of these, *repP– 17A–oriV*, as the representative PInc-17 replication region because, although plasmids carry both regions, plasmids carrying only one of the two have so far been found exclusively with *repP-17A–oriV*. We also identified and experimentally validated the *repP-18A–oriV* region of PInc-18 using a mini-replicon assay (Figure S1 and Supplemental Text S1).

Plasmid sequences were collected and the RIP genes in the plasmids were identified based on sequence and structural homology to known RIPs (Figures S1, S2). Reference plasmids (n = 87), including those newly sequenced in this study, were selected to cover the 15 original Inc groups and 22 subgroups (Table S1) and their RIP genes were extracted and compiled into the repP database (n = 87, https://doi.org/10.6084/m9.figshare.31883911). Each PInc group and its subgroups were defined as follows: a plasmid was assigned to a group if its RIP gene showed nucleotide similarity to a reference RIP (BLASTn; ≥90% identity and ≥60% length coverage), considering only the best hit. Plasmids harboring multiple distinct RIPs were recognized as multi-replicons (e.g., PInc-10 and PInc-11).

Accordingly, we assigned 2,351 plasmids in PLSDB (59,895 plasmids) to PInc based on RIP genes detected using the repP database (Table S2), including 530 plasmids unclassified by PlasmidFinder and 1,514 unclassified by MOB-suite. The detected RIP gene sequences were curated into the PseudomonasRepDB database after redundancy removal (Table 2, n = 404, https://doi.org/10.6084/m9.figshare.26778175). Subsequent analysis using PseudomonasRepDB v1.1 assigned 2,366 plasmids to PInc (https://doi.org/10.6084/m9.figshare.26778175), including 535 plasmids unclassified by PlasmidFinder. Thus, PInc substantially expands the coverage of replicon-based classification for *Pseudomonas*-associated plasmids, increasing the number of classifiable plasmids by 29.1 % for PlasmidFinder and 181% for MOB-suite. Notably, PInc-classified plasmids span not only *Pseudomonas* but also diverse *Enterobacterales* hosts, reflecting the broad host range of several PInc plasmids (https://doi.org/10.6084/m9.figshare.26768173). Overall, PInc provides a systematic sequence-based framework that improves classification depth and coverage across plasmids in PLSDB.

**Table 2.** Representative plasmid replicon databases and their sequence features.

| **Host bacteria** | **Database** | **Latest version** | **Sequence** | **Coding region** | **Reference** |
|---|---|---|---|---|---|
| ***Enterococcus* spp.**<br>***Staphylococcus* spp.**<br>***Enterobacterales*** | PlasmidFinder | v2.1<br>(Apr 19, 2025) | 488 | Full-length, partial, or not included | <sup>9</sup> |
| ***Enterococcus* spp.**<br>***Staphylococcus* spp.**<br>***Enterobacterales*** | MOB-typer* | v3.1.9<br>(Jun 4, 2024) | 2,685 | Full-length, partial, or not included | <sup>85,117</sup> |
| ***Acinetobacter* spp.** | AcinetobacterPlasmidTyping | v3.0<br>(Feb 12, 2025) | 555 | Full-length | <sup>60</sup> |
| ***Pseudomonas* spp.** | PseudomonasRepDB | v1.1<br>(This study) | 404 | Full-length, or partial | This study |
| **All bacteria** | WHRepDB | v1.1<br>(This study) | 8,983 | Partial (WH domain) | This study |

\*The “IncP” labels by the MOB-typer database should be interpreted with caution. In some cases, sequences are reported as “IncP”, although they do not correspond to the correct incompatibility group classification<sup>32</sup>.

### Comparative genomics analysis highlights PInc group characteristics

To characterize the genomic and evolutionary diversity of plasmids classified by PInc, we conducted a comprehensive pangenomic analysis across the PInc groups (Tables S2, S3-1∼S3-14). Three key patterns emerged. First, several PInc groups (notably PInc-2, PInc-17, and PInc-18) exhibited exceptionally large genome sizes (Figure 1A). Second, specific groups including PInc-4, PInc-10 and PInc-11, displayed distinctive multi-replicon coupling patterns, suggesting long-term modular evolution of their replication systems. Third, PInc-1 plasmids displayed substantial variation in plasmid size, GC content, and RIP sequence, although the extent of variation differed among these features (Figure 1AB, Table S2 and S4).

**Figure 1.**
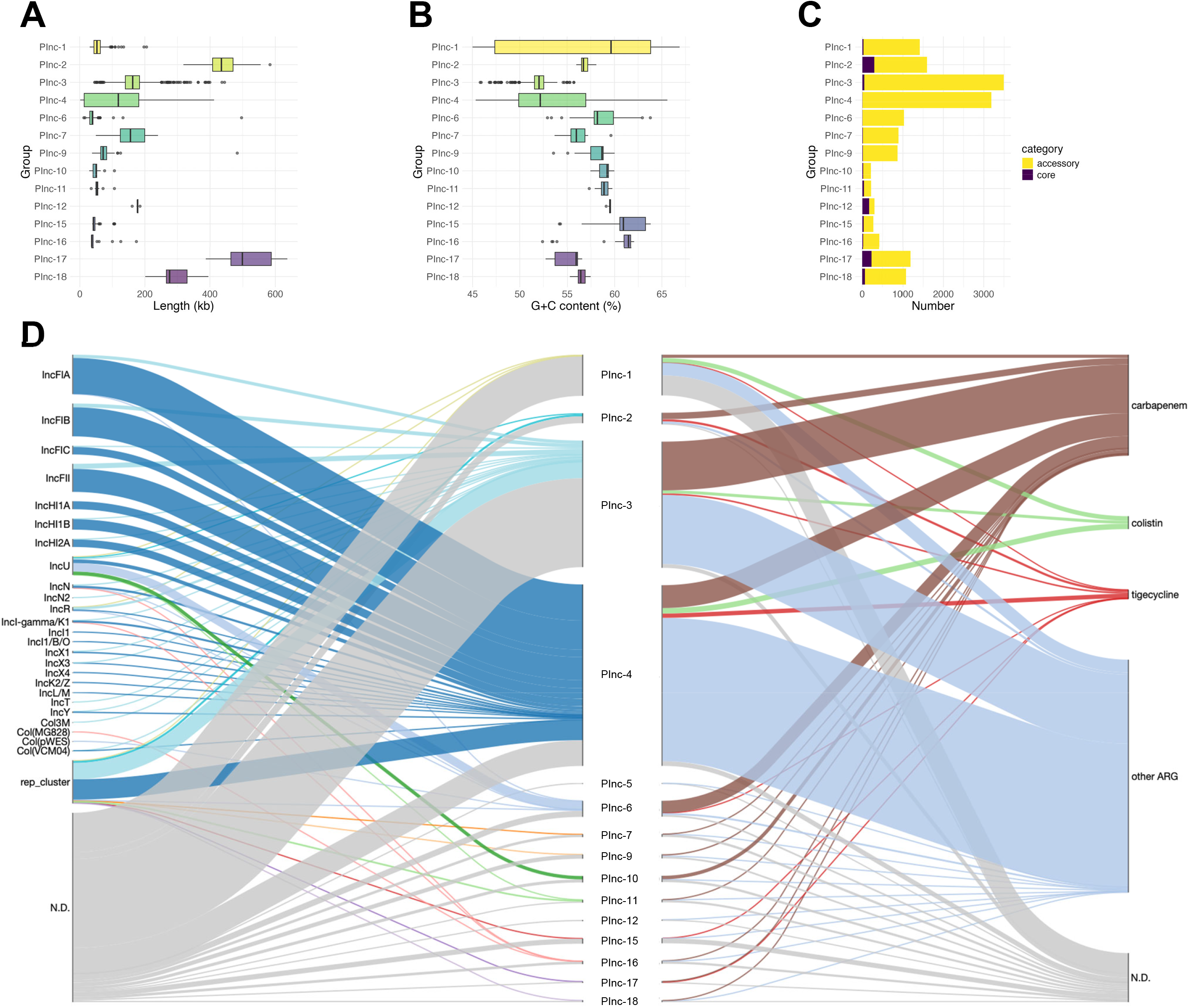
Features of plasmids of PInc groups in the PLSDB. Distribution of (A) plasmid sizes (kb) and (B) GC contents (%), and (C) numbers of core and accessory genes of each PInc plasmids in the PLSDB (n = 2,351). (D) Relationship of each PInc plasmids to other plasmid replicons (left) and ARGs (right) in the PLSDB. “rep_cluster” contains different types of replicons identified by MOB-typer. ‘N.D.’ indicates not detected. A sankey diagram was generated using the SankeyWidget library in Python through in-house programming.

Plasmid size distribution across PInc groups (Figure 1A) revealed that PInc-2, PInc-17, and PInc-18 included relatively large plasmids. However, plasmid size alone does not indicate a greater proportion of accessory genes and the relative contributions of conserved backbone genes and accessory genes differed among these groups (Figure 1C). The broad GC-content distributions of PInc-1 and PInc-4 (Figure 1B) may reflect their occurrence in diverse host backgrounds^44–46^ and/or the association of related replication modules with otherwise divergent plasmid backbones. The latter possibility may be particularly relevant to PInc-4, which showed a broad plasmid-size distribution (Figure 1A) and a small conserved core (Figure 1C). Core-genome analysis further showed that PInc-2, PInc-12, and PInc-17 had relatively large conserved core sets, whereas, PInc-1 had a comparatively small core-gene proportion (Figure 1C). Here, we define a multi-replicon plasmid as a single plasmid carrying two or more distinct replication systems, detected as distinct RIP or replicon assignments. Figure 1D summarizes the additional replicon types detected in plasmids carrying each PInc replicon. Multi-replicon plasmids were frequent in PInc-4 (70.9%), PInc-11 (65.2%), PInc-6 (60.5%), and PInc-10 (51.0%), whereas more than 90% of PInc-1, PInc-5, PInc-7, PInc-9, and PInc-15 plasmids carried only a single detected replicon (Figure 1D, Table S2). The associated replicon types also differed among groups. PInc-4 plasmids were associated with a wide variety of additional replicons, whereas PInc-10 and PInc-11 plasmids were most often associated with each other or with a limited set of additional replicons, including IncU or rep_cluster_2025 (Figure 1D, Table S2). More than 60% of the PInc-2, PInc-3, PInc-4, PInc-6, PInc-10, and PInc-17 plasmids harbored ARGs. Among these groups, PInc-2, PInc-6, PInc-10, and PInc-17 included plasmids carrying clinically important ARGs, conferring resistance to carbapenems, tigecycline, or colistin (Figure 1D). More detailed pangenomic characterization for each PInc was provided in Supplemental Text S1.

### Homology-guided collection of RIP sequences expands the diversity of plasmid initiators

To investigate the evolutionary relationships of RIPs, we first assessed protein sequence homology among 251 non-redundant RIP sequences from PInc plasmids using BLASTp (Figure S3). Several PInc groups exhibited substantial sequence diversity, particularly PInc-1, PInc-4, and PInc-15, for which the most divergent RIP pairs shared only 34.05%, 28.81%, and 56.14% amino acid sequence identity, respectively. By contrast, the minimum pairwise identities within each of the remaining PInc groups exceeded 73% (Table S4). BLASTp also detected homologous RIP pairs among specific combinations of groups: among PInc-2, PInc-3, and PInc-17 (31.1– 58.7% identity); between PInc-1 and PInc-12 (20.5–27.5%); and between PInc-10 and PInc-11 (50.7–53.4%). Thus, the observed sequence similarities between these groups suggest that they are evolutionarily related.

To extend the analysis beyond the sensitivity of BLASTp, we next performed iterative sequence searches using RIPs from the PInc groups as queries to collect more diverse homologs from PLSDB and the UniProt Archive (UniParc). To improve coverage of RIP sequence diversity, we also included eight additional RIPs outside the PInc groups as additional queries (Table S5). The detailed procedures are described in Supplemental Text S1 and Figure S4. In total, the RIPs identified using this approach were classified into 14 homologous groups and then clustered at 50% amino acid sequence identity within each group, yielding 5,345 representative sequences. This expanded sequence set provided the basis for resolving deeper relationships among plasmid RIPs.

### Remote homology analysis reveals evolutionary divergence among plasmid RIPs

Remote homology among RIPs from the 14 homologous groups was assessed using HHblits^47^, which performs sensitive sequence-based homology detection through HMM-HMM comparison. With the exception of G-PInc-6, which comprises sequences collected using PInc-6 RIPs as queries, RIPs from the remaining 13 groups shared a homologous region (Figure S5). Based on the significance of homology among groups, these 13 groups could be further divided into two supergroups. One supergroup, comprising seven groups, showed homology with >95% probability in 20 of 21 possible pairwise group comparisons, whereas the other, comprising six groups, did so in all 15 possible comparisons (Figure S5).

Considering available experimental structures and AlphaFold3^48^ predictions of RIPs (Figure S6), we classified these two supergroups as single winged-helix (sWH) and double winged-helix (dWH) according to their domain architecture. The sWH supergroup comprises seven groups with a single WH domain, whereas the dWH supergroup comprises six groups with two WH domains. In dWH RIPs, available structures of RepE (plasmid F; PDB: 2Z9O), Pi (R6K; PDB: 2NRA), RepA (pPS10; PDB: 1HKQ), and TrfA (RK2/RP4)^49^ all contain two conserved winged-helix domains, here termed WH1 and WH2. By contrast, only two crystal structures are available for sWH RIPs^50^ (pSK41, PDB: 5KBJ; pTZ2162, PDB: 4PT7), although AlphaFold3 predicts that the remaining members also adopt a single-WH-domain architecture. HHblits detected significant homology (>80% probability) between the sWH and dWH supergroups in 18 of 42 possible pairwise group comparisons (6 × 7). In all cases, the homologous region mapped to the WH domain of sWH RIPs and to WH2, the C-terminal WH domain, of dWH RIPs.

By contrast, RIPs of G-PInc-6 showed no detectable homology to any other groups by HHblits, with probabilities of less than 1% in all comparisons. We therefore examined their evolutionary origin separately by searching against the Pfam database using HHblits. G-PInc-6 RIPs matched PF03090, corresponding to the replicase family, with 98.4% probability. This domain is present in the ColE2 RIP^51^, a member of the archaeo-eukaryotic primases (AEPs)^52^ superfamily. Consistent with this assignment, G-PInc-6 RIPs^53,54^ also contain a PriCT domain implicated in DNA unwinding^51^. Together, these observations identify G-PInc-6 RIPs as AEP-superfamily initiators and therefore as evolutionarily distinct from the WH-containing RIPs.

On the basis of this shared homologous domain across the sWH and dWH supergroups, we define these proteins as a WH RIPs superfamily. This framework also provides a basis for interpreting their deeper evolutionary relationships. A previous structure-based study reported deep evolutionary relationships among helix-turn-helix (HTH) domains, including WH domains^55^, although evolutionary relationships across diverse families of plasmid WH RIPs were not systematically investigated. Moreover, the N-terminal WH1 domain of dWH RIPs has been proposed to be evolutionarily related to the WH domain of archaeal Orc1/Cdc6 and eukaryotic Orc proteins^56^, both of which function in origin recognition for the chromosomal replication^57^. By contrast, the evolutionary origin of WH2 has remained unresolved. Our analysis identifies homology between WH2 and the WH domain of sWH RIPs, uncovering an evolutionary link within plasmid replication initiators. Notably, the WH2 domain of pPS10 RepA binds the 3′ end of the iteron and is the major determinant of sequence specificity^56,58^, suggesting that this conserved WH module represents a core origin-recognition element across the WH RIP superfamily. More broadly, the coexistence of such divergent RIPs lineages, including both WH RIPs and AEP-superfamily initiators, within the traditional Inc framework exposes the limitations of historical phenotypic classification and supports the need for a sequence-based PInc system. The conserved WH module across the 13 homologous groups then provided a common anchor for large-scale phylogenetic reconstruction.

### WH RIP phylogeny reveals replication initiator diversity in over 100,000 plasmids

We reconstructed a phylogeny of WH RIPs from the 13 homologous groups using HHblits-defined homologous regions spanning the conserved WH domain and its flanking sequences. The maximum-likelihood tree was built from 4,769 representative sequences clustered at 50% identity (Figure 2A, Figure S4B). On the basis of branch support, we defined eight clades (A to H) and 42 subclades using high-confidence thresholds (UFBoot2: ≥95% or SH-aLRT: ≥80%) (Figure 2B). At the deepest level, sWH and dWH RIPs formed two superclades separated by a major split in the tree, supported by both UFBoot2 (88%) and SH-aLRT (89.4%). This split was also consistently identified as a phylogenetic root by minimal ancestor deviation analysis^59^. These results define a deeply structured WH RIP phylogeny and suggest an early evolutionary transition involving either gain of an additional WH domain or loss of one ancestral WH domain.

**Figure 2.**
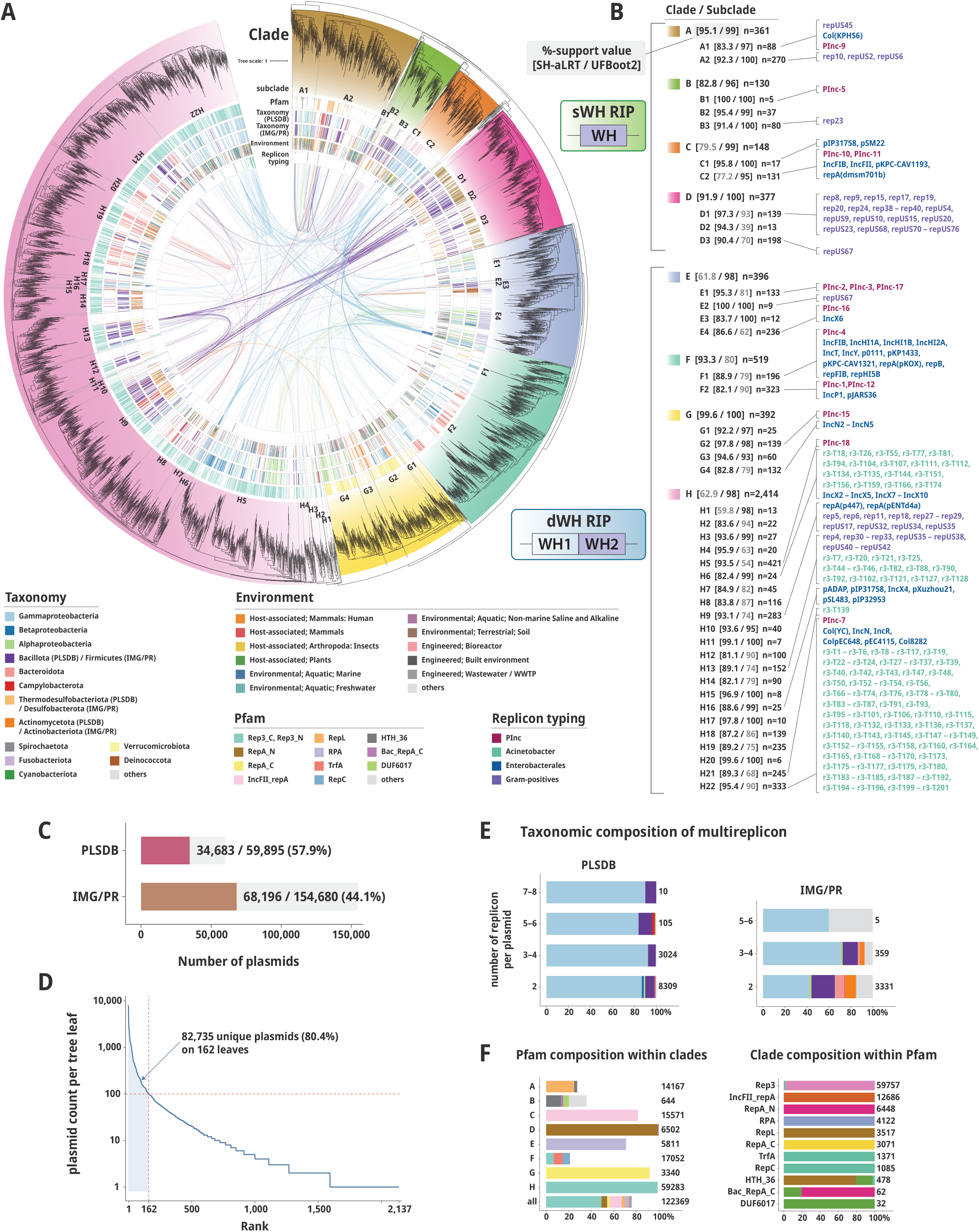
Large-scale phylogeny of WH RIPs and its links to plasmid diversity across public databases. (A) Maximum-likelihood phylogeny of WH RIPs reconstructed from HHblits-defined homologous regions spanning the conserved WH domain and its flanking sequences. The tree was built from 4,769 representative sequences clustered at 50% identity. Annotation rings show clade and subclade assignments together with stacked-bar summaries of Pfam annotation, taxonomy, environment, and replicon-typing information associated with linked plasmids. Curved lines drawn in the centre connect WH RIPs co-occurring in the same multi-replicon plasmids; line colours indicate the taxonomy of the corresponding plasmid hosts. Links for plasmids without taxonomic assignment were omitted from this visualization. (B) Summary of the eight clades (A–H) and 42 subclades. Numbers in brackets indicate support values (%) for major nodes in the order SH-aLRT/UFBoot2. Support labels are shown in light gray when either the SH-aLRT value is <80% or the UFBoot2 value is <95%. Representative PInc groups and other known replicon types associated with each clade are indicated. (C) Numbers and proportions of plasmids in PLSDB and IMG/PR linked to leaves in the WH RIP phylogeny. (D) Rank-abundance distribution of linked plasmids per tree leaf. (E) Taxonomic composition of multi-replicon plasmids carrying multiple WH RIPs in PLSDB and IMG/PR, stratified by the number of WH replicons per plasmid. Gray indicates plasmids without taxonomic assignment; in IMG/PR, all gray fractions correspond to taxonomy-undefined plasmids. (F) Pfam annotation profiles of phylogeny-linked RIPs across clades (left) and clade composition within each Pfam category (right). Sample sizes are indicated next to the bars.

We next placed this phylogeny in the context of plasmid diversity by surveying associated plasmids in the public databases PLSDB and IMG/PR. PLSDB is enriched in plasmids from cultivated isolates, whereas IMG/PR is largely composed of metagenome-derived plasmids. Protein sequences predicted from 214,575 presumably complete plasmids (59,895 from PLSDB and 154,680 from IMG/PR) were mapped to the closest leaves in the WH RIP tree (Figure 2C). This analysis identified 122,369 phylogeny-linked RIPs associated with 102,879 plasmids (47.9% of all analyzed plasmids), with linked plasmids represented in all subclades. Linked plasmids were distributed broadly across the tree, but their abundance varied markedly among leaves. A total of 2,137 leaves (44.8% of all 4,769 leaves) were linked to plasmids, and 162 of these leaves (3.4%) were each associated with more than 100 plasmids (Figure 2D). Together, these leaves accounted for 82,735 plasmids (80.4% of phylogeny-linked plasmids). Of the linked plasmids, 15,131 (14.7% of phylogeny-linked plasmids) were multi-replicons carrying multiple WH RIPs (Figure 2E). Most of these multi-replicons were derived from Gammaproteobacteria, although other taxa, including Bacillota, Actinomycetota, Bacteroidota, and Campylobacterota, were also represented. Overall, these observations show that the WH RIP phylogeny captures broad plasmid diversity across all subclades.

We then asked how much of this diversity is captured by existing annotation frameworks. Of the 122,369 phylogeny-linked RIPs, 60,202 (49.2%) from 57,087 plasmids were newly identified and were not detected by existing replicon typing tools, including PlasmidFinder, MOB-suite, and AcinetobacterPlasmidTyping^60^, or by the Pinc framework defined here. When mapped onto the WH RIP phylogeny, replicon types were broadly distributed across clades (Figure 2B). In particular, RIPs from 2,768 PInc-assigned plasmids were distributed across 16 leaves spanning seven of the eight clades, except for clade D. Likewise, 44, 33, and 33 leaves were associated with Gram-positive, Enterobacterales, and *Acinetobacter* replicon types, respectively. In total, only 125 leaves (2.6% of all 4,769 leaves) were linked to current typing schemes, indicating that much of the RIP diversity remains uncaptured by existing plasmid typing frameworks. Furthermore, 29,600 RIPs (24.2%) from 26,133 plasmids lacked any Pfam domain annotation (Figure 2F), and in clades A, B, and F, more than half of the RIPs were unannotated. Together, these results indicate that a substantial fraction of RIP diversity remains uncaptured by both nucleotide-level replicon typing and protein-level domain annotation.

### Structural and sequence diversification across WH RIP clades

We examined the structural diversity of representative WH RIPs from PInc plasmids, focusing on domain organization and predicted three-dimensional structures (Figure 3 and Figure S7). Pfam-based domain annotation, together with WH1/WH2 assignments based on previous studies^49,61^ (see Methods), identified WH domains linked to conserved replication-associated domains, including RepA_N/C, Rep3_N/C, and IncFII_repA. The Bac_RepA_C domain, which does not overlap the WH region, has been implicated in interactions with DnaG primase^50^. On the basis of available crystal structures, we inferred that sWH RIPs bind DNA as dimers, whereas dWH RIPs bind DNA as monomers. These observations indicate that WH RIPs combine a conserved WH core with distinct replication-associated domain architecture across clades.

**Figure 3.**
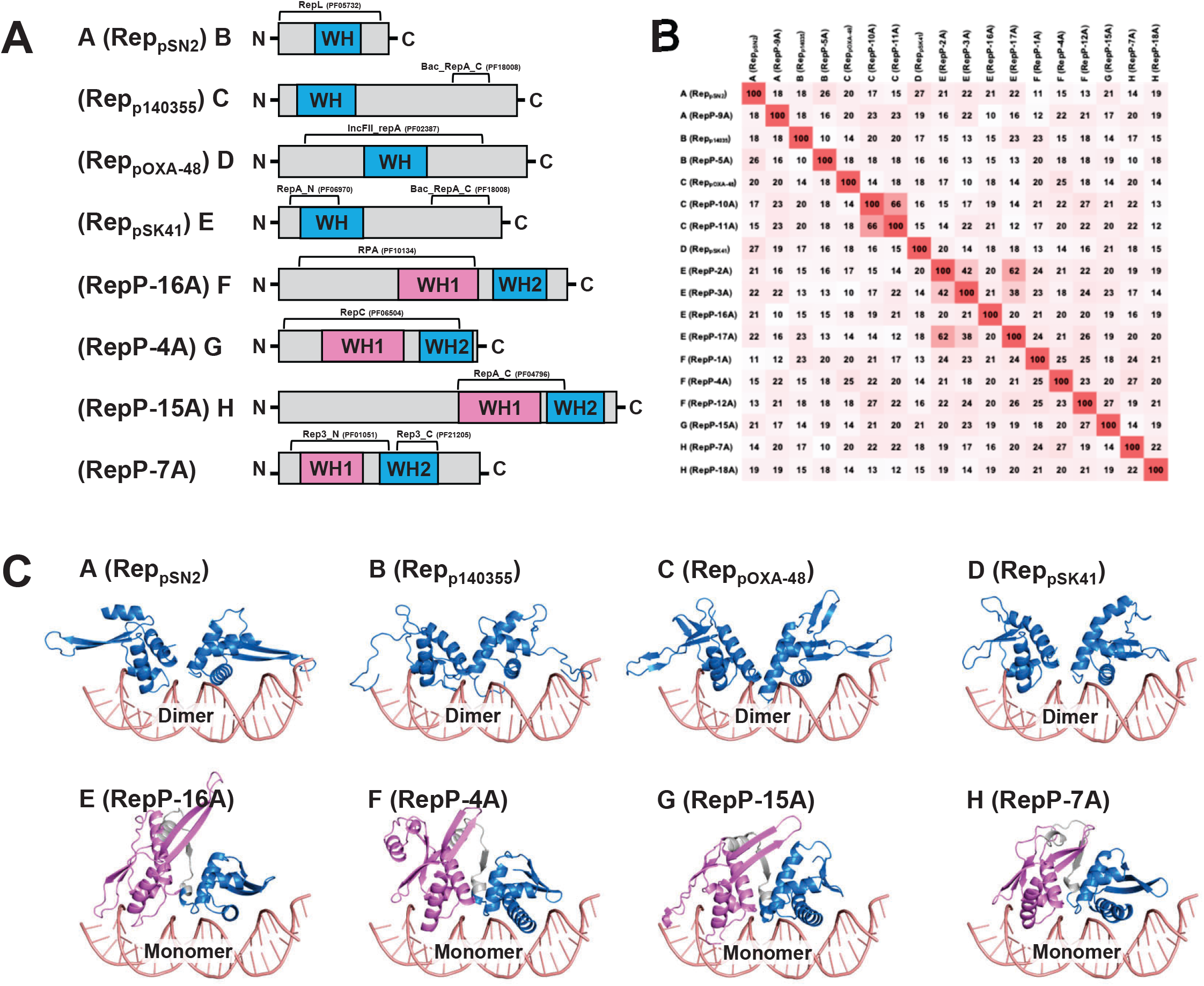
Domain architecture and predicted structural features of representative RIPs from the eight WH clades. (A) Domain organization of representative RIPs. RIPs A–D contain a single WH domain, whereas RIPs E–H contain two WH domains (WH1 and WH2). WH1 is shown in magenta; WH2 and single WH domains are shown in cyan. (B) Pairwise protein sequence identity among representative Rep proteins. (C) Predicted DNA-bound structures of the conserved regions corresponding to the WH domain shown in panel A. For RIPs A–D, the single WH domain is shown; for RIPs E–H, WH1 and WH2 together with the intervening linker are shown. WH1 is shown in magenta; WH2 and single WH domains are shown in blue. These are not full-length RIP structures. RIPs A–D are predicted to function as dimers, whereas RIPs E–H are predicted to function as monomers.

Sequence features derived from the multiple alignment used for phylogenetic reconstruction further distinguished clades A–H (Figure S8). In particular, both gap patterns in the alignment and the lengths of the N– and C-terminal regions flanking the conserved WH domain showed clade-specific features, while still exhibiting substantial variation within individual clades, such as clades C and E. Overall, these results highlight structural diversification of WH RIPs across clades, reflecting the diversification of replication systems underlying plasmid evolution.

### WH RIP phylogeny reveals host and environmental constraints on plasmid dispersal

We examined the taxonomic distribution of WH RIP clades using metadata from PLSDB and IMG/PR. PLSDB follows NCBI taxonomy, whereas IMG/PR follows GTDB taxonomy (Figure 4A). Despite differences in the taxonomic systems and data sources, several clade–taxonomy associations were consistent across both datasets. Clades B and D were strongly associated with Bacillota, clades C and F with Gammaproteobacteria, and clade E with Alphaproteobacteria. Clade H showed the broadest taxonomic distribution and, in IMG/PR, accounted for most Bacteroidota and large fractions of Campylobacterota and Fusobacteriota. This was particularly evident in IMG/PR, which contained 22,390 Bacteroidota plasmids, most of which were assigned to clades A and H. At the subclade level, however, clade H was partitioned into groups with distinct taxonomic associations, indicating that its broad host range is composed of multiple more restricted evolutionary units (Figure S9). Clade G also showed a broad host range, with strong representation in Actinomycetota. These patterns indicate that WH RIP evolution is structured by host lineage, while some clades have undergone broader host transitions than others.

**Figure 4.**
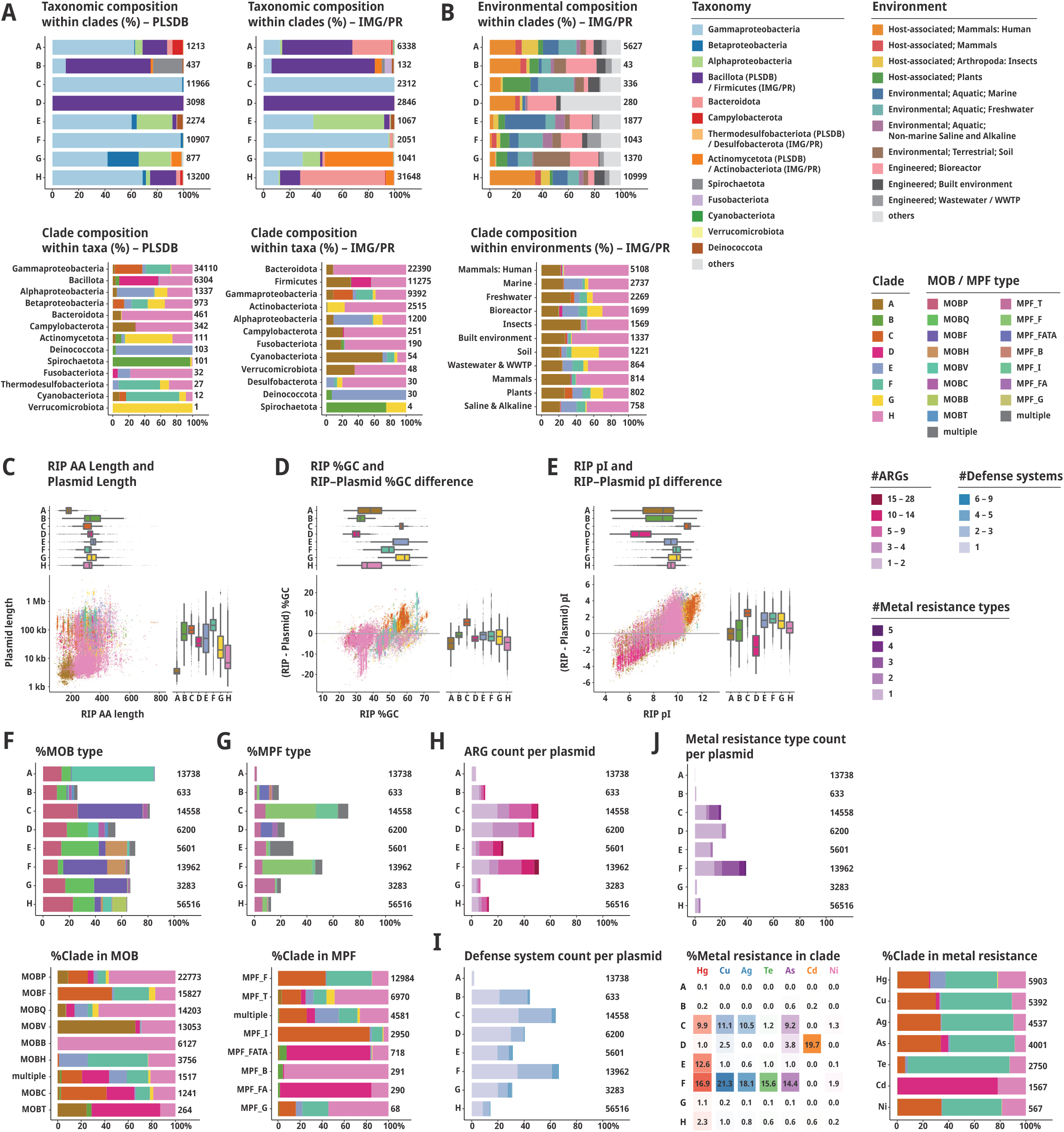
Taxonomic, ecological, compositional and functional diversification across the eight WH RIP clades. (A) Taxonomic distribution of WH RIP clades among linked plasmids in PLSDB and IMG/PR. Upper panels show taxonomic composition within each clade; lower panels show clade composition within each taxon. PLSDB follows NCBI taxonomy, whereas IMG/PR follows GTDB taxonomy. (B) Environmental distribution of WH RIP clades among IMG/PR-linked plasmids derived from metagenomic and metatranscriptomic datasets. Upper panel shows environmental composition within each clade; lower panel shows clade composition within each environment. (C) Clade-wise distributions of RIP amino-acid length and plasmid length. Paired RIP and plasmid lengths are jointly visualized in the scatter plot, and the boxplots summarize the distribution of each variable across clades. (D) Clade-wise distributions of RIP GC content and its difference from the GC content of the corresponding plasmid. The scatter plot jointly visualizes RIP GC content and its difference from the corresponding plasmid GC content, and the boxplots summarize the distribution of each variable across clades. (E) Clade-wise distributions of RIP isoelectric point (pI) and its difference from the median pI of the corresponding plasmid proteome. The scatter plot jointly visualizes RIP pI and its difference from the median pI of the corresponding plasmid proteome, and the boxplots summarize the distribution of each variable across clades. (F) Distribution of MOB classes across clades. Upper panel shows MOB-class composition within each clade; lower panel shows clade composition within each MOB class. (G) Distribution of MPF classes across clades. Upper panel shows MPF-class composition within each clade; lower panel shows clade composition within each MPF class. (H) Distribution of antimicrobial resistance (AMR) genes across clades, shown as the proportions of plasmids carrying different numbers of ARGs per plasmid. (I) Distribution of defense systems across clades, shown as the proportions of plasmids carrying different numbers of defense systems per plasmid. (J) Distribution of metal resistance genes across clades. Upper panel shows the proportions of plasmids carrying different numbers of metal resistance determinants per plasmid; lower panels show the proportions of individual metal resistance classes within each clade and clade composition within each metal resistance class. Sample sizes are indicated next to the bars.

For environmental distribution, we analyzed IMG/PR-linked plasmids, for which environmental metadata were available from the corresponding metagenomic and metatranscriptomic datasets. Although most WH RIP clades were represented across multiple habitat categories, their relative environmental distributions varied among clades (Figure 4B, upper panel). Clade H was broadly distributed across habitat categories (Figure 4B, upper panel) and accounted for a major fraction of RIPs across most environments (Figure 4B, lower panel). This broad distribution also masked substantial heterogeneity within clade H, as individual subclades showed more specific ecological associations (Figures 2A, S9). Animal-associated environments, including human, other mammals, and insects, as well as the built environment, were dominated by clades H and A (Figure 4B, lower panel). By contrast, marine, freshwater, soil, saline and alkaline environments, as well as plant-associated and bioreactor samples, showed more diverse clade mixtures, with increased contributions from clades C, E, F and G. Among these, clade E was more prominent in marine-associated plasmids, whereas clades C and F contributed more strongly to freshwater– and plant-associated environments, and clade G was more strongly associated with soil environments. These observations suggest that plasmid ecological distributions reflect clade-associated variation in dispersal and environmental association. We therefore asked whether clade-associated variation also extended to architectural, compositional and functional properties.

### Distinct architectural and functional profiles of the eight WH RIP clades

Several clade-associated trends were apparent in plasmid size, RIP length, GC composition and isoelectric point (pI) (Figure 4C–E). Clades A and H tended to occupy the lower end of the plasmid size range but differed substantially in RIP length (Figure 4C). Clade A was associated with the smallest plasmids and the shortest RIPs (median plasmid length 3.2 kb; median RIP length 185 aa), whereas clade H also tended to occur in relatively small plasmids (median 6.8 kb) but carried substantially longer RIPs (median 325 aa). By contrast, the remaining clades were associated with progressively larger plasmids, with median lengths ranging from 19.4 kb to 144 kb, indicating that plasmid size is itself a characteristic feature of WH RIP clades. Clade-associated trends were also observed in GC composition (Figure 4D). Clades B and D were associated with the lowest-GC plasmids (median 33.4% and 33.6%, respectively), which may reflect their taxonomic bias towards Bacillota, whereas clades E and G showed the highest plasmid GC values. RIP GC content broadly tracked plasmid-wide GC content, but clade C was distinctive in showing higher GC in RIPs than in their plasmid backgrounds, with a similar but weaker tendency in clades E, F and G. Differences in pI were even more pronounced (Figure 4E). Clade C showed a marked positive shift relative to the plasmid proteome, clade D showed the opposite trend, and clades E, F and G were more moderately shifted towards higher pI. The elevated pI of clade C RIPs is likely explained by enrichment of positively charged residues, particularly lysine and arginine, and may reflect compositional adaptation related to DNA binding. Together, these patterns reveal clade-associated trends in plasmid architecture, sequence composition and RIP biochemical properties. We then asked whether these clade-associated trends extended to transfer-related and accessory functions.

Mobility and accessory functions also varied sharply across WH RIP clades. MOBP and MOBQ were detected across most clades, whereas other MOB classes showed stronger clade specificity (Figure 4F). Notably, clade A was enriched in MOBV despite its association with small plasmids, suggesting retained mobilization capacity. By contrast, clades C, F and G were enriched in MOBF, whereas clades E and H showed relatively stronger contributions from MOBQ. MPF repertoires also differed among clades. MPF modules were nearly absent from clade A, whereas clades C and F were dominated by MPF_F and clade D by MPF_FATA (Figure 4G). AMR and defense systems showed similarly uneven distributions across clades (Figure 4H). Clades C, D and F were relatively enriched in ARG-rich plasmids, whereas clades A, B, G and H were dominated by plasmids with few ARGs. Defense system content showed a partly overlapping but distinct pattern, with clades B, C, D and F more frequently carrying multiple defense systems, whereas clades A and H showed low defense systems content (Figure 4I). Metal resistance genes also showed strong clade specificity (Figure 4J). Clades C and F were enriched in plasmids carrying multiple metal resistance determinants, with tellurite resistance being especially prominent in clade F. Clade E was distinctive in its association with mercury resistance, whereas clade D showed a more selective pattern dominated by cadmium and arsenic resistance. These patterns indicate that WH RIP clades differ in the functional modules that shape plasmid transmission and adaptation.

### Integrated evolutionary landscape of the eight WH RIP clades

To summarize the major features associated with each lineage, we integrated the structural, compositional, taxonomic and ecological characteristics of the eight WH RIP clades (Figure 5). This comparison shows that the major split between the sWH and dWH superclades is accompanied by broad differences in domain organization and plasmid architecture, whereas individual clades are further distinguished by characteristic combinations of plasmid size, RIP length, GC composition, biochemical properties, host distribution, environmental association and accessory functional traits. Notably, clades G and H, as well as clades B, C and D, form well-supported higher-level assemblages, with UFBoot support values of 99 and 92, respectively, suggesting that these sets of clades represent coherent evolutionary subdivisions within the WH RIP phylogeny. Together, these patterns support the view that the eight WH RIP clades represent distinct evolutionary trajectories of plasmid replication systems. Extending this replication-centred framework to additional initiator classes, including AEP-type and rolling-circle replication proteins, should further broaden the evolutionary map of plasmids.

**Figure 5.**
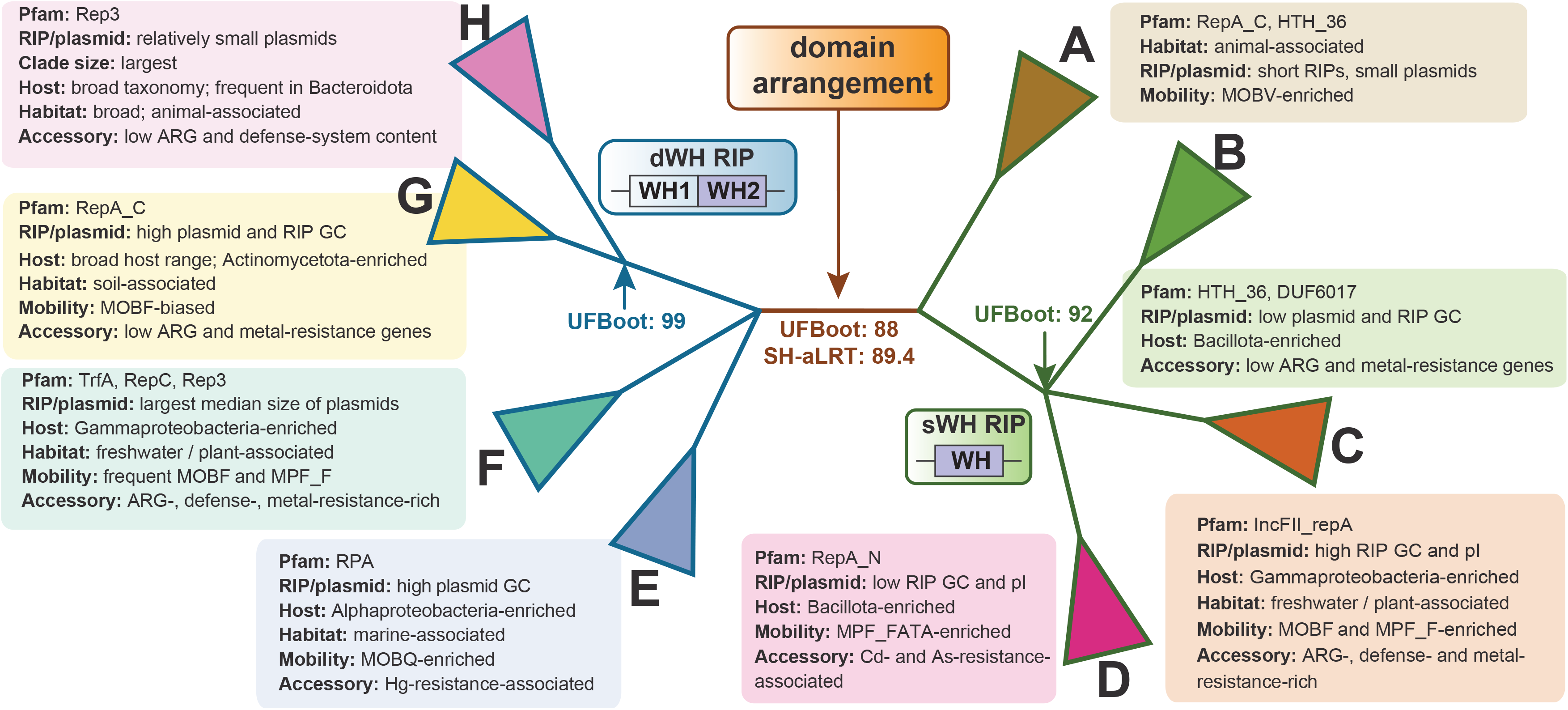
Integrated overview of the eight WH RIP clades. The tree is partitioned into the sWH and dWH superclades, corresponding to RIPs with single– and double-WH domain arrangements, respectively; these contrasting arrangements are illustrated schematically. For each clade, the figure summarizes representative Pfam domains, characteristic plasmid and RIP features, relative clade size where notable, compositional properties, predominant host taxonomic associations, environmental distributions, mobility-related features, and accessory functional traits. The figure integrates the phylogenetic relationships shown in Figure 2 with the dominant compositional, taxonomic, ecological, and functional patterns observed in Figure 4. Descriptors such as “enriched,” “rich,” “biased,” “frequent,” and “associated” summarize the predominant tendencies within each clade rather than fixed categorical thresholds. “Animal-associated” refers to the human-, other mammal-, and insect-associated environmental categories shown in Figure 4B. UFBoot support values are shown for the node uniting clades G and H and for the node uniting clades B, C, and D, whereas both UFBoot and SH-aLRT support values are shown for the split between the sWH and dWH superclades. All branch support values were derived from the WH RIP phylogeny shown in Figure 2.

### Resources for WH RIP identification and plasmid classification

We provide a software package for identification and classification of WH RIPs based on the protein homology relationships defined in this study (https://github.com/yosuken/WHTreeMapper). In addition, although the analyses described above relied on amino acid-level detection of WH domains, practical application to sequence-based identification requires extending detection to the nucleotide level. To this end, total of 48,743 nucleotide sequences corresponding to the conserved WH region of RIP genes (https://doi.org/10.6084/m9.figshare.31969347), including clades A to H and their subclades, were extracted from PLSDB plasmids and curated into the WHRepDB database after redundancy removal (Table 2, n = 8,983, https://doi.org/10.6084/m9.figshare.31883887). To facilitate community use of these resources, we integrated PseudomonasRepDB and WHRepDB into DFAST (DDBJ Fast Annotation and Submission Tool, https://dfast.ddbj.nig.ac.jp/)^62^. Through the advanced annotation options in DFAST, users can screen sequences against additional reference datasets for antimicrobial resistance genes, virulence factors, mobile genetic elements, and plasmid replicons. The available plasmid-typing resources include PseudomonasRepDB, WHRepDB, and AcinetobacterPlasmidTyping. This implementation enables users to apply the PInc and WH RIP-based classification frameworks directly within a widely accessible genome annotation and submission workflow. The command-line version is also available for local use from the GitHub repository at https://github.com/nigyta/dfast_core. Together, these resources enable high-resolution detection and typing of plasmids, including from complex metagenomic data, and provide an accessible platform for characterizing plasmid diversity, host range, and transmission across diverse environments.

To compare WHRepDB with existing replicon typing systems, we analysed PlasmidFinder v2.1, WHRepDB v1.1, and PseudomonasRepDB v1.1 assignments for 59,895 PLSDB plasmids. WHRepDB and/or PseudomonasRepDB detected 34,387 plasmids, including 10,963 without PlasmidFinder assignments (Figure S10). Comparisons of PlasmidFinder types with WHRepDB subclades revealed diverse co-occurrence and assignment patterns (Figures. S11 and S12). Direct reference-sequence comparisons identified WHRepDB matches for 193/488 PlasmidFinder, 457/555 AcinetobacterPlasmidTyping, and 345/404 PseudomonasRepDB entries (Figure S13; https://doi.org/10.6084/m9.figshare.35124934). None of the 35 PInc-6 reference sequences matched WHRepDB, consistent with their replication initiators belonging to the AEP rather than the WH superfamily. These results indicate that WHRepDB complements existing replicon typing systems, although plasmid-level co-occurrence does not necessarily indicate correspondence between individual replication loci.

## Conclusion

In this study, we establish PInc, a curated and experimentally grounded replicon classification framework anchored in historically defined incompatibility groups of *Pseudomonas* plasmids. By coupling experimental identification of replication initiation proteins (RIPs) with sequence-based classification, PInc expands the repertoire of identifiable plasmid replicons and provides a framework for classifying plasmids through their replication systems.

Homology searches beyond PInc plasmids revealed that most replication initiators analysed here share a conserved winged-helix (WH) domain, defining a WH RIP superfamily that unifies a broad range of plasmid replication systems. Remote homology analysis further resolved this superfamily into single– and double-winged-helix superclades and uncovered homology between the WH domain of single-WH RIPs and the WH2 domain of double-WH RIPs, suggesting a shared origin-recognition module across diverse plasmids. By contrast, PInc-6 RIPs were assigned to the AEP superfamily, showing that plasmids grouped historically by incompatibility can encode fundamentally distinct classes of replication initiators.

Using the conserved WH region, we reconstructed a large-scale phylogeny that linked WH RIPs to over 100,000 plasmids across public databases. This phylogeny resolved eight major clades, revealed clade-specific host and environmental distributions, and captured substantial RIP diversity not represented in current typing tools or annotation schemes. It also placed several lineages that are poorly covered by existing plasmid typing systems, including Bacteroidota, Campylobacterota, Thermodesulfobacterota and Fusobacteriota, within a unified WH RIP phylogeny, while bringing together plasmids from cultivated isolates and metagenomic datasets within a single evolutionary framework. More broadly, our results suggest that RIP-based phylogeny can provide the long-missing evolutionary backbone for higher-order plasmid classification and begin to do for plasmids what hallmark gene-based phylogenies have enabled in virology. At present, this framework is limited to plasmids carrying WH RIPs. Expanding comparable phylogenetic frameworks across additional initiator classes will be an important next step towards a more comprehensive evolutionary map of bacterial mobile DNA.

## Methods

### Bacterial strains, plasmids, and culture conditions

The bacterial strains and plasmids used are listed in Table S7. *E. coli* and *Pseudomonas* strains were cultured in Luria broth (LB) or M9 broth^63^ at 30°C (*P. fluorescens* SBW25^64^ and *P*. *putida* KT2440^65,66^) or 37°C (*P. aeruginosa* PAO1^67^ and its derivatives). LB plates containing 1.5% agar were used for filter mating. Ampicillin (Ap, 50 μg/mL), piperacillin (Pip, 16 μg/mL), kanamycin (Km, 12.5 or 25 μg/mL), rifampicin (Rif, 50 μg/mL), streptomycin (Sm 150 μg/mL), and tetracycline (Tc, 50 μg/mL for the others) were added to the medium.

### DNA manipulation

Total DNA from the bacterial strains was extracted using a NucleoSpin Tissue Kit (TAKARA BIO, Japan). Polymerase chain reaction (PCR) was performed on a T100 thermal cycler (Bio-Rad Laboratories, USA) using the KOD One PCR Master Mix (TOYOBO, Japan). The primers used in this study are listed in Table S8. The following amplification conditions were used: 30 cycles at 98°C for 10 s, 67°C for 5 s, and 68°C for 1 s. The pUC-GW plasmid with DNA fragments including putative RIP genes and *oriV*s of each plasmid, except those for RP1-1 and pMG26, was synthesised by the custom service of Azenta Life Science, USA. Restriction enzymes (New England Biolabs, USA or TAKARA BIO), the HiYieldTM Gel/PCR DNA fragments Extraction kit (RBC Bioscience, Taiwan), NEBuilder Hifi DNA Assembly system (New England Biolabs), and competent *E. coli* JM109 and DH5α cells (RBC Bioscience) were used for cloning the DNA fragments. Plasmid DNA from the bacterial strains was extracted using the NucleoSpin Plasmid EasyPure Kit (TAKARA BIO). Electroporation to introduce the plasmids into different bacterial strains was performed using a MicroPulser electroporator (Bio-Rad Laboratories). All other procedures were performed according to standard methods^63^.

### Complete sequencing of the plasmids by next-generation sequencing

The nucleotide sequences of the plasmid DNAs were determined as follows: whole genomic DNA extracted from their host (*P. aeruginosa* PAO1808, a derivative strain of PAO1^68^) was sequenced using the HiSeq2500 platform (Illumina, USA). Trimmed high-quality short reads (read length >140 bp and quality score >15) were assembled using SPAdes v3.14.0 using the plasmid option^69^. If circular contig(s) could not be found, the host chromosomal DNA was removed by mapping the resultant contigs on the host genome sequences [their deposited sequences, that is, NCBI RefSeq accession number NC_002516 (*P. aeruginosa* PAO1)] using Geneious Prime software v2020.01^70^. Subsequently, the plasmid reads were extracted from the remaining contigs using SAMtools v1.7^71^ and SeqKit v0.8.0^72^ and reassembled using SPAdes^73^. For RP1-1 and pMG26, gaps in plasmids were closed by PCR with the designed primer sets based on the ends of contigs and Sanger sequencing of the PCR products.

### Gene annotation of plasmids

Gene annotations of the plasmids were performed using DFAST-core v1.2.5^62^ with an in-house database of plasmid sequences collected from NCBI RefSeq and then corrected manually. Synteny analyses of plasmids were performed using Easyfig v2.2.2^74^. Other *in silico* analyses were performed using Geneious Prime v2020.01. The annotated plasmid sequences were deposited under the following GenBank accession numbers: LC685025, LC685026, LC685027, LC685593, and LC700336.

### Detection and phenotype identification of ARGs

ARG sequences were detected using Staramr v0.10.0 and ResFinder database v2.2.1 updated on 2023-10-27 (https://bitbucket.org/genomicepidemiology/resfinder_db)^75,76^ with the following criteria: ≥ 90% identity and ≥ 60% coverage. Clinically important ARGs, including genes conferring resistance to carbapenem, colistin, or tigecycline, in Gram-negative bacteria were further detected based on the phenotype information of ARGs in AMRFinderPlus v3.11.26 updated on 2023-11-16 (https://www.ncbi.nlm.nih.gov/pathogens/antimicrobial-resistance/AMRFinder/)^77^.

### Prediction of RIP genes and *oriV*s of each plasmid

To identify the genes encoding the RIP and origin of vegetative replication (*oriV*) regions of Rms139, Rms163, RP1-1, R716, pQBR103, pPT23-C1_1, two prediction approaches were used; after coding sequence (CDS) annotations using DFAST-core, cds.fna and protein.faa files generated by DFAST-core were used for the following: (i) nucleotide sequence-based searches were performed using BLASTn; (ii) amino acid sequence-based protein structure prediction and the structure-based searches were performed as described later. The GC-skew value, which has been commonly used to measure the asymmetry in the composition of two nucleotides, guanine (G) and cytosine (C), along a DNA strand and is defined as (G-C)/(G+C), of the plasmids was used by Webskew (https://genskew.csb.univie.ac.at/webskew) to detect the changes in the GC skew, which indicates the DNA replication origins or termini^78,79^. Subsequently, the nucleotide or amino acid sequences of putative CDSs around the above GC-skew shift point(s) were subjected to methods (i) or (ii). The putative promoter region of the RIP gene was predicted using BPROM (http://www.softberry.com/berry.phtml?topic=bprom&group=help&subgroup=gfindb)^80^. Subsequently, the predicted RIP and *oriV* sequences of each PInc groups were cloned into the pUC19^81^-derivative or pUC-GW-Amp vector (Azenta Life Science). Transformation of *P. aeruginosa* PAO1808, *P. putida* KT2440 or *P. fluorescens* SBW25 was performed with each of the resultant plasmids (at most 472 ng/μL), and the cultures were spread onto LB+Pi, or LB+Km plates. After incubation at 30 or 37°C overnight, colonies on the plates were isolated. Next, the presence of the mini-plasmid was confirmed by PCR with specific primers after extraction of total DNA of transformants. Further details of the mini-replicon assays, including construct design, transformation conditions, and experimental outcomes, are provided in Supplemental Text S1.

### Protein structure prediction of RIPs

AlphaFold3-mediated protein structure prediction was performed using the amino acid sequences of the RIP gene products in representative *Pseudomonas* plasmids (https://doi.org/10.6084/m9.figshare.26778898) on the AlphaFold server^48^. A structure-based protein homology search using the predicted structural models was performed for the PDB100 database on the FoldSeek server in 3Di/AA mode^82^. The predicted and experimental structures were visualised and compared using PyMOL v3.1.3.1. The predicted structures were coloured based on the root mean square deviation (RMSD) from the reference experimental structures, i.e., the Pi or Rep19A^50,83^ proteins, which are RIPs of the R6K and pCA-347 plasmids, respectively, using the ColorByRMSD script (https://pymolwiki.org/index.php/ColorByRMSD). The distances between aligned C-alpha atom pairs were stored as B-factors of these residues, which are coloured using a colour spectrum, with blue specifying the minimum pairwise RMSD and red indicating the maximum. Unaligned residues were coloured gray.

### Identification, retrieval, and re-annotation of plasmids from public databases

Complete plasmid sequences were downloaded in FASTA format using the Entrez Programming Utilities (E-utilities; https://www.ncbi.nlm.nih.gov/books/NBK25501/) from a public database (PLSDB 2023_11_23_v2) (https://doi.org/10.6084/m9.figshare.26768173)^7,84^. To classify the plasmids in the PLSDB based on their rep sequence similarity, the RIP gene sequences were detected using Staramr v0.10.0 (https://github.com/phac-nml/staramr) and rep databases of MOB-suite v3.1.8 updated on 2023-12-12 (https://github.com/phac-nml/mob-suite) (https://doi.org/10.6084/m9.figshare.26768170)^85^, PlasmidFinder v2.1 updated on 2023-12-04 (https://bitbucket.org/genomicepidemiology/plasmidfinder_db) (https://doi.org/10.6084/m9.figshare.26778199)^86^, our custom-made *Pseudomonas* plasmid RIP gene database (termed repP and PseudomonasRepDB database, https://doi.org/10.6084/m9.figshare.31883911 and https://doi.org/10.6084/m9.figshare.26778175), and our custom-made plasmid RIP WH database (termed WHRepDB database, https://doi.org/10.6084/m9.figshare.31883887) with the following criteria: ≥90% identity and ≥60% coverage (Table 2). The Tus protein, known for binding to DNA replication terminus (*ter*) sites of replicons, encoded by rep_cluster_1115 in the MOB-suite database, is not included in our custom-made RIP reference sequences. The repP database was constructed using experimentally curated RIP genes from the 87 reference *Pseudomonas* plasmids shown in Table S1.

Furthermore, duplicated RIP sequences between the MOB-suite and repP databases (Table S6) and incorrect RIP sequences, including uncertain IncP and rep_cluster_1115, were manually removed from the MOB-suite database. The modified MOB-suite database (https://doi.org/10.6084/m9.figshare.26768167) was used to detect multi-replicon *Pseudomonas* plasmids, which are single plasmids that can belong to more than one group other than the PInc groups. Among the candidate PInc-1 (IncP-1) plasmids identified based on RIP sequence similarity, some records represented laboratory-constructed plasmids or plasmid derivatives rather than naturally occurring plasmids. Because their artificial origin was not always evident from the sequence-record metadata, we reviewed the corresponding literature and identified six such plasmids (PD584_L3, PD689_delTT1, pJM105, pJM50, pRK2013, and pUZ8002) ^87^. These sequences were excluded from the final PInc-1 dataset. The raw data of matches between reference database entries (Figure S13) was deposited in https://doi.org/10.6084/m9.figshare.35124934.

For the final dataset, the sequence length (bp) and GC content (%, %GC) were calculated using SeqKit v2.3.0 (https://bioinf.shenwei.me/seqkit/)^72^. Plasmid sequences belonging to each PInc groups were re-annotated using DFAST-core, with a reference genome from each group to ensure consistency in the functional annotations.

### Comparative genomics of plasmids

Comparative genomics of all the plasmids of the PInc groups were performed using PIRATE^88^ (https://github.com/SionBayliss/PIRATE). Here, we used DIAMOND^89^ v0.9.14 rather than BLAST^90^ to identify homologous genes (protein families) in PIRATE. PIRATE was run on default parameters with a high-scoring segment pair (HSP) query length threshold of 0.5, a sequence similarity step range of 10, 20, 30, 40, 50, 60, 70, 80, 90, 95, and 98%, and an inflation value of 2 in MCL (v14.137), a Markov cluster algorithm (https://micans.org/mcl/). The core genes in each PInc group were those that were present in more than 95% of the plasmids included in the pangenome analysis. Statistical computing and result visualisation were performed using R v4.1.3 (https://www.r-project.org/). The output of SeqKit and PIRATE are available in Figshare (https://doi.org/10.6084/m9.figshare.26789416).

### Homolog collection and phylogenetic analysis of RIPs

Detailed methods are provided in Supplemental Text S1 and illustrated in Figure S4. In brief, to analyse the evolutionary relationships of RIPs, we identified 2,374 RIPs that originated from 2,332 PInc plasmids. Subsequently, 251 non-redundant proteins at 100% identity were extracted using MMseqs^91^ cluster v15-6f452. Sequence homology between the non-redundant proteins was assessed using BLASTp^90^, and the resultant hits were visualised via CPAP v0.3.0 (https://github.com/yosuken/CPAP). To detect remote homology between PInc groups and collect comprehensive RIP sequences, we extended the sequence diversity through a multistep homology search process. First, we searched ORFs of PLSDB plasmids predicted using orfM^92^ (n = 20,331,985) for homologous RIPs using hmmsearch in HMMER^93^ v3.3.2. In addition, 8 additional RIPs sequences that are distant from RIPs were manually picked up and homology searches using diamond blastp against the ORFs of PLSDB plasmids were performed. This process generated 19 homologous groups of RIPs. Second, UniParc proteins (downloaded in Feb 2024; n = 607,912,929) were searched with hmmsearch using 19 HMMs generated from the homologous groups. We merged the UniParc sequences into 19 homologous groups, and filtered out potential contamination of other functional proteins, including various transcriptional factors, using hmmsearch against the Pfam database. After merging the 19 homologous groups into 14 based on the UniParc sequence overlapping, we selected 5,488 representative sequences for the 14 homologous groups. To investigate whether these groups exhibit homology with each other, a sensitive hidden Markov model HMM–HMM comparison was performed using HHblits^94^ v3.3.0. We extracted the homologous regions of the alignments of the 13 groups (containing 4,793 sequences in total). These regions were aligned using MAFFT^95^ v7.455.

After removing gappy sequences in the alignment, a maximum-likelihood phylogeny of WH RIPs was reconstructed from the resulting 150-position alignment using IQ-TREE^96^ v2.2.2.6. We used the parameters “––perturb 0.2 ––nstop 500”, following the software manual recommendation for datasets containing many short sequences. The best fit model (Q.pfam+F+R10) was selected using ModelFinder^97^. Branch support was assessed with UFBoot2^98^ and SH-aLRT^99^ in IQ-TREE. We discarded the abnormally long branches using TreeShrink^100^. To evaluate the root of the WH RIP phylogeny independently of domain-based interpretation, we applied minimal ancestor deviation (MAD) rooting to the unrooted maximum-likelihood tree using MAD^59^ v2.2 with default settings. The MAD-inferred root was then compared with the major split separating the sWH and dWH superclades. The final tree was rooted at the sWH-dWH split using gotree^101^ v0.4.0.

### Recruitment of associated plasmids onto the WH RIP phylogeny

The tree-associated plasmids from PLSDB^7^ (version 2023_11_23_v2) and IMG/PR^6^ (version 2023-08-08_1) were identified as follows. Open reading frames (ORFs) were predicted using Prodigal^102^. Following the developer’s recommendations, plasmids longer than 100 kb were processed in the default mode, whereas plasmids of 100 kb or shorter were processed in metagenomic mode “––meta”. The predicted ORFs were then searched against HMM profiles representing the 42 WH RIP subclades using PiPP v0.3.0 (https://github.com/yosuken/PiPP), which runs hmmsearch in HMMER v3.3.2^93^, with the options “––evalue 1e-5 ––evaluedom 1e-2 ––minhmmcovdom 0.2 –minhmmcov 0.8 ––only-detect”. For each best hit, the region of the ORF covered by the HMM match was extracted. These extracted regions were then assigned to the aligned representative WH RIP sequences used for phylogenetic reconstruction (n = 4,769) using diamond^89^ blastp (v2.1.9) with the options “––ultra-sensitive ––id 40 ––subject-cover 80 ––max-target-seqs 1 ––dbsize 1e+9 ––evalue 1e-5”. The extracted regions mapped to the tree were then organized at both the ORF and plasmid levels. For plasmid-level analyses, mapped WH RIP-containing ORFs were grouped according to their source plasmids, and plasmids carrying two or more such ORFs were classified as multi-replicons. For summaries at the clade, subclade, or individual RIP level, plasmids were deduplicated within each category, so that multiple ORFs from the same plasmid assigned to the same category were counted once. The number of WH replicons per plasmid, however, was defined by the total number of mapped WH RIP-containing ORFs detected in that plasmid.

### Plasmid metadata and functional annotation associated with the WH RIP phylogeny

Taxonomic metadata for tree-associated plasmids were obtained from the source databases: NCBI taxonomy for PLSDB and GTDB taxonomy for IMG/PR. Taxonomic distributions were summarized using the manually curated higher-rank host categories shown in Figure 4A, with class-level categories used for the phylum Pseudomonadota and phylum-level categories used for several other bacterial lineages. Because nomenclature differs between the two databases, original names were retained for each source but interpreted as corresponding higher-level lineages when appropriate (e.g., Bacillota in PLSDB versus Firmicutes in IMG/PR, Actinomycetota versus Actinobacteriota, and Thermodesulfobacteriota versus Desulfobacterota). Low-frequency taxa were grouped into an “others” category for visualization.

Environmental analyses were performed using IMG/PR-linked plasmids only, because environmental metadata was available for these plasmids. Habitat categories shown in Figure 4B were defined based on the original metadata labels, including host-associated environments, aquatic environments (marine, freshwater, and non-marine saline and alkaline), terrestrial soil, engineered environments (bioreactor, built environment, and wastewater/WWTP), and “others”. Records annotated as “Engineered; Modeled; Simulated communities” were treated as lacking environmental information and excluded from this analysis, because they do not specify a source habitat.

Replicon typing was performed using PlasmidFinder^9^ v2.1 updated on 2023-12-04, AcinetobacterPlasmidTyping with database version feb2025_v3, and PseudomonasRepDB for PInc assignment. For all three databases, hits were identified using PlasmidFinder with a minimum nucleotide identity threshold of 90% “––threshold 90” and a minimum coverage of 60% “––mincov 60”. Additional replicon typing was performed using MOB-typer in MOB-suite v3.1.8 with the equivalent options “–– min_rep_ident 90 ––min_rep_cov 60”. Pfam annotations of tree-associated RIPs were identified using hmmsearch in HMMER v3.3.2 against the Pfam v38.1 database and parsed using parse_hmmsearch.rb (https://github.com/yosuken/parse_hmmsearch) with the thresholds “––gene-evalue 1e-5 ––evalue 1e-2 ––min-ali-cov-dom 0.2 ––min-ali-cov 0.5”. For visualization of the annotation rings in Figure 2A, taxonomic, environmental, and replicon-typing information associated with mapped plasmids was summarized as stacked-bar compositions for each tree-linked leaf. The WH RIP phylogeny and its annotation rings in Figure 2A were visualized using iTOL v7^103^.

MOB and MPF classes were identified using MacSyFinder^104^ v2.1.4 with the CONJScan/Plasmids models^105^ under default settings. For MOB classification, hits with gene_name annotated as T4SS_MOB(X) and model_fqn annotated as CONJScan/Plasmids/MOB were used. For MPF classification, hits with model_fqn annotated as CONJScan/Plasmids/T4SS_type(X) were used. For Figures 4F and 4G, MOB and MPF annotations were summarized at the plasmid level for clade-associated plasmids. Plasmids carrying two or more MOB classes or MPF classes were assigned to the category “multiple”. Plasmids without detectable MOB or MPF annotations were retained in the calculations of clade-wise proportions but were not displayed as separate categories. Antimicrobial resistance (AMR) genes were identified using AMRFinderPlus v3.12.8 with database version 2024-07-22.1 under default settings. Hits annotated with Element subtype = AMR were retained. Metal resistance genes were identified in the same manner as AMR genes, except that hits annotated with Element subtype = METAL were retained. For Figure 4J, metal resistance was summarized at the plasmid level for clade-associated plasmids. The number of resistant metals per plasmid was defined as the number of distinct metal resistance classes detected in each plasmid, irrespective of the number of genes assigned to each class. Defense systems were identified using DefenseFinder^106^ v1.3.0 with default settings. For Figure 4I, defense systems were summarized at the plasmid level for clade-associated plasmids, and the number of defense systems per plasmid was defined on a system basis. Anti-defense systems were excluded from these counts. Plasmids were grouped into the bins shown in Figure 4I.

For Figures 4C–E, analyses were performed at the level of clade-associated RIPs. Plasmid length was defined as the total nucleotide length of the source plasmid, and RIP length as the amino-acid length of the corresponding RIP. RIP GC content was calculated from the nucleotide sequence of the RIP-encoding CDS, whereas plasmid GC content was calculated from the full plasmid nucleotide sequence. The RIP–plasmid GC difference was defined as RIP %GC minus plasmid %GC. The isoelectric point (pI) of RIPs and plasmid-encoded proteins was calculated using pepstats in EMBOSS v6.6.0 with default settings. Plasmid proteome pI was defined as the median pI of all predicted proteins encoded by each plasmid, and the RIP–plasmid proteome pI difference was defined as RIP pI minus plasmid proteome pI. Unless otherwise noted, analyses were performed either at the plasmid level or at the RIP level, depending on the panel. Figures 2C–E and 4F–J summarize plasmid-level properties, whereas Figure 2F and Figures 4C–E summarize RIP-level properties. Figures 2A and 2B are based on the representative WH RIP sequences used for phylogenetic reconstruction.

Data visualization for Figures 2 and 4 was performed in R v4.1.2 using the ggplot2 package, except that phylogenetic tree visualization was carried out using iTOL. Figures 2, 4, and 5 were subsequently assembled and finalized in Adobe Illustrator based on the resulting plots. The phylogenetic tree, multiple sequence alignment, associated plasmid metadata, and iTOL visualization files are available in Figshare (https://doi.org/10.6084/m9.figshare.28837559).

## Data Availability

All datasets generated and analyzed during this study have been deposited in publicly accessible repositories and are freely available to reviewers as follows:

Replicon reference database (repP database): Experimentally supported replication initiation protein (RIP) sequences from representative *Pseudomonas* plasmids used for PInc classification are available at Figshare: https://doi.org/10.6084/m9.figshare.31883911 (nucleotides), https://doi.org/10.6084/m9.figshare.26778898 (amino acids).

*Pseudomonas* plasmid dataset with PInc classification (PseudomonasRepDB): Curated plasmid dataset derived from PLSDB, including PInc classification results and associated annotations: https://doi.org/10.6084/m9.figshare.26778175.

WH RIP database (WHRepDB): Database of winged-helix (WH) domain-containing replication initiation proteins used for large-scale homology search and classification: https://doi.org/10.6084/m9.figshare.31883887.

WH-domain sequence dataset: A total of 48,743 WH-domain sequences identified from

RIP genes and used for downstream phylogenetic and structural analyses: https://doi.org/10.6084/m9.figshare.31969347.

Phylogenetic datasets and tree files: Multiple sequence alignment, maximum-likelihood phylogenetic tree, associated plasmid metadata, and iTOL visualization files: https://doi.org/10.6084/m9.figshare.28837559.

PLSDB-derived plasmid dataset (version 2023_11_23_v2): https://doi.org/10.6084/m9.figshare.26768173.

Original and modified MOB-suite database used for replicon detection: https://doi.org/10.6084/m9.figshare.26768170 (original), https://doi.org/10.6084/m9.figshare.26768167 (modified) PlasmidFinder database used in this study: https://doi.org/10.6084/m9.figshare.26778199.

The raw data of matches between reference database entries (Figure S13) https://doi.org/10.6084/m9.figshare.35124934

Additional analysis outputs (pangenome analysis, sequence statistics, etc.): https://doi.org/10.6084/m9.figshare.26789416.

The annotated plasmid sequences have been deposited in the GenBank/EMBL/DDBJ under accession numbers LC685025, LC685026, LC685027, LC685593, and LC700336.

## Code Availability

We provide WHTreeMapper, a software package for identifying and classifying WH RIPs based on the protein homology relationships established in this study (https://github.com/yosuken/WHTreeMapper). Sequence homology among the non-redundant proteins was assessed and visualised using CPAP v0.3.0 (https://github.com/yosuken/CPAP). Regions corresponding to hmmsearch hits were extracted using PiPP v0.3.0 (https://github.com/yosuken/PiPP).

## Supplementary Data Statement

Supplementary Data (Supplemental Text S1, Supplemental Figures: Figs. S1-S17, and Supplemental Tables S1-S9) are available online.

## Acknowledgements

We are grateful to Prof. S. Iyobe of Gunma University School of Medicine and Prof. M. Tsuda of Tohoku University for providing Rms139, Rms163, RP1-1, R716, and pMG26. We also thank Dr. Steve Petrovski of the La Trobe University for providing the sequence of R91-5, and Dr. Michael Brockhurst and Dr. Christian Harrison of the University of Manchester for providing *Pseudomonas fluorescens* SBW25 and plasmid pQBR103. Computation time was provided by the Super Computer System, Institute for Chemical Research, Kyoto University.

## Declaration of generative AI and AI-assisted technologies in the manuscript preparation process

During the preparation of this work, the authors used ChatGPT (OpenAI) for language editing and improving the clarity of English expression. The authors also used GitHub Copilot (GitHub/Microsoft) and Claude Code (Anthropic) to assist in the development of parts of the code. All generated code and final outputs were reviewed and verified by the authors, who take full responsibility for them. These tools were not used to generate scientific content or interpret data. All content was critically reviewed and edited by the authors, who take full responsibility for the final manuscript.

## Contributions

YN, HS, MSu, MSh conceived, designed, and supervised the study. YN, KKa, TK, NI, YM, MTs, TM, HX, YTa, Yts, AH, RM, HD, HS, MSu, MSh performed experiments and data analysis. YN, RM, HD, KKi, CS-M, HN, HS, MSu, MSh wrote, reviewed, and edited the manuscript. All authors contributed to manuscript revision, read and approved the submitted version.

## References

1. Wiedenbeck, J. & Cohan, F. M. Origins of bacterial diversity through horizontal genetic transfer and adaptation to new ecological niches. FEMS Microbiol. Rev. 35, 957–976 (2011).

2. Haudiquet, M., de Sousa, J. M., Touchon, M. & Rocha, E. P. C. Selfish, promiscuous and sometimes useful: how mobile genetic elements drive horizontal gene transfer in microbial populations. Philos. Trans. R. Soc. Lond. B Biol. Sci. 377, 20210234 (2022).

3. Helinski, D. R. A brief history of plasmids. EcoSal Plus 10, eESP00282021 (2022).

4. Starling, S. Plasmid evolution in the time of antibiotics. Nat. Rev. Microbiol. 24, 95 (2026).

5. Cazares, A. et al. Pre– and postantibiotic epoch: the historical spread of antimicrobial resistance. Science 390, eadr1522 (2025).

6. Camargo, A. P. et al. IMG/PR: a database of plasmids from genomes and metagenomes with rich annotations and metadata. Nucleic Acids Res. 52, D164– D173 (2024).

7. Schmartz, G. P. et al. PLSDB: advancing a comprehensive database of bacterial plasmids. Nucleic Acids Res. 50, D273–D278 (2022).

8. Garcillán-Barcia, M. P., Redondo-Salvo, S. & de la Cruz, F. Plasmid classifications. Plasmid 126, 102684 (2023).

9. Carattoli, A. et al. In silico detection and typing of plasmids using PlasmidFinder and plasmid multilocus sequence typing. Antimicrob. Agents Chemother. 58, 3895–3903 (2014).

10. Fernández-López, R., Redondo, S., Garcillán-Barcia, M. P. & de la Cruz, F. Towards a taxonomy of conjugative plasmids. Curr. Opin. Microbiol. 38, 106–113 (2017).

11. Redondo-Salvo, S. et al. Pathways for horizontal gene transfer in bacteria revealed by a global map of their plasmids. Nat. Commun. 11, 3602 (2020).

12. Garcillán-Barcia, M. P., Redondo-Salvo, S., Vielva, L. & de la Cruz, F. MOBscan: automated annotation of MOB relaxases. Methods Mol. Biol. 2075, 295–308 (2020).

13. Coluzzi, C., Garcillán-Barcia, M. P., de la Cruz, F. & Rocha, E. P. C. Evolution of plasmid mobility: origin and fate of conjugative and nonconjugative plasmids. Mol. Biol. Evol. 39, msac115 (2022).

14. Garcillán-Barcia, M. P., Francia, M. V. & de la Cruz, F. The diversity of conjugative relaxases and its application in plasmid classification. FEMS Microbiol. Rev. 33, 657–687 (2009).

15. Koonin, E. V., Kuhn, J. H., Dolja, V. V. & Krupovic, M. Megataxonomy and global ecology of the virosphere. ISME J. 18, wrad042 (2024).

16. International Committee on Taxonomy of Viruses Executive Committee. The new scope of virus taxonomy: partitioning the virosphere into 15 hierarchical ranks. Nat. Microbiol. 5, 668–674 (2020).

17. Moradali, M. F., Ghods, S. & Rehm, B. H. A. *Pseudomonas aeruginosa* lifestyle: a paradigm for adaptation, survival, and persistence. Front. Cell. Infect. Microbiol. 7, 39 (2017).

18. Thomas, C. M. & Haines, A. S. Plasmids of the genus Pseudomonas. In Pseudomonas: Volume 1 Genomics, Life Style and Molecular Architecture (ed. Ramos, J.-L.) 197–231 (Kluwer Academic/Plenum Publishers, New York, 2004).

19. Shintani, M. & Suzuki, H. Plasmids and their hosts. In DNA Traffic in the Environment (eds Nishida, H. & Oshima, T.) 109–133 (Springer, Singapore, 2019).

20. Sawada, Y., Yaginuma, S., Tai, M., Iyobe, S. & Mitsuhashi, S. Plasmid-mediated penicillin beta-lactamases in *Pseudomonas aeruginosa*. Antimicrob. Agents Chemother. 9, 55–60 (1976).

21. Sagai, H. et al. Classification of R plasmids by incompatibility in *Pseudomonas aeruginosa*. Antimicrob. Agents Chemother. 10, 573–578 (1976).

22. Summers, A. O. & Jacoby, G. A. Plasmid-determined resistance to boron and chromium compounds in *Pseudomonas aeruginosa*. Antimicrob. Agents Chemother. 13, 637–640 (1978).

23. Kawakami, Y., Mikoshiba, F., Nagasaki, S., Matsumoto, H. & Tazaki, T. Prevalence of *Pseudomonas aeruginosa* strains possessing R factor in a hospital. J. Antibiot. 25, 607–609 (1972).

24. Hedges, R. W. & Jacob, A. E. Mobilization of plasmid-borne drug resistance determinants for transfer from *Pseudomonas aeruginosa* to *Escherichia coli*. Mol. Gen. Genet. 140, 69–79 (1975).

25. Ingram, L., Sykes, R. B., Grinsted, J., Saunders, J. R. & Richmond, M. H. A transmissible resistance element from a strain of *Pseudomonas aeruginosa* containing no detectable extrachromosomal DNA. J. Gen. Microbiol. 72, 269–279 (1972).

26. Shintani, M., Suzuki, H., Nojiri, H. & Suzuki, M. Reconsideration of the previously classified incompatibility groups of plasmids, IncP-1 and IncP-11. Environ. Microbiol. 25, 1071–1076 (2023).

27. Bryan, L. E., Semaka, S. D., van den Elzen, H. M., Kinnear, J. E. & Whitehouse, R. L. Characteristics of R931 and other *Pseudomonas aeruginosa* R factors. Antimicrob. Agents Chemother. 3, 625–637 (1973).

28. Jacoby, G. A. Plasmid determined resistance to carbenicillin and gentamicin in Pseudomonas aeruginosa. In Plasmids and Transposons: Environmental Effects and Maintenance Mechanisms (eds Stuttard, C. & Rozee, K. R.) 83–96 (Academic Press, New York, 1980).

29. Bradley, D. E. Specification of the conjugative pili and surface mating systems of *Pseudomonas* plasmids. J. Gen. Microbiol. 129, 2545–2556 (1983).

30. Cain, D. & Holloway, B. W. Prime plasmids derived from the IncP-10 plasmid R91-5 in *Pseudomonas putida*. FEMS Microbiol. Lett. 24, 97–101 (1984).

31. Davies, S. & Krishnapillai, V. DNA sequence analysis of the replication region of the *Pseudomonas aeruginosa* plasmid R91-5. J. Genet. 69, 101–112 (1990).

32. Jacoby, G. A. Resistance plasmids of Pseudomonas. In The Biology of Pseudomonas (ed. Sokatch, J. R.) 265–293 (Academic Press, Orlando, 1986).

33. Suzuki, M., Suzuki, H., Nishimura, Y., Nojiri, H. & Shintani, M. Enhancing plasmid typing with MOB-typer: resolving IncP and other incompatibility group misclassifications in *Pseudomonas*. Microb. Genom. 11, 001491 (2025).

34. Kawalek, A. et al. Genome sequence of *Pseudomonas aeruginosa* PAO1161, a PAO1 derivative with the ICEPae1161 integrative and conjugative element. BMC Genomics 21, 14 (2020).

35. Boronin, A. M. Diversity of *Pseudomonas* plasmids: to what extent? FEMS Microbiol. Lett. 100, 461–467 (1992).

36. Van der Auwera, G. A. et al. Plasmids captured in *C. metallidurans* CH34: defining the PromA family of broad-host-range plasmids. Antonie Van Leeuwenhoek 96, 193–204 (2009).

37. Law, A. et al. Biosolids as a source of antibiotic resistance plasmids for commensal and pathogenic bacteria. Front. Microbiol. 12, 606409 (2021).

38. Yanagiya, K. et al. Novel self-transmissible and broad-host-range plasmids exogenously captured from anaerobic granules or cow manure. Front. Microbiol. 9, 2602 (2018).

39. Tett, A. et al. Sequence-based analysis of pQBR103; a representative of a unique, transfer-proficient mega plasmid resident in the microbial community of sugar beet. ISME J. 1, 331–340 (2007).

40. Köhler, K. A. K. et al. Complete genome sequence of *Pseudomonas* sp. strain VLB120 a solvent tolerant, styrene degrading bacterium, isolated from forest soil. J. Biotechnol. 168, 729–730 (2013).

41. Chen, F. et al. VIM-encoding IncpSTY plasmids and chromosome-borne integrative and mobilizable elements (IMEs) and integrative and conjugative elements (ICEs) in *Pseudomonas*. Ann. Clin. Microbiol. Antimicrob. 21, 10 (2022).

42. Hayakawa, M. et al. Hitherto-unnoticed self-transmissible plasmids widely distributed among different environments in Japan. Appl. Environ. Microbiol. 88, e01114–22 (2022).

43. Tokuda, M. et al. Transconjugant range of PromA plasmids in microbial communities is predicted by sequence similarity with the bacterial host chromosome. Microb. Genom. 9, 001043 (2023).

44. Lawrence, J. G. & Ochman, H. Amelioration of bacterial genomes: rates of change and exchange. J. Mol. Evol. 44, 383–397 (1997).

45. Rocha, E. P. & Danchin, A. Base composition bias might result from competition for metabolic resources. Trends Genet. 18, 291–294 (2002).

46. Dietel, A.-K., Merker, H., Kaltenpoth, M. & Kost, C. Selective advantages favour high genomic AT-contents in intracellular elements. PLoS Genet. 15, e1007778 (2019).

47. Remmert, M., Biegert, A., Hauser, A. & Söding, J. HHblits: lightning-fast iterative protein sequence searching by HMM-HMM alignment. Nat. Methods 9, 173–175 (2012).

48. Abramson, J. et al. Accurate structure prediction of biomolecular interactions with AlphaFold 3. Nature 630, 493–500 (2024).

49. Wegrzyn, K. et al. Defining a novel domain that provides an essential contribution to site-specific interaction of Rep protein with DNA. Nucleic Acids Res. 49, 3394– 3408 (2021).

50. Schumacher, M. A., et al. Mechanism of staphylococcal multiresistance plasmid replication origin assembly by the RepA protein. Proc. Natl Acad. Sci. USA 111, 9121–9126 (2014).

51. Itou, H., Yagura, M., Shirakihara, Y. & Itoh, T. Structural basis for replication origin unwinding by an initiator primase of plasmid ColE2-P9: duplex DNA unwinding by a single protein. J. Biol. Chem. 290, 3601–3611 (2015).

52. Iyer, L. M., Koonin, E. V., Leipe, D. D. & Aravind, L. Origin and evolution of the archaeo-eukaryotic primase superfamily and related palm-domain proteins: structural insights and new members. Nucleic Acids Res. 33, 3875–3896 (2005).

53. Haines, A. S., Jones, K., Cheung, M. & Thomas, C. M. The IncP-6 plasmid Rms149 consists of a small mobilizable backbone with multiple large insertions. J. Bacteriol. 187, 4728–4738 (2005).

54. Haines, A. S. et al. Plasmids from freshwater environments capable of IncQ retrotransfer are diverse and include pQKH54, a new IncP-1 subgroup archetype. Microbiology 152, 2689–2701 (2006).

55. Aravind, L., Anantharaman, V., Balaji, S., Babu, M. M. & Iyer, L. M. The many faces of the helix-turn-helix domain: transcription regulation and beyond. FEMS Microbiol. Rev. 29, 231–262 (2005).

56. Giraldo, R. & Fernández-Tresguerres, M. E. Twenty years of the pPS10 replicon: insights on the molecular mechanism for the activation of DNA replication in iteron-containing bacterial plasmids. Plasmid 52, 69–83 (2004).

57. Giraldo, R. Common domains in the initiators of DNA replication in *Bacteria*, *Archaea* and *Eukarya*: combined structural, functional and phylogenetic perspectives. FEMS Microbiol. Rev. 26, 533–554 (2003).

58. Díaz-López, T., Dávila-Fajardo, C., Blaesing, F., Lillo, M. P. & Giraldo, R. Early events in the binding of the pPS10 replication protein RepA to single iteron and operator DNA sequences. J. Mol. Biol. 364, 909–920 (2006).

59. Tria, F. D. K., Landan, G. & Dagan, T. Phylogenetic rooting using minimal ancestor deviation. *Nat*. Ecol. Evol. 1, 0193 (2017).

60. Lam, M. M. C. & Hamidian, M. Examining the role of *Acinetobacter baumannii* plasmid types in disseminating antimicrobial resistance. *npj Antimicrob*. Resist. 2, 1 (2024).

61. Komori, H. et al. Crystal structure of a prokaryotic replication initiator protein bound to DNA at 2.6 Å resolution. EMBO J. 18, 4597–4607 (1999).

62. Tanizawa, Y., Fujisawa, T. & Nakamura, Y. DFAST: a flexible prokaryotic genome annotation pipeline for faster genome publication. Bioinformatics 34, 1037–1039 (2018).

63. Sambrook, J. & Russell, D. W. Molecular Cloning: A Laboratory Manual 3rd edn (Cold Spring Harbor Laboratory Press, Cold Spring Harbor, NY, 2001).

64. Theodosiou, L., Farr, A. D. & Rainey, P. B. Barcoding populations of *Pseudomonas fluorescens* SBW25. J. Mol. Evol. 91, 254–262 (2023).

65. Bagdasarian, M. et al. Specific-purpose plasmid cloning vectors. II. Broad host range, high copy number, RSF1010-derived vectors, and a host-vector system for gene cloning in Pseudomonas. Gene 16, 237–247 (1981).

66. Nelson, K. E. et al. Complete genome sequence and comparative analysis of the metabolically versatile *Pseudomonas putida* KT2440. Environ. Microbiol. 4, 799–808 (2002).

67. Stover, C. K. et al. Complete genome sequence of *Pseudomonas aeruginosa* PAO1, an opportunistic pathogen. Nature 406, 959–964 (2000).

68. Royle, P. L., Matsumoto, H. & Holloway, B. W. Genetic circularity of the *Pseudomonas aeruginosa* PAO chromosome. J. Bacteriol. 145, 145–155 (1981).

69. Center for Algorithmic Biotechnology. plasmidSPAdes. https://cab.spbu.ru/software/plasmid-spades/ (n.d.).

70. Kearse, M. et al. Geneious Basic: an integrated and extendable desktop software platform for the organization and analysis of sequence data. Bioinformatics 28, 1647–1649 (2012).

71. Li, H. et al. The Sequence Alignment/Map format and SAMtools. Bioinformatics 25, 2078–2079 (2009).

72. Shen, W., Le, S., Li, Y. & Hu, F. SeqKit: a cross-platform and ultrafast toolkit for FASTA/Q file manipulation. PLoS ONE 11, e0163962 (2016).

73. Bankevich, A. et al. SPAdes: a new genome assembly algorithm and its applications to single-cell sequencing. J. Comput. Biol. 19, 455–477 (2012).

74. Sullivan, M. J., Petty, N. K. & Beatson, S. A. Easyfig: a genome comparison visualizer. Bioinformatics 27, 1009–1010 (2011).

75. Bharat, A. et al. Correlation between phenotypic and in silico detection of antimicrobial resistance in *Salmonella enterica* in Canada using Staramr. Microorganisms 10, 292 (2022).

76. Zankari, E. et al. Identification of acquired antimicrobial resistance genes. J. Antimicrob. Chemother. 67, 2640–2644 (2012).

77. Feldgarden, M. et al. AMRFinderPlus and the Reference Gene Catalog facilitate examination of the genomic links among antimicrobial resistance, stress response, and virulence. Sci. Rep. 11, 12728 (2021).

78. Lobry, J. R. Origin of replication of *Mycoplasma genitalium*. Science 272, 745–746 (1996).

79. Necşulea, A. & Lobry, J. R. A new method for assessing the effect of replication on DNA base composition asymmetry. Mol. Biol. Evol. 24, 2169–2179 (2007).

80. Solovyev, V. & Salamov, A. Automatic annotation of microbial genomes and metagenomic sequences. In Metagenomics and Its Applications in Agriculture, Biomedicine and Environmental Studies (ed. Li, R. W.) 61–78 (Nova Science Publishers, New York, 2011).

81. Yanisch-Perron, C., Vieira, J. & Messing, J. Improved M13 phage cloning vectors and host strains: nucleotide sequences of the M13mp18 and pUC19 vectors. Gene 33, 103–119 (1985).

82. van Kempen, M. et al. Fast and accurate protein structure search with Foldseek. Nat. Biotechnol. 42, 243–246 (2024).

83. Swan, M. K., Bastia, D. & Davies, C. Crystal structure of π initiator protein–iteron complex of plasmid R6K: implications for initiation of plasmid DNA replication. Proc. Natl Acad. Sci. USA 103, 18481–18486 (2006).

84. Galata, V., Fehlmann, T., Backes, C. & Keller, A. PLSDB: a resource of complete bacterial plasmids. Nucleic Acids Res. 47, D195–D202 (2019).

85. Robertson, J. & Nash, J. H. E. MOB-suite: software tools for clustering, reconstruction and typing of plasmids from draft assemblies. Microb. Genom. 4, e000206 (2018).

86. Carattoli, A. & Hasman, H. PlasmidFinder and in silico pMLST: identification and typing of plasmid replicons in whole-genome sequencing (WGS). Methods Mol. Biol. 2075, 285–294 (2020).

87. Suzuki, H., Moriguchi, K., Shintani, M., Suzuki, M. & Nojiri, H. Insights from public database sequences related to the replication initiation protein TrfA of the IncP-1 plasmid RK2. Plasmid 134, 102756 (2025).

88. Bayliss, S. C., Thorpe, H. A., Coyle, N. M., Sheppard, S. K. & Feil, E. J. PIRATE: a fast and scalable pangenomics toolbox for clustering diverged orthologues in bacteria. GigaScience 8, giz119 (2019).

89. Buchfink, B., Reuter, K. & Drost, H.-G. Sensitive protein alignments at tree-of-life scale using DIAMOND. Nat. Methods 18, 366–368 (2021).

90. Altschul, S. F. et al. Gapped BLAST and PSI-BLAST: a new generation of protein database search programs. Nucleic Acids Res. 25, 3389–3402 (1997).

91. Steinegger, M. & Söding, J. MMseqs2 enables sensitive protein sequence searching for the analysis of massive data sets. Nat. Biotechnol. 35, 1026–1028 (2017).

92. Woodcroft, B. J., Boyd, J. A. & Tyson, G. W. OrfM: a fast open reading frame predictor for metagenomic data. Bioinformatics 32, 2702–2703 (2016).

93. Eddy, S. R. Accelerated profile HMM searches. PLoS Comput. Biol. 7, e1002195 (2011).

94. Steinegger, M. et al. HH-suite3 for fast remote homology detection and deep protein annotation. BMC Bioinformatics 20, 473 (2019).

95. Katoh, K. & Standley, D. M. MAFFT multiple sequence alignment software version 7: improvements in performance and usability. Mol. Biol. Evol. 30, 772–780 (2013).

96. Minh, B. Q. et al. IQ-TREE 2: new models and efficient methods for phylogenetic inference in the genomic era. Mol. Biol. Evol. 37, 1530–1534 (2020).

97. Kalyaanamoorthy, S., Minh, B. Q., Wong, T. K. F., von Haeseler, A. & Jermiin, L. S. ModelFinder: fast model selection for accurate phylogenetic estimates. Nat. Methods 14, 587–589 (2017).

98. Hoang, D. T., Chernomor, O., von Haeseler, A., Minh, B. Q. & Vinh, L. S. UFBoot2: improving the ultrafast bootstrap approximation. Mol. Biol. Evol. 35, 518– 522 (2018).

99. Guindon, S. et al. New algorithms and methods to estimate maximum-likelihood phylogenies: assessing the performance of PhyML 3.0. Syst. Biol. 59, 307–321 (2010).

100. Mai, U. & Mirarab, S. TreeShrink: fast and accurate detection of outlier long branches in collections of phylogenetic trees. BMC Genomics 19, 272 (2018).

101. Lemoine, F. & Gascuel, O. Gotree/Goalign: toolkit and Go API to facilitate the development of phylogenetic workflows. NAR Genom. Bioinform. 3, lqab075 (2021).

102. Hyatt, D. et al. Prodigal: prokaryotic gene recognition and translation initiation site identification. BMC Bioinformatics 11, 119 (2010).

103. Letunic, I. & Bork, P. Interactive Tree of Life (iTOL) v6: recent updates to the phylogenetic tree display and annotation tool. Nucleic Acids Res. 52, W78–W82 (2024).

104. Néron, B., et al. MacSyFinder v2: improved modelling and search engine to identify molecular systems in genomes. Peer Community J. 3, e28 (2023).

105. Cury, J., Abby, S. S., Doppelt-Azeroual, O., Néron, B. & Rocha, E. P. C. Identifying conjugative plasmids and integrative conjugative elements with CONJscan. Methods Mol. Biol. 2075, 265–283 (2020).

106. Tesson, F. et al. Systematic and quantitative view of the antiviral arsenal of prokaryotes. Nat. Commun. 13, 2561 (2022).

107. Pansegrau, W. et al. Complete nucleotide sequence of Birmingham IncP alpha plasmids. Compilation and comparative analysis. J. Mol. Biol. 239, 623–663 (1994).

108. Shintani, M., Suzuki, H., Nojiri, H. & Suzuki, M. Precise classification of antimicrobial resistance-associated IncP-2 megaplasmids for molecular epidemiological studies on *Pseudomonas* species. J. Antimicrob. Chemother. 77, 1203–1205 (2022).

109. Llanes, C., Gabant, P., Couturier, M., Bayer, L. & Plesiat, P. Molecular analysis of the replication elements of the broad-host-range RepA/C replicon. Plasmid 36, 26–35 (1996).

110. Fricke, W. F. et al. Comparative genomics of the IncA/C multidrug resistance plasmid family. J. Bacteriol. 191, 4750–4757 (2009).

111. Scholz, P. et al. Complete nucleotide sequence and gene organization of the broad-host-range plasmid RSF1010. Gene 75, 271–288 (1989).

112. Maeda, K. et al. Complete nucleotide sequence of carbazole/dioxin-degrading plasmid pCAR1 in *Pseudomonas resinovorans* strain CA10 indicates its mosaicity and the presence of large catabolic transposon Tn4676. J. Mol. Biol. 326, 21–33 (2003).

113. Shintani, M. et al. Characterization of the replication, maintenance, and transfer features of the IncP-7 plasmid pCAR1, which carries genes involved in carbazole and dioxin degradation. Appl. Environ. Microbiol. 72, 3206–3216 (2006).

114. Takahashi, Y., Shintani, M., Yamane, H. & Nojiri, H. The complete nucleotide sequence of pCAR2: pCAR2 and pCAR1 were structurally identical IncP-7 carbazole degradative plasmids. Biosci. Biotechnol. Biochem. 73, 744–746 (2009).

115. Greated, A., Lambertsen, L., Williams, P. A. & Thomas, C. M. Complete sequence of the IncP-9 TOL plasmid pWW0 from *Pseudomonas putida*. Environ. Microbiol. 4, 856–871 (2002).

116. Sota, M. et al. Genomic and functional analysis of the IncP-9 naphthalene-catabolic plasmid NAH7 and its transposon Tn4655 suggests catabolic gene spread by a tyrosine recombinase. J. Bacteriol. 188, 4057–4067 (2006).

117. Douarre, P.-E., Mallet, L., Radomski, N., Felten, A. & Mistou, M.-Y. Analysis of COMPASS, a new comprehensive plasmid database revealed prevalence of multireplicon and extensive diversity of IncF plasmids. Front. Microbiol. 11, 483 (2020).

