## Supplementary material for "A replication-centered phylogeny illuminates the evolutionary landscape of bacterial plasmids": Figures_S1-S17

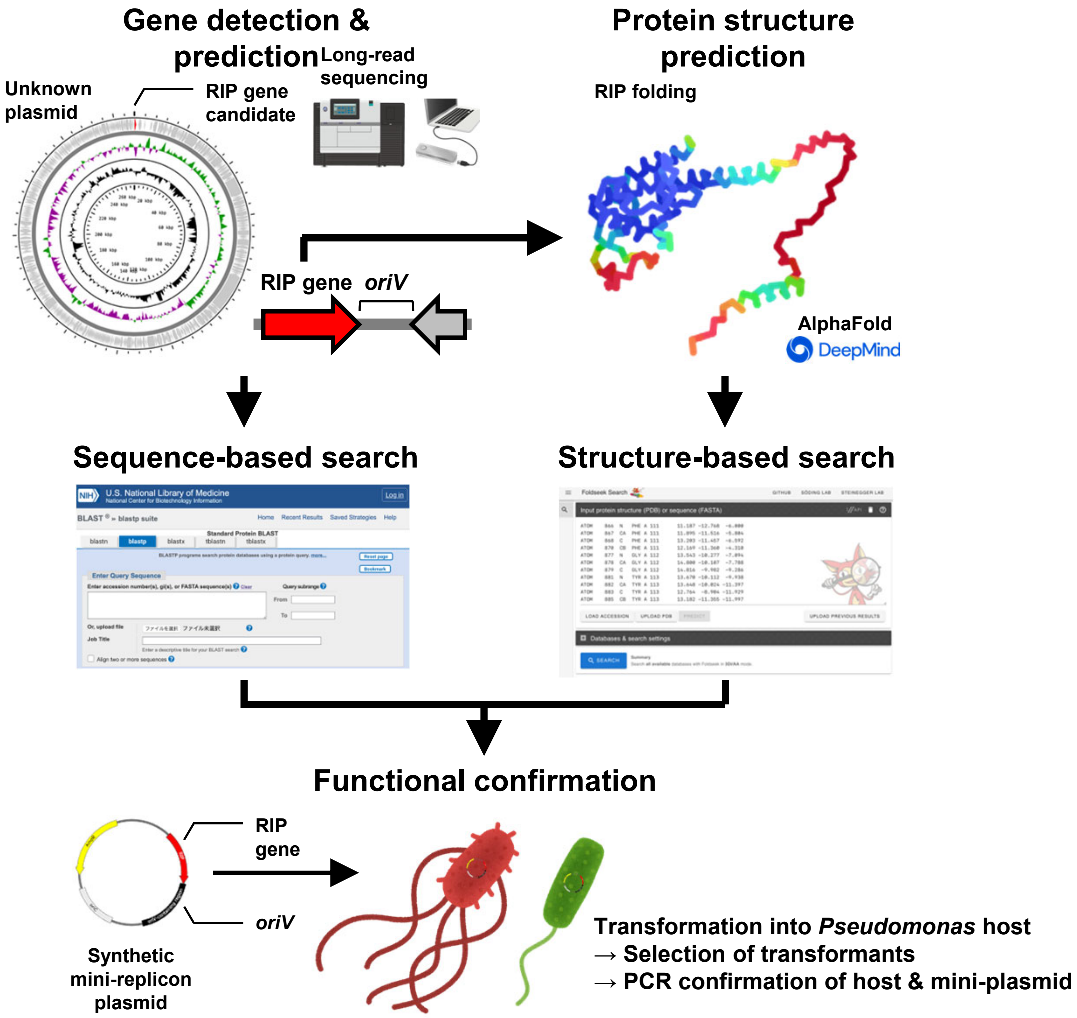

Gene detection & prediction
Long-read sequencing
Unknown plasmid
RIP gene candidate
Protein structure prediction
RIP folding
RIP gene
oriV
AlphaFold
DeepMind
Sequence-based search
Structure-based search
Functional confirmation
RIP gene
oriV
Synthetic mini-replicon plasmid
Transformation into Pseudomonas host
→ Selection of transformants
→ PCR confirmation of host & mini-plasmid

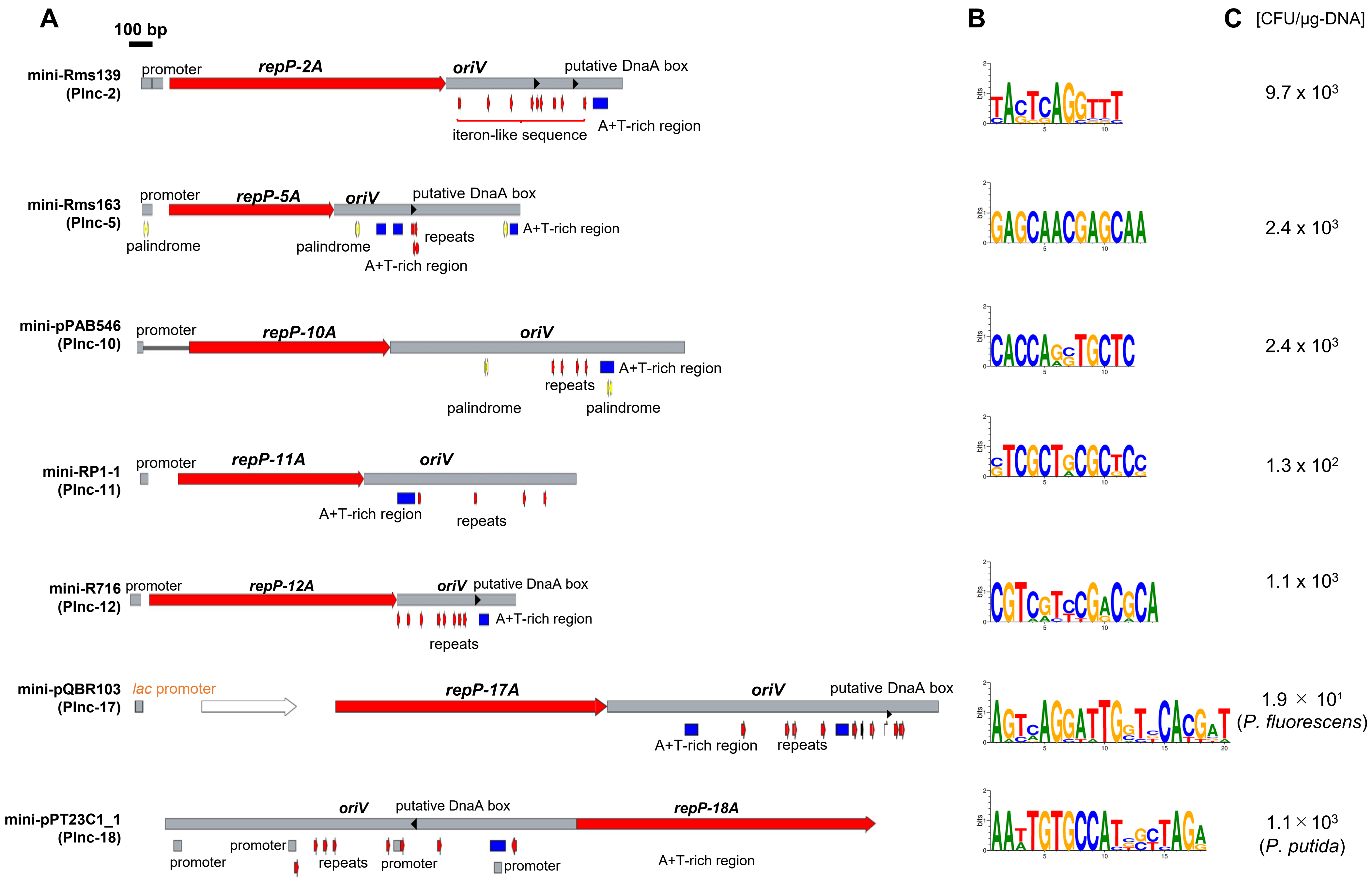
A
100 bp
mini-Rms139 (PInc-2)
promoter
repP-2A
oriV
putative DnaA box
iteron-like sequence
A+T-rich region
mini-Rms163 (PInc-5)
promoter
repP-5A
oriV
putative DnaA box
palindrome
palindrome
repeats
A+T-rich region
A+T-rich region
mini-pPAB546 (PInc-10)
promoter
repP-10A
oriV
A+T-rich region
repeats
palindrome
palindrome
mini-RP1-1 (PInc-11)
promoter
repP-11A
oriV
A+T-rich region
repeats
mini-R716 (PInc-12)
promoter
repP-12A
oriV
putative DnaA box
A+T-rich region
repeats
mini-pQBR103 (PInc-17)
lac promoter
repP-17A
oriV
putative DnaA box
A+T-rich region
repeats
mini-pPT23C1_1 (PInc-18)
oriV
putative DnaA box
repP-18A
promoter
promoter
repeats
promoter
promoter
A+T-rich region
B
C
[CFU/μg-DNA]
9.7 x 10^3
2.4 x 10^3
2.4 x 10^3
1.3 x 10^2
1.1 x 10^3
1.9 × 10^1 (P. fluorescens)
1.1 × 10^3 (P. putida)

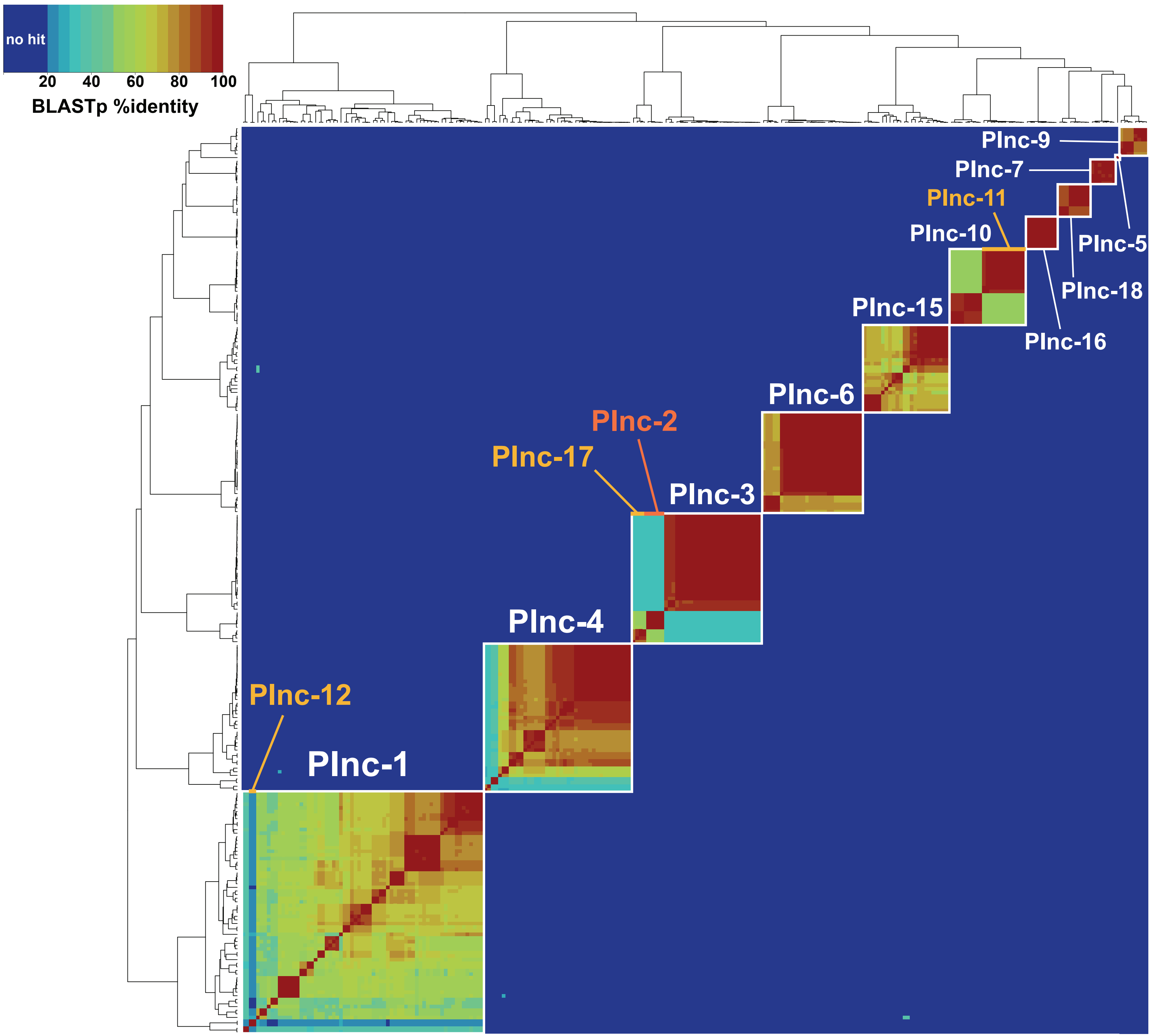

no hit
20
40
60
80
100
BLASTp %identity
Plnc-9
Plnc-7
Plnc-11
Plnc-10
Plnc-5
Plnc-18
Plnc-15
Plnc-16
Plnc-6
Plnc-2
Plnc-17
Plnc-3
Plnc-4
Plnc-12
Plnc-1

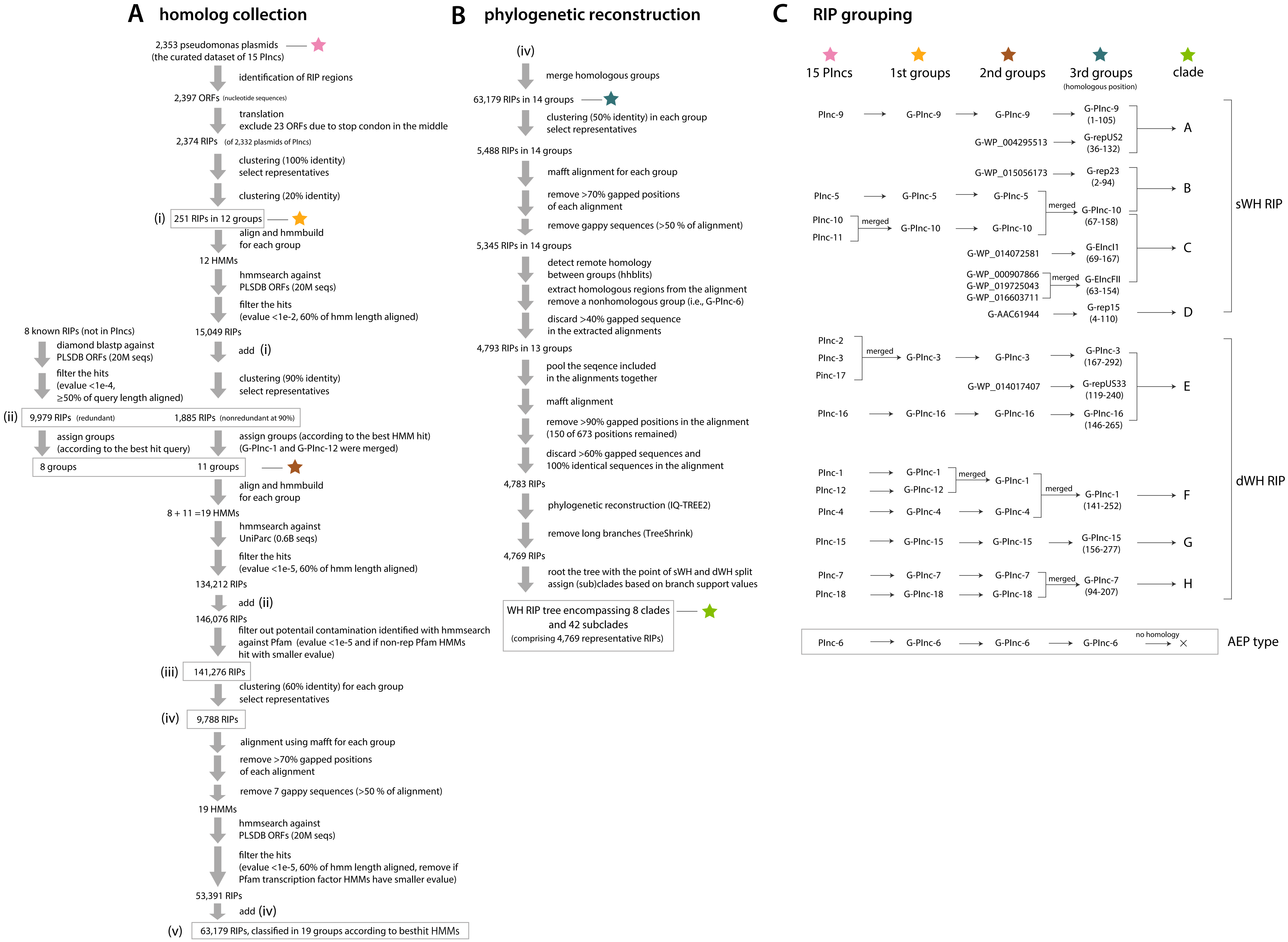

A homolog collection
2,353 pseudomonas plasmids
(the curated dataset of 15 PIncs)
identification of RIP regions
2,397 ORFs (nucleotide sequences)
translation
exclude 23 ORFs due to stop condon in the middle
2,374 RIPs (of 2,332 plasmids of PIncs)
clustering (100% identity)
select representatives
clustering (20% identity)
(i) 251 RIPs in 12 groups
align and hmmbuild
for each group
12 HMMs
hmmsearch against
PLSDB ORFs (20M seqs)
filter the hits
(evalue <1e-2, 60% of hmm length aligned)
15,049 RIPs
add (i)
clustering (90% identity)
select representatives
8 known RIPs (not in PIncs)
diamond blastp against
PLSDB ORFs (20M seqs)
filter the hits
(evalue <1e-4,
≥50% of query length aligned)
(ii) 9,979 RIPs (redundant)
1,885 RIPs (nonredundant at 90%)
assign groups
(according to the best hit query)
assign groups (according to the best HMM hit)
(G-PInc-1 and G-PInc-12 were merged)
8 groups
11 groups
align and hmmbuild
for each group
8 + 11 =19 HMMs
hmmsearch against
UniParc (0.6B seqs)
filter the hits
(evalue <1e-5, 60% of hmm length aligned)
134,212 RIPs
add (ii)
146,076 RIPs
filter out potentail contamination identified with hmmsearch
against Pfam (evalue <1e-5 and if non-rep Pfam HMMs
hit with smaller evalue)
(iii) 141,276 RIPs
clustering (60% identity) for each group
select representatives
(iv) 9,788 RIPs
alignment using mafft for each group
remove >70% gapped positions
of each alignment
remove 7 gappy sequences (>50 % of alignment)
19 HMMs
hmmsearch against
PLSDB ORFs (20M seqs)
filter the hits
(evalue <1e-5, 60% of hmm length aligned, remove if
Pfam transcription factor HMMs have smaller evalue)
53,391 RIPs
add (iv)
(v) 63,179 RIPs, classified in 19 groups according to besthit HMMs
B phylogenetic reconstruction
(iv)
merge homologous groups
63,179 RIPs in 14 groups
clustering (50% identity) in each group
select representatives
5,488 RIPs in 14 groups
mafft alignment for each group
remove >70% gapped positions
of each alignment
remove gappy sequences (>50 % of alignment)
5,345 RIPs in 14 groups
detect remote homology
between groups (hhblits)
extract homologous regions from the alignment
remove a nonhomologous group (i.e., G-PInc-6)
discard >40% gapped sequence
in the extracted alignments
4,793 RIPs in 13 groups
pool the seqence included
in the alignments together
mafft alignment
remove >90% gapped positions in the alignment
(150 of 673 positions remained)
discard >60% gapped sequences and
100% identical sequences in the alignment
4,783 RIPs
phylogenetic reconstruction (IQ-TREE2)
remove long branches (TreeShrink)
4,769 RIPs
root the tree with the point of sWH and dWH split
assign (sub)clades based on branch support values
WH RIP tree encompassing 8 clades
and 42 subclades
(comprising 4,769 representative RIPs)
C RIP grouping
15 PIncs
1st groups
2nd groups
3rd groups
(homologous position)
clade
PInc-9
G-PInc-9
G-PInc-9
G-PInc-9
(1-105)
G-WP_004295513
G-repUS2
(36-132)
A
G-WP_015056173
G-rep23
(2-94)
B
PInc-5
G-PInc-5
G-PInc-5
merged
G-PInc-10
(67-158)
PInc-10
merged
G-PInc-10
G-PInc-10
PInc-11
G-WP_014072581
G-EIncI1
(69-167)
C
G-WP_000907866
G-WP_019725043
G-WP_016603711
merged
G-EIncFII
(63-154)
G-AAC61944
G-rep15
(4-110)
D
sWH RIP
PInc-2
PInc-3
Pinc-17
merged
G-PInc-3
G-PInc-3
G-PInc-3
(167-292)
G-WP_014017407
G-repUS33
(119-240)
E
PInc-16
G-PInc-16
G-PInc-16
G-PInc-16
(146-265)
PInc-1
G-PInc-1
merged
G-PInc-1
PInc-12
G-PInc-12
merged
G-PInc-1
(141-252)
F
PInc-4
G-PInc-4
G-PInc-4
PInc-15
G-PInc-15
G-PInc-15
G-PInc-15
(156-277)
G
PInc-7
G-PInc-7
G-PInc-7
merged
G-PInc-7
(94-207)
H
PInc-18
G-PInc-18
G-PInc-18
dWH RIP
PInc-6
G-PInc-6
G-PInc-6
G-PInc-6
no homology
AEP type

### A homology detection between the 3rd groups

**value: hhblits %-probability**

**template**

| | | sWH | | | | | | | dWH | | | | | | AEP |
|---|---|---|---|---|---|---|---|---|---|---|---|---|---|---|---|
| | | G-PInc-9 | G-repUS2 | G-rep23 | G-PInc-10 | G-EIncI1 | G-EIncFII | G-rep15 | G-PInc-3 | G-repUS33 | G-PInc-16 | G-PInc-1 | G-PInc-15 | G-PInc-7 | G-PInc-6 |
| **query** sWH | G-PInc-9 | 100 | 99.4 | 98.8 | 97.3 | 96 | 96.8 | 99 | 97.4 | 18.6 | 92.5 | 96.9 | 56.8 | 72.6 | 0.2 |
| | G-repUS2 | 99.5 | 99.9 | 98.5 | 97.4 | 96.6 | 97.3 | 98.8 | 92.4 | 10.4 | 87.8 | 97.4 | 75.9 | 90.7 | 0.1 |
| | G-rep23 | 99 | 98.5 | 99.9 | 89.4 | 73.8 | 89 | 99.8 | 98.2 | 7.6 | 93.6 | 97.4 | 70.5 | 85.5 | 0 |
| | G-PInc-10 | 98.6 | 98.3 | 96.9 | 100 | 99.6 | 99.8 | 98.9 | 55.4 | 3.6 | 8 | 61.2 | 3 | 6.9 | 0.1 |
| | G-EIncI1 | 98.1 | 97.9 | 96.9 | 99.8 | 100 | 99.9 | 93.3 | 68.2 | 1.4 | 9.6 | 38.3 | 1.9 | 7.9 | 0.1 |
| | G-EIncFII | 98.7 | 98.4 | 91.5 | 99.8 | 99.9 | 100 | 92.9 | 57.5 | 7.2 | 11.6 | 48.8 | 4.1 | 8.1 | 0.4 |
| | G-rep15 | 99.2 | 98.9 | 99.7 | 97.7 | 95.7 | 96.5 | 100 | 70.4 | 5.1 | 55.9 | 91.2 | 12.2 | 41.4 | 0.1 |
| dWH | G-PInc-3 | 97.3 | 93.6 | 97.8 | 37.9 | 14.2 | 64.3 | 72.4 | 100 | 98.1 | 100 | 99.8 | 99.9 | 99.5 | 0.1 |
| | G-repUS33 | 93.8 | 91.5 | 94.9 | 38.9 | 91.9 | 96 | 1.6 | 99.6 | 100 | 99.4 | 99.5 | 99.2 | 99.2 | 0.1 |
| | G-PInc-16 | 92.2 | 68 | 74.9 | 7.8 | 1.2 | 25.5 | 31.5 | 100 | 95.9 | 100 | 99.9 | 99.9 | 99.5 | 0.3 |
| | G-PInc-1 | 96.1 | 85.2 | 97.6 | 42.4 | 11.5 | 65.1 | 89.5 | 99.9 | 97.2 | 99.9 | 100 | 99.8 | 99.4 | 0.1 |
| | G-PInc-15 | 64.5 | 26.2 | 3.9 | 1.6 | 0.8 | 11.8 | 11.8 | 99.9 | 95.2 | 99.9 | 99.9 | 100 | 99.4 | 0.2 |
| | G-PInc-7 | 85.8 | 36.6 | 82.8 | 2.7 | 6.3 | 24.8 | 26.7 | 99.9 | 98.8 | 99.8 | 99.8 | 99.8 | 100 | 0.1 |
| AEP | G-PInc-6 | 0.6 | 0.2 | 0.2 | 0.3 | 0.6 | 0.3 | 0.7 | 0.2 | 0.1 | 0.3 | 0.5 | 0.3 | 0.2 | 100 |

**value: hhblits hit region**

**template**

| | | sWH | | | | | | | dWH | | | | | | AEP |
|---|---|---|---|---|---|---|---|---|---|---|---|---|---|---|---|
| | | G-PInc-9 | G-repUS2 | G-rep23 | G-PInc-10 | G-EIncI1 | G-EIncFII | G-rep15 | G-PInc-3 | G-repUS33 | G-PInc-16 | G-PInc-1 | G-PInc-15 | G-PInc-7 | G-PInc-6 |
| **query** sWH | G-PInc-9 | 1-144 | 1-101 | 1-92 | 23-93 | 23-95 | 23-96 | 1-94 | 12-100 | 8-71 | 14-80 | 12-89 | 14-70 | 4-76 | 43-58 |
| | G-repUS2 | 36-131 | 1-135 | 36-123 | 51-125 | 52-125 | 32-124 | 35-125 | 34-126 | 33-104 | 30-111 | 15-120 | 31-103 | 39-109 | 101-106 |
| | G-rep23 | 2-99 | 1-96 | 1-226 | 34-98 | 18-95 | 20-97 | 2-226 | 2-98 | 5-65 | 2-76 | 3-87 | 5-64 | 3-68 | 35-93 |
| | G-PInc-10 | 64-170 | 61-165 | 71-164 | 1-232 | 2-182 | 9-185 | 68-148 | 45-164 | 45-143 | 71-135 | 98-133 | 61-125 | 65-131 | 148-167 |
| | G-EIncI1 | 66-182 | 62-168 | 66-172 | 2-190 | 1-299 | 6-193 | 63-163 | 59-169 | 69-151 | 70-142 | 63-152 | 68-109 | 63-168 | 123-133 |
| | G-EIncFII | 60-162 | 57-152 | 63-156 | 8-177 | 5-179 | 1-263 | 61-150 | 35-148 | 61-157 | 58-135 | 25-129 | 53-97 | 61-194 | 142-164 |
| | G-rep15 | 4-114 | 2-113 | 3-270 | 23-112 | 5-113 | 7-113 | 1-272 | 6-111 | 7-84 | 19-93 | 21-105 | 22-83 | 7-89 | 56-72 |
| dWH | G-PInc-3 | 200-311 | 200-311 | 197-313 | 224-289 | 236-301 | 181-301 | 207-303 | 1-316 | 71-315 | 1-292 | 13-283 | 48-308 | 79-305 | 236-252 |
| | G-repUS33 | 167-237 | 167-241 | 166-282 | 200-241 | 181-293 | 173-241 | 173-273 | 22-285 | 1-333 | 3-258 | 7-254 | 182-315 | 19-277 | 227-287 |
| | G-PInc-16 | 187-250 | 184-249 | 177-254 | 211-259 | 88-174 | 159-265 | 187-251 | 4-262 | 57-262 | 1-268 | 5-267 | 39-267 | 59-263 | 160-237 |
| | G-PInc-1 | 166-244 | 165-239 | 163-247 | 196-247 | 181-239 | 146-238 | 172-251 | 19-248 | 63-234 | 13-250 | 1-252 | 46-251 | 62-246 | 16-38 |
| | G-PInc-15 | 201-280 | 178-279 | 187-266 | 223-283 | 80-185 | 78-186 | 194-259 | 45-284 | 73-270 | 31-271 | 20-275 | 1-285 | 72-268 | 223-244 |
| | G-PInc-7 | 121-200 | 108-218 | 119-223 | 153-206 | 136-223 | 100-198 | 127-200 | 4-225 | 11-225 | 13-205 | 6-206 | 8-222 | 1-265 | 109-183 |
| AEP | G-PInc-6 | 292-310 | 26-41 | 115-128 | 221-309 | 212-264 | 292-313 | 290-311 | 292-309 | 15-155 | 50-127 | 91-179 | 95-148 | 268-281 | 1-364 |

### B extraction of homologous region

**sWH** Extract regions of G-PInc-9 that are homologous with dWH. Then, for other groups, extract homologous regions hit to 1-105 of G-PInc-9.

G-PInc-9
- 12-100 --> G-PInc-3
- 14-80 --> G-PInc-16
- 12-89 --> G-PInc-1
- 14-70 --> G-PInc-15
- 4-76 --> G-PInc-7

take union → 4-100 → take margin (5aa for both end) → 1-105 of G-PInc-9 →

- 36-132 of G-repUS2
- 2-94 of G-rep23
- 67-158 of G-PInc-1
- 69-167 of G-EIncI1
- 63-154 of G-EIncFII
- 4-110 of G-rep15

**dWH** Eextract regions of G-PInc-1, homologous with sWH. Then, for other groups, extract homologous regions hit to 141-256 of G-PInc-1.

G-PInc-1
- 166-244 --> G-PInc-9
- 165-239 --> G-repUS2
- 163-247 --> G-rep23
- 146-238 --> G-EIncFII
- 172-251 --> G-rep15

take union → 146-251 → take margin (5aa for both end) → 141-256 of G-PInc-1 →

- 167-292 of G-PInc3
- 119-240 of G-repUS33
- 146-265 of G-PInc-16
- 156-277 of G-PInc-15
- 94-207 of G-PInc-7

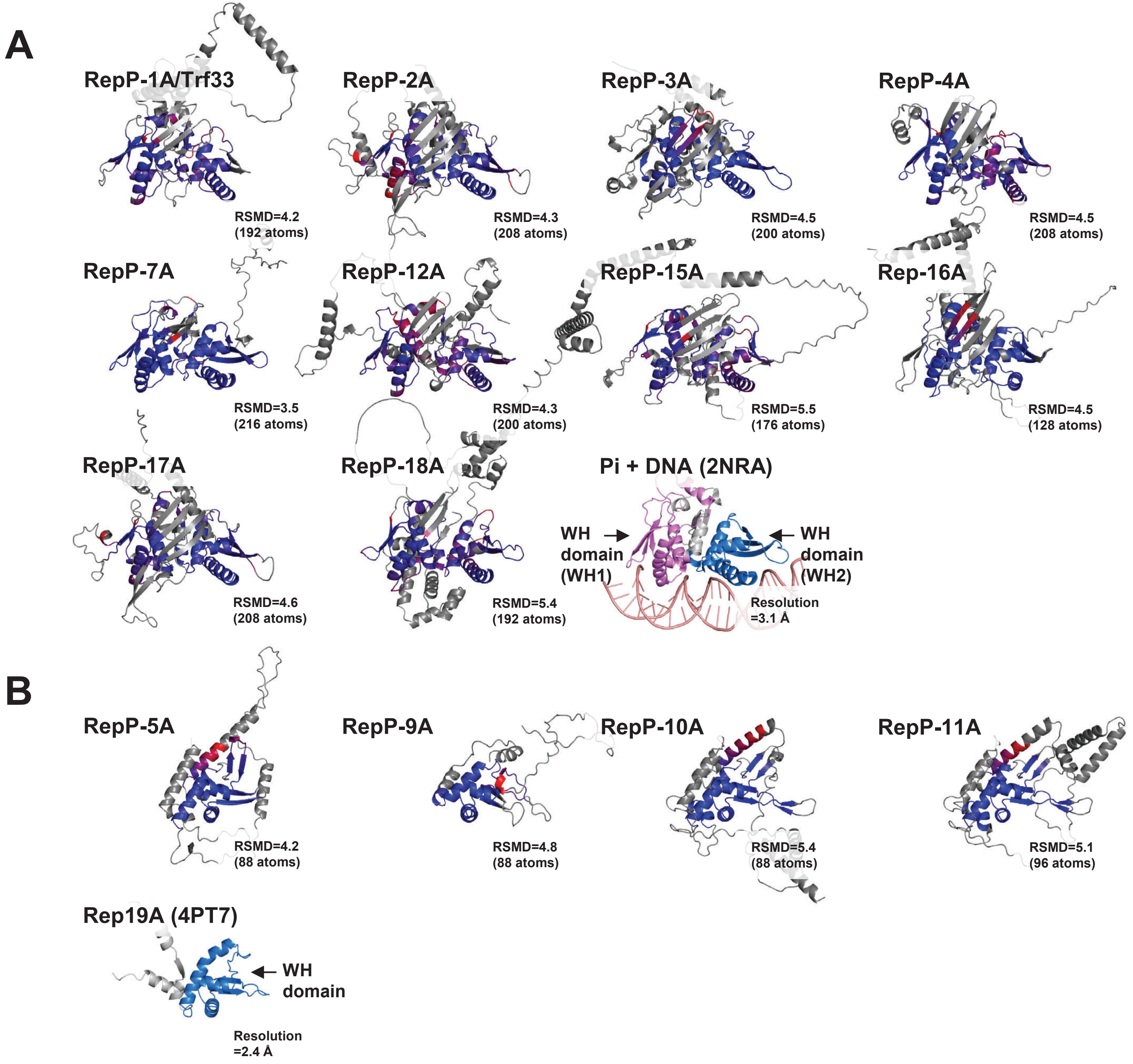
A
RepP-1A/Trf33
RSMD=4.2
(192 atoms)
RepP-2A
RSMD=4.3
(208 atoms)
RepP-3A
RSMD=4.5
(200 atoms)
RepP-4A
RSMD=4.5
(208 atoms)
RepP-7A
RSMD=3.5
(216 atoms)
RepP-12A
RSMD=4.3
(200 atoms)
RepP-15A
RSMD=5.5
(176 atoms)
Rep-16A
RSMD=4.5
(128 atoms)
RepP-17A
RSMD=4.6
(208 atoms)
RepP-18A
RSMD=5.4
(192 atoms)
Pi + DNA (2NRA)
WH domain (WH1)
WH domain (WH2)
Resolution =3.1 Å
B
RepP-5A
RSMD=4.2
(88 atoms)
RepP-9A
RSMD=4.8
(88 atoms)
RepP-10A
RSMD=5.4
(88 atoms)
RepP-11A
RSMD=5.1
(96 atoms)
Rep19A (4PT7)
WH domain
Resolution =2.4 Å

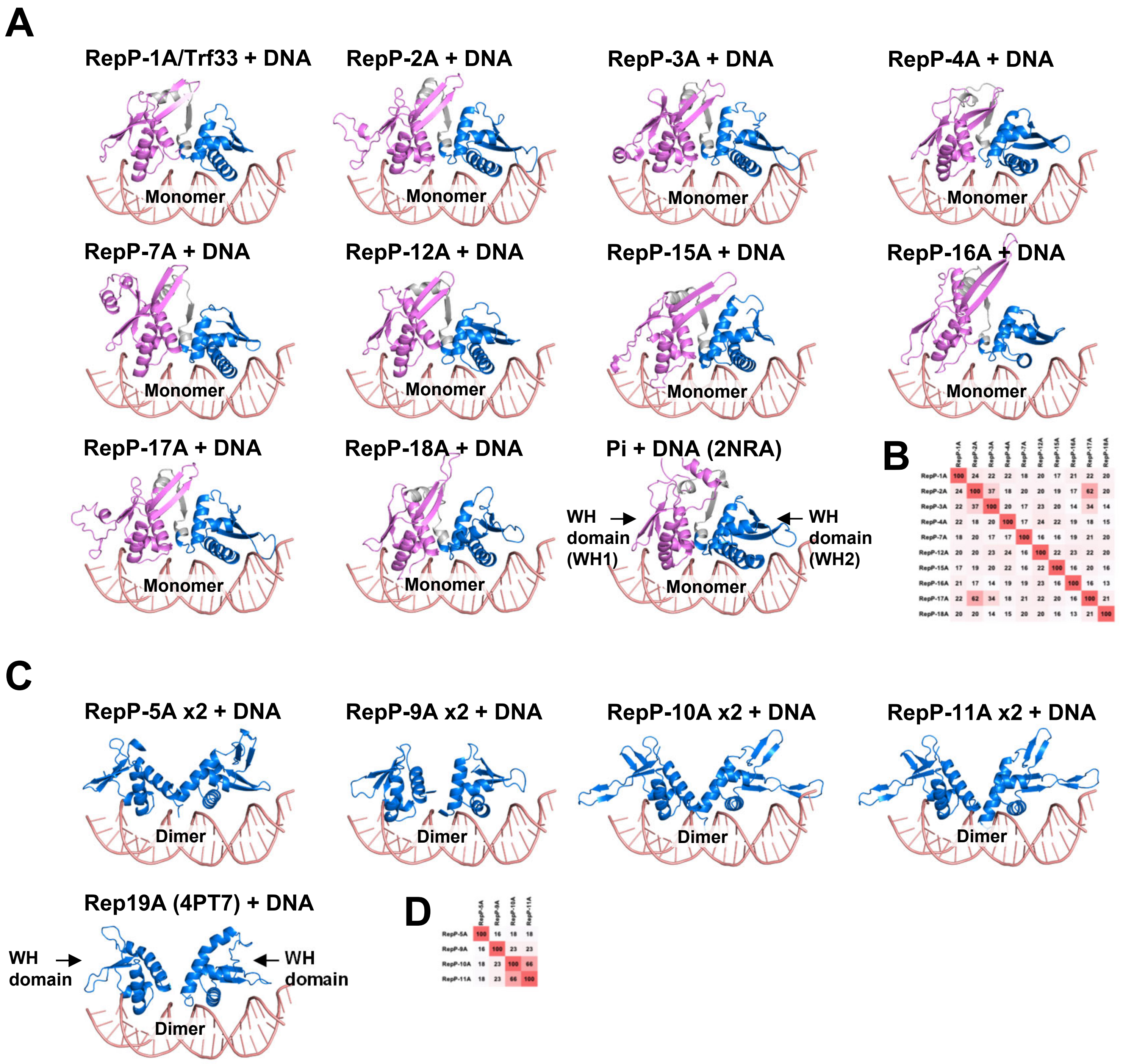


B

| | RepP-1A | RepP-2A | RepP-3A | RepP-4A | RepP-7A | RepP-12A | RepP-15A | RepP-16A | RepP-17A | RepP-18A |
|---|---|---|---|---|---|---|---|---|---|---|
| RepP-1A | 100 | 24 | 22 | 22 | 18 | 20 | 17 | 21 | 22 | 20 |
| RepP-2A | 24 | 100 | 37 | 18 | 20 | 20 | 19 | 17 | 62 | 20 |
| RepP-3A | 22 | 37 | 100 | 20 | 17 | 23 | 20 | 14 | 34 | 14 |
| RepP-4A | 22 | 18 | 20 | 100 | 17 | 24 | 22 | 19 | 18 | 15 |
| RepP-7A | 18 | 20 | 17 | 17 | 100 | 16 | 16 | 19 | 21 | 20 |
| RepP-12A | 20 | 20 | 23 | 24 | 16 | 100 | 22 | 23 | 22 | 20 |
| RepP-15A | 17 | 19 | 20 | 22 | 16 | 22 | 100 | 16 | 20 | 16 |
| RepP-16A | 21 | 17 | 14 | 19 | 19 | 23 | 16 | 100 | 16 | 13 |
| RepP-17A | 22 | 62 | 34 | 18 | 21 | 22 | 20 | 16 | 100 | 21 |
| RepP-18A | 20 | 20 | 14 | 15 | 20 | 20 | 16 | 13 | 21 | 100 |

D

| | RepP-5A | RepP-9A | RepP-10A | RepP-11A |
|---|---|---|---|---|
| RepP-5A | 100 | 16 | 18 | 18 |
| RepP-9A | 16 | 100 | 23 | 23 |
| RepP-10A | 18 | 23 | 100 | 66 |
| RepP-11A | 18 | 23 | 66 | 100 |

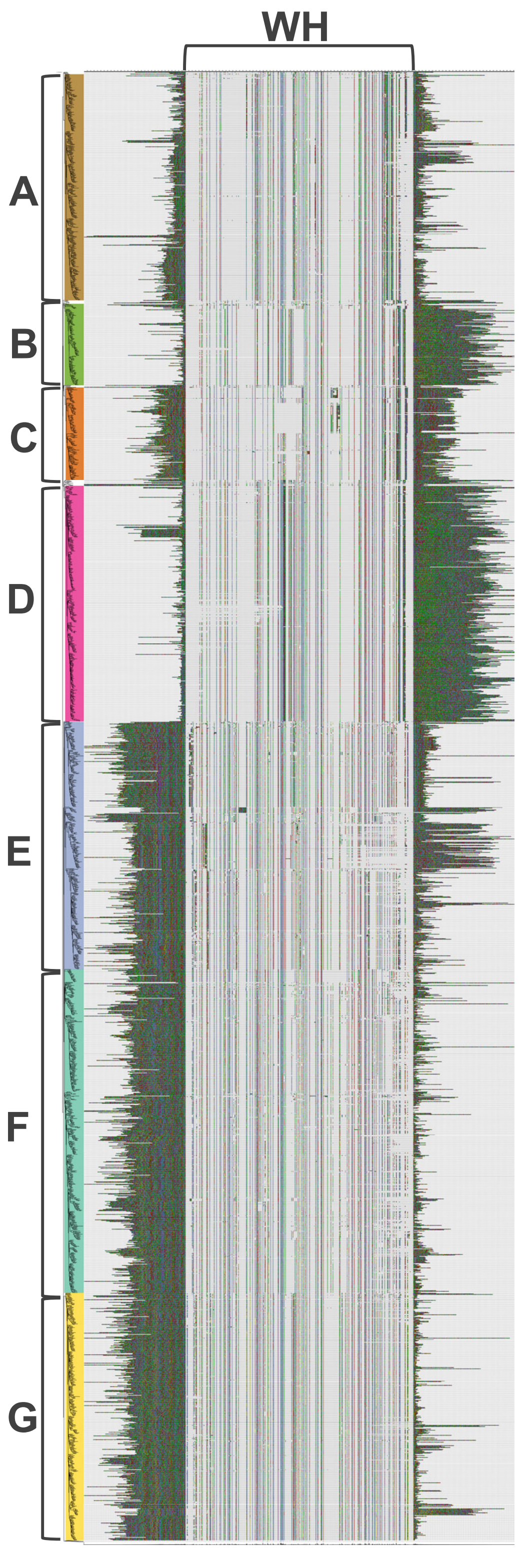
WH
A
B
C
D
E
F
G

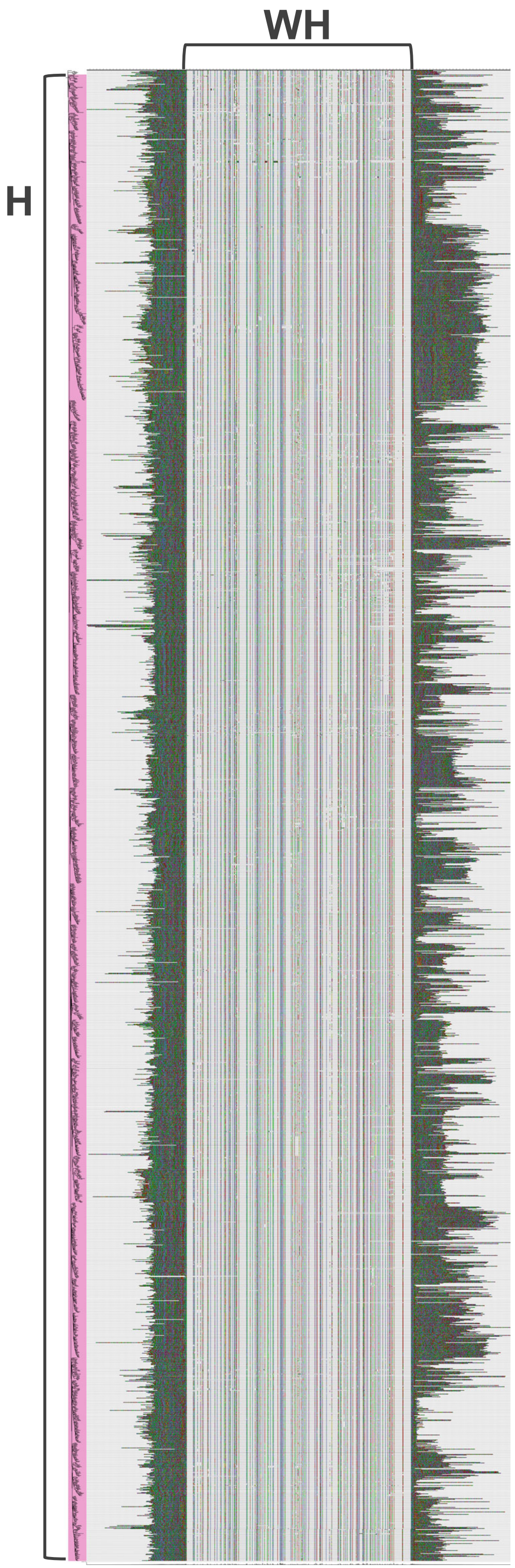
WH
H

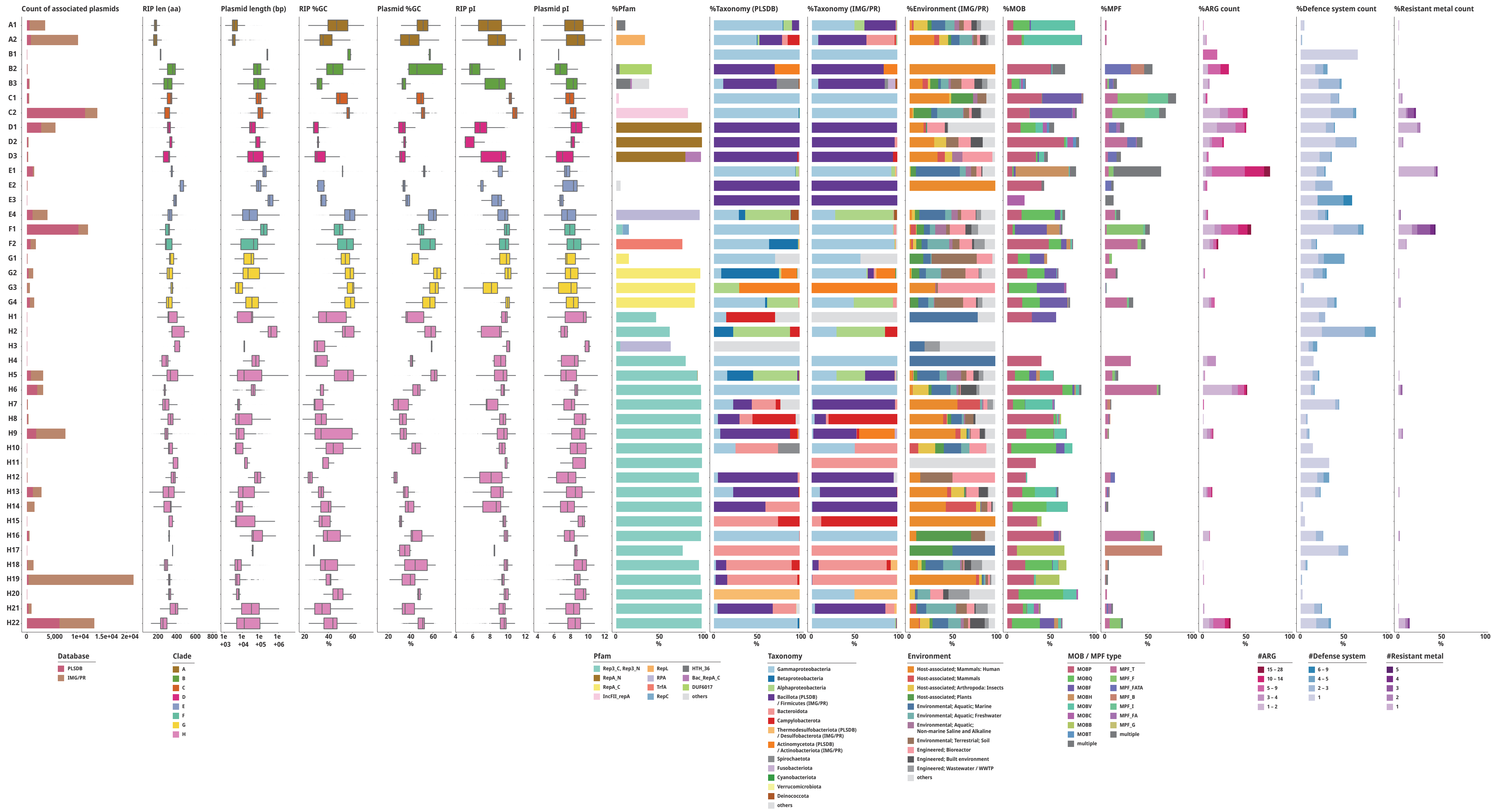

Count of associated plasmids
RIP len (aa)
Plasmid length (bp)
RIP %GC
Plasmid %GC
RIP pI
Plasmid pI
%Pfam
%Taxonomy (PLSDB)
%Taxonomy (IMG/PR)
%Environment (IMG/PR)
%MOB
%MPF
%ARG count
%Defence system count
%Resistant metal count
A1
A2
B1
B2
B3
C1
C2
D1
D2
D3
E1
E2
E3
E4
F1
F2
G1
G2
G3
G4
H1
H2
H3
H4
H5
H6
H7
H8
H9
H10
H11
H12
H13
H14
H15
H16
H17
H18
H19
H20
H21
H22
0 5,000 1e+04 1.5e+04 2e+04
200 400 600 800
1e+03 1e+04 1e+05 1e+06
20 40 60
%
20 40 60
%
4 6 8 10 12
4 6 8 10 12
0 50 100
%
Database
PLSDB
IMG/PR
Clade
A
B
C
D
E
F
G
H
Pfam
Rep3_C, Rep3_N
RepA_N
RepA_C
IncFII_repA
RepL
RPA
TrfA
RepC
HTH_36
Bac_RepA_C
DUF6017
others
Taxonomy
Gammaproteobacteria
Betaproteobacteria
Alphaproteobacteria
Bacillota (PLSDB) / Firmicutes (IMG/PR)
Bacteroidota
Campylobacterota
Thermodesulfobacteriota (PLSDB) / Desulfobacterota (IMG/PR)
Actinomycetota (PLSDB) / Actinobacteriota (IMG/PR)
Spirochaetota
Fusobacteriota
Cyanobacteriota
Verrucomicrobiota
Deinococcota
others
Environment
Host-associated; Mammals: Human
Host-associated; Mammals
Host-associated; Arthropoda: Insects
Host-associated; Plants
Environmental; Aquatic; Marine
Environmental; Aquatic; Freshwater
Environmental; Aquatic; Non-marine Saline and Alkaline
Environmental; Terrestrial; Soil
Engineered; Bioreactor
Engineered; Built environment
Engineered; Wastewater / WWTP
others
MOB / MPF type
MOBP
MOBQ
MOBF
MOBH
MOBV
MOBC
MOBB
MOBT
multiple
MPF_T
MPF_F
MPF_FATA
MPF_B
MPF_I
MPF_FA
MPF_G
multiple
#ARG
15 – 28
10 – 14
5 – 9
3 – 4
1 – 2
#Defense system
6 – 9
4 – 5
2 – 3
1
#Resistant metal
5
4
3
2
1

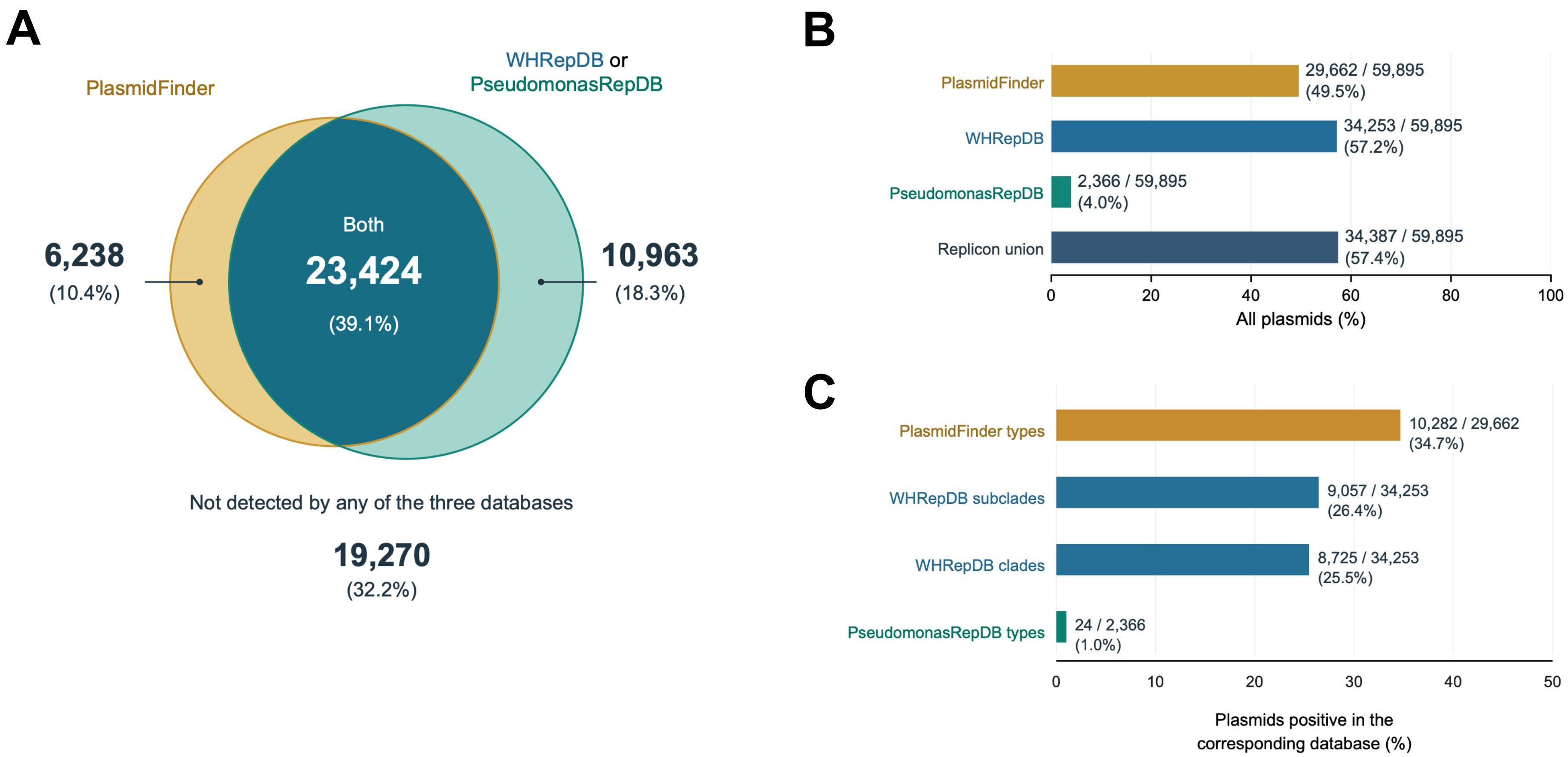
A
PlasmidFinder
WHRepDB or
PseudomonasRepDB
6,238
(10.4%)
Both
23,424
(39.1%)
10,963
(18.3%)
Not detected by any of the three databases
19,270
(32.2%)
B
PlasmidFinder
29,662 / 59,895
(49.5%)
WHRepDB
34,253 / 59,895
(57.2%)
PseudomonasRepDB
2,366 / 59,895
(4.0%)
Replicon union
34,387 / 59,895
(57.4%)
0
20
40
60
80
100
All plasmids (%)
C
PlasmidFinder types
10,282 / 29,662
(34.7%)
WHRepDB subclades
9,057 / 34,253
(26.4%)
WHRepDB clades
8,725 / 34,253
(25.5%)
PseudomonasRepDB types
24 / 2,366
(1.0%)
0
10
20
30
40
50
Plasmids positive in the
corresponding database (%)

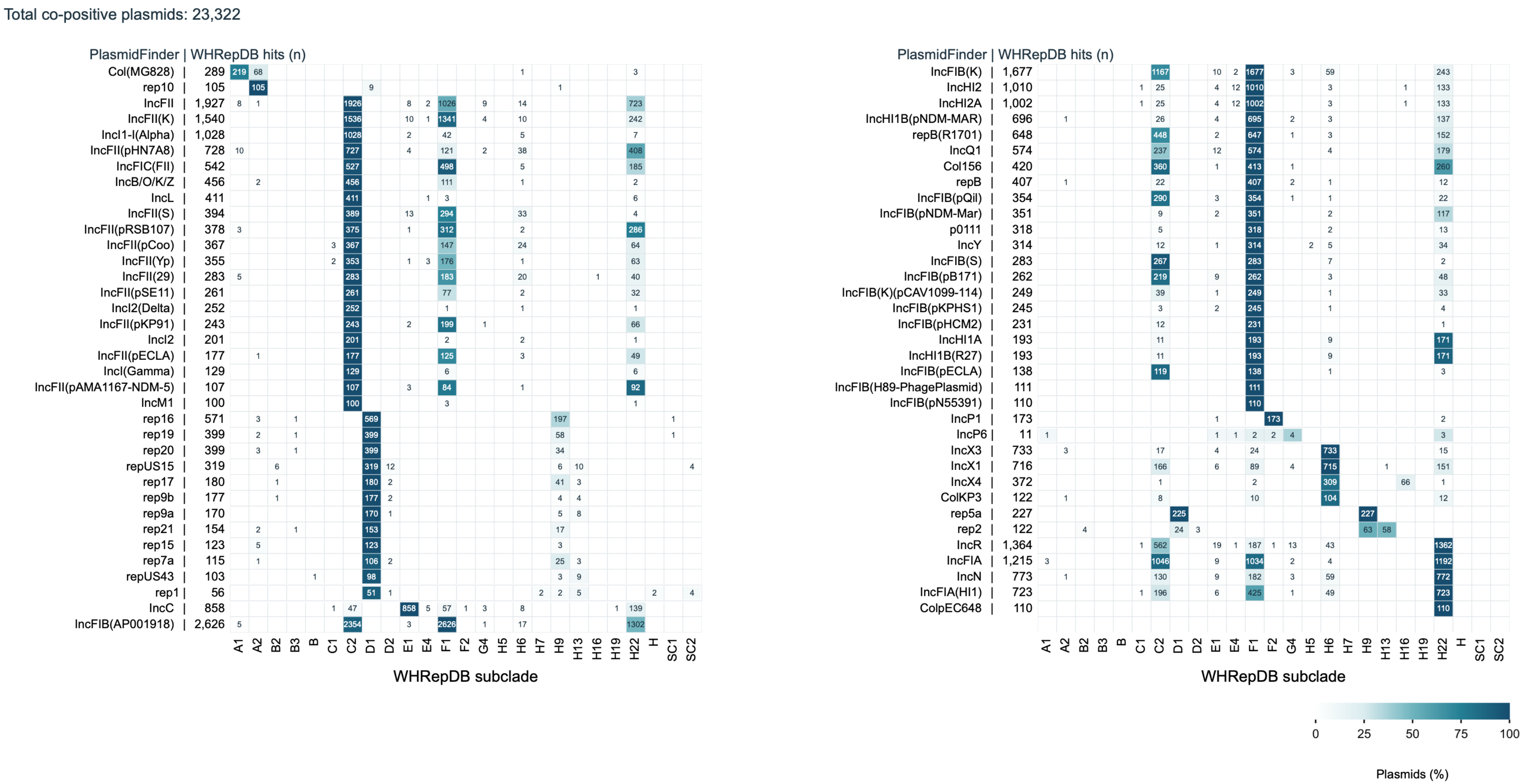
Total co-positive plasmids: 23,322
PlasmidFinder | WHRepDB hits (n)
Col(MG828) | 289
rep10 | 105
IncFII | 1,927
IncFII(K) | 1,540
IncI1-I(Alpha) | 1,028
IncFII(pHN7A8) | 728
IncFIC(FII) | 542
IncB/O/K/Z | 456
IncL | 411
IncFII(S) | 394
IncFII(pRSB107) | 378
IncFII(pCoo) | 367
IncFII(Yp) | 355
IncFII(29) | 283
IncFII(pSE11) | 261
IncI2(Delta) | 252
IncFII(pKP91) | 243
IncI2 | 201
IncFII(pECLA) | 177
IncI(Gamma) | 129
IncFII(pAMA1167-NDM-5) | 107
IncM1 | 100
rep16 | 571
rep19 | 399
rep20 | 399
repUS15 | 319
rep17 | 180
rep9b | 177
rep9a | 170
rep21 | 154
rep15 | 123
rep7a | 115
repUS43 | 103
rep1 | 56
IncC | 858
IncFIB(AP001918) | 2,626
A1 A2 B2 B3 B C1 C2 D1 D2 E1 E4 F1 F2 G4 H5 H6 H7 H9 H13 H16 H19 H22 H SC1 SC2
WHRepDB subclade
PlasmidFinder | WHRepDB hits (n)
IncFIB(K) | 1,677
IncHI2 | 1,010
IncHI2A | 1,002
IncHI1B(pNDM-MAR) | 696
repB(R1701) | 648
IncQ1 | 574
Col156 | 420
repB | 407
IncFIB(pQil) | 354
IncFIB(pNDM-Mar) | 351
p0111 | 318
IncY | 314
IncFIB(S) | 283
IncFIB(pB171) | 262
IncFIB(K)(pCAV1099-114) | 249
IncFIB(pKPHS1) | 245
IncFIB(pHCM2) | 231
IncHI1A | 193
IncHI1B(R27) | 193
IncFIB(pECLA) | 138
IncFIB(H89-PhagePlasmid) | 111
IncFIB(pN55391) | 110
IncP1 | 173
IncP6 | 11
IncX3 | 733
IncX1 | 716
IncX4 | 372
ColKP3 | 122
rep5a | 227
rep2 | 122
IncR | 1,364
IncFIA | 1,215
IncN | 773
IncFIA(HI1) | 723
ColpEC648 | 110
A1 A2 B2 B3 B C1 C2 D1 D2 E1 E4 F1 F2 G4 H5 H6 H7 H9 H13 H16 H19 H22 H SC1 SC2
WHRepDB subclade
0 25 50 75 100
Plasmids (%)

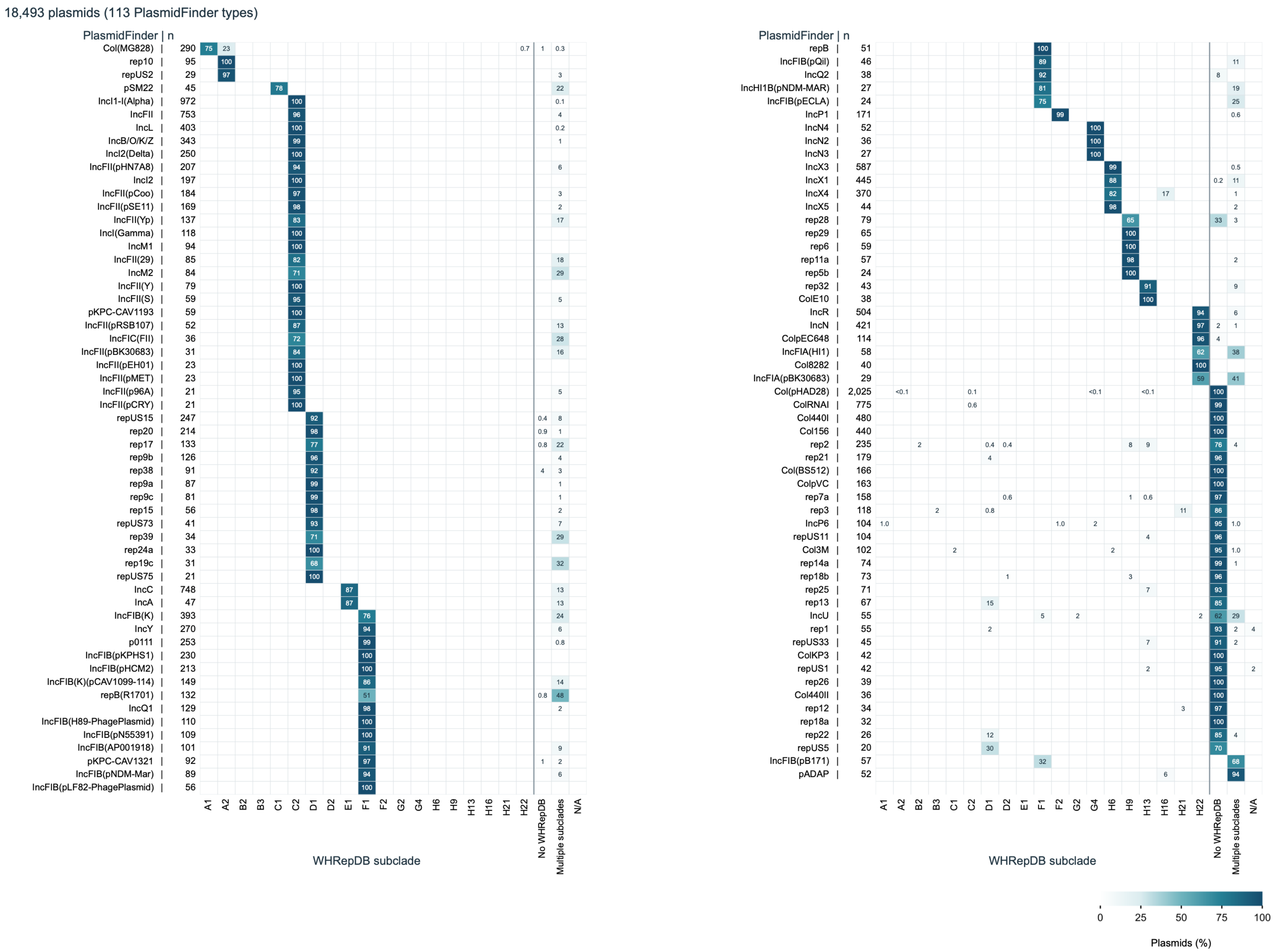
18,493 plasmids (113 PlasmidFinder types)
PlasmidFinder | n
Col(MG828) | 290
rep10 | 95
repUS2 | 29
pSM22 | 45
IncI1-I(Alpha) | 972
IncFII | 753
IncL | 403
IncB/O/K/Z | 343
IncI2(Delta) | 250
IncFII(pHN7A8) | 207
IncI2 | 197
IncFII(pCoo) | 184
IncFII(pSE11) | 169
IncFII(Yp) | 137
IncI(Gamma) | 118
IncM1 | 94
IncFII(29) | 85
IncM2 | 84
IncFII(Y) | 79
IncFII(S) | 59
pKPC-CAV1193 | 59
IncFII(pRSB107) | 52
IncFIC(FII) | 36
IncFII(pBK30683) | 31
IncFII(pEH01) | 23
IncFII(pMET) | 23
IncFII(p96A) | 21
IncFII(pCRY) | 21
repUS15 | 247
rep20 | 214
rep17 | 133
rep9b | 126
rep38 | 91
rep9a | 87
rep9c | 81
rep15 | 56
repUS73 | 41
rep39 | 34
rep24a | 33
rep19c | 31
repUS75 | 21
IncC | 748
IncA | 47
IncFIB(K) | 393
IncY | 270
p0111 | 253
IncFIB(pKPHS1) | 230
IncFIB(pHCM2) | 213
IncFIB(K)(pCAV1099-114) | 149
repB(R1701) | 132
IncQ1 | 129
IncFIB(H89-PhagePlasmid) | 110
IncFIB(pN55391) | 109
IncFIB(AP001918) | 101
pKPC-CAV1321 | 92
IncFIB(pNDM-Mar) | 89
IncFIB(pLF82-PhagePlasmid) | 56
A1 A2 B2 B3 C1 C2 D1 D2 E1 F1 F2 G2 G4 H6 H9 H13 H16 H21 H22 No WHRepDB Multiple subclades N/A
WHRepDB subclade
PlasmidFinder | n
repB | 51
IncFIB(pQil) | 46
IncQ2 | 38
IncHI1B(pNDM-MAR) | 27
IncFIB(pECLA) | 24
IncP1 | 171
IncN4 | 52
IncN2 | 36
IncN3 | 27
IncX3 | 587
IncX1 | 445
IncX4 | 370
IncX5 | 44
rep28 | 79
rep29 | 65
rep6 | 59
rep11a | 57
rep5b | 24
rep32 | 43
ColE10 | 38
IncR | 504
IncN | 421
ColpEC648 | 114
IncFIA(HI1) | 58
Col8282 | 40
IncFIA(pBK30683) | 29
Col(pHAD28) | 2,025
ColRNAI | 775
Col440I | 480
Col156 | 440
rep2 | 235
rep21 | 179
Col(BS512) | 166
ColpVC | 163
rep7a | 158
rep3 | 118
IncP6 | 104
repUS11 | 104
Col3M | 102
rep14a | 74
rep18b | 73
rep25 | 71
rep13 | 67
IncU | 55
rep1 | 55
repUS33 | 45
ColKP3 | 42
repUS1 | 42
rep26 | 39
Col440II | 36
rep12 | 34
rep18a | 32
rep22 | 26
repUS5 | 20
IncFIB(pB171) | 57
pADAP | 52
A1 A2 B2 B3 C1 C2 D1 D2 E1 F1 F2 G2 G4 H6 H9 H13 H16 H21 H22 No WHRepDB Multiple subclades N/A
WHRepDB subclade
0 25 50 75 100
Plasmids (%)

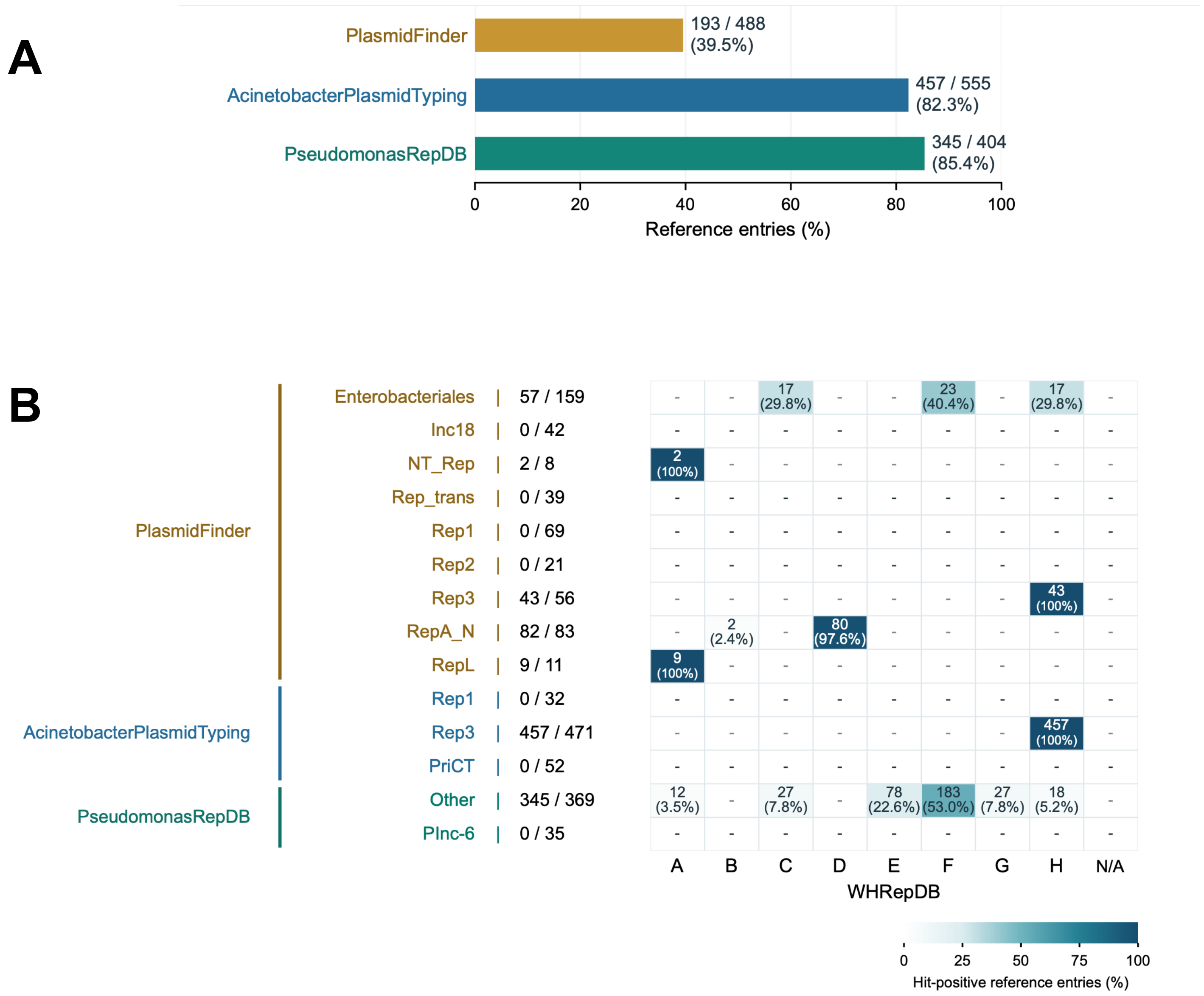
A
PlasmidFinder
AcinetobacterPlasmidTyping
PseudomonasRepDB
193 / 488 (39.5%)
457 / 555 (82.3%)
345 / 404 (85.4%)
Reference entries (%)
B
PlasmidFinder
AcinetobacterPlasmidTyping
PseudomonasRepDB
Enterobacteriales | 57 / 159
Inc18 | 0 / 42
NT_Rep | 2 / 8
Rep_trans | 0 / 39
Rep1 | 0 / 69
Rep2 | 0 / 21
Rep3 | 43 / 56
RepA_N | 82 / 83
RepL | 9 / 11
Rep1 | 0 / 32
Rep3 | 457 / 471
PriCT | 0 / 52
Other | 345 / 369
PInc-6 | 0 / 35
A B C D E F G H N/A
WHRepDB
Hit-positive reference entries (%)

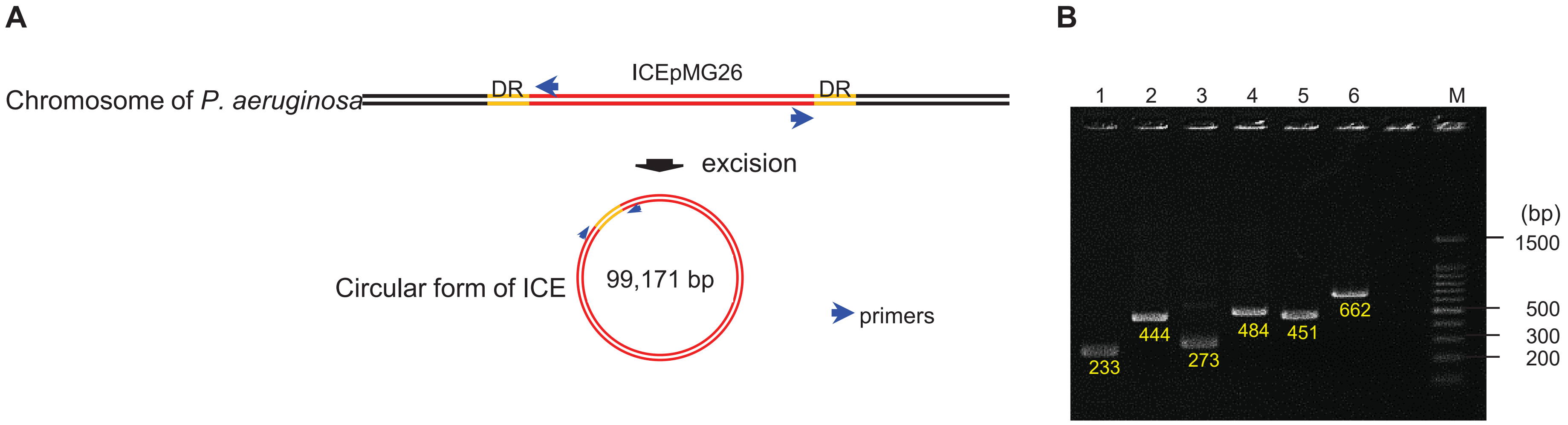
A
Chromosome of P. aeruginosa
DR
ICEpMG26
DR
excision
Circular form of ICE
99,171 bp
primers
B
1
2
3
4
5
6
M
(bp)
1500
500
300
200
233
444
273
484
451
662

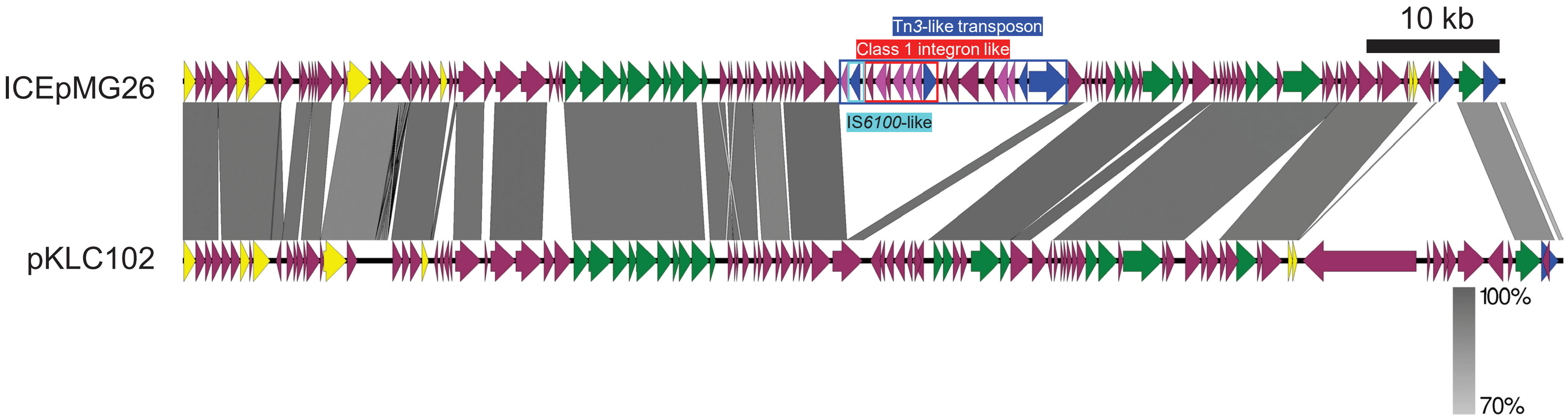

Tn3-like transposon
Class 1 integron like
10 kb
ICEpMG26
IS6100-like
pKLC102
100%
70%

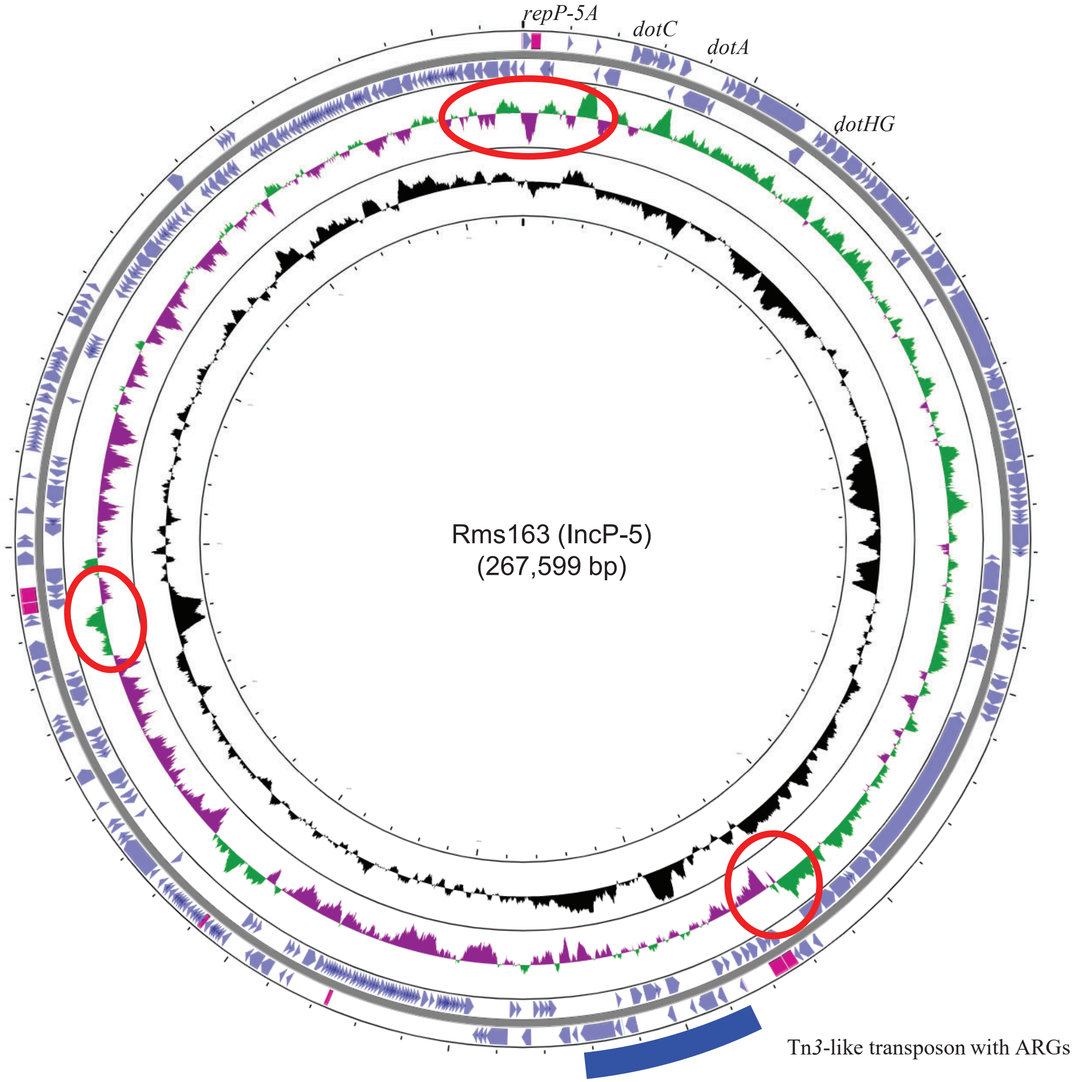

repP-5A
dotC
dotA
dotHG
Rms163 (IncP-5)
(267,599 bp)
Tn3-like transposon with ARGs

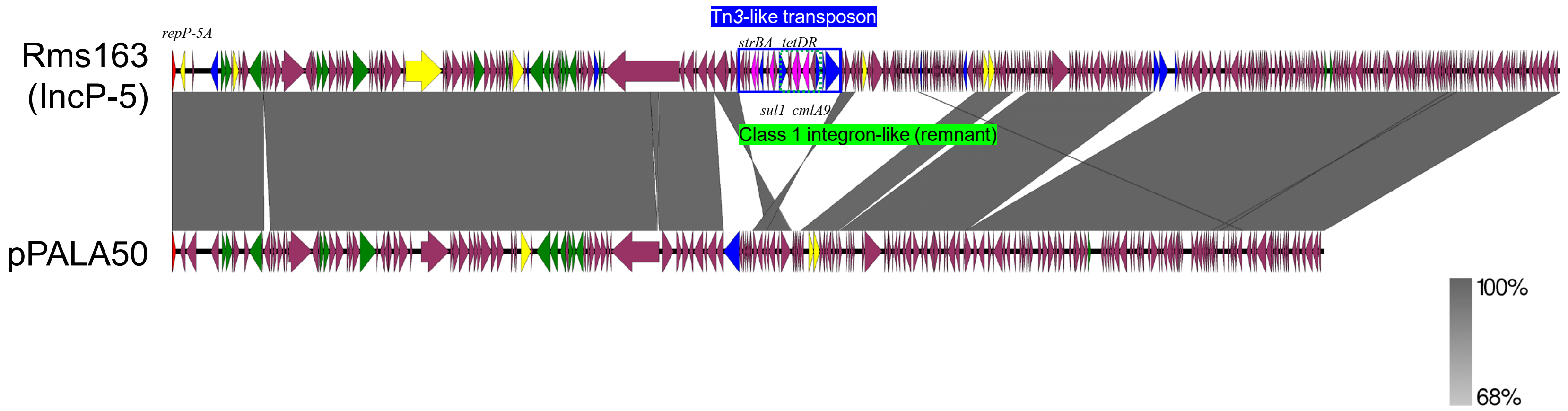

Tn3-like transposon
repP-5A
strBA tetDR
Rms163
(IncP-5)
sul1 cmlA9
Class 1 integron-like (remnant)
pPALA50
100%
68%
