## Supplemental text S1 for "A replication-centered phylogeny illuminates the evolutionary landscape of bacterial plasmids"

**The Supplemental materials include:**

Supplemental Text S1

Supplemental Figures: Figures. S1-S17.

Supplemental Tables (provided in Excel format): Tables S1-S9.

**Supplemental Text S1**

**Identification of the RIP gene in each R plasmid**

The IncP-2 plasmid Rms139 harbours a single RIP gene, called *repP-2A*, with its putative *oriV* identified between nucleotides 2,395 and 3,150^1^. This *oriV* region contains nine repeats of an 11-bp sequence [5’-TA(G/C)TCAGGTTT-3’], two putative DnaA boxes [5’-TT(A/T)(T/G)C(T/C)(A/T)CA-3’], and an AT-rich region (Figure S2). A mini-Rms139A construct, containing the *repP-2A* gene and *oriV*, was successfully introduced into *P. aeruginosa* via electroporation, achieving a transformation frequency of 9.7 × 10³ CFU/μg-DNA. No transformants were detected with the control vector pUC-GW-AMP (<3.4 × 10⁰ CFU/μg-DNA). This result indicates the functional role of *repP-2A* in replication.

Initially, no RIP gene was detected in the IncP-5 plasmid Rms163 through nucleotide sequence-based searches. Subsequently, a GC-skew plot of Rms163 was generated (Figure S16), and three nearby CDSs were selected for three-dimensional structure prediction. One CDS showed structural similarity to the replication initiator A family proteins, specifically to Rep19A of pCA-347 with an RMSD of 4.2 (Figure S6B). Downstream of this gene, a putative *oriV* region was identified with characteristic repeats, palindromic sequences, a putative DnaA box, and an AT-rich region (Figure S2). Transformation of *P. aeruginosa* PAO1 with a mini-Rms163A construct containing the RIP gene candidate and *oriV* resulted in a transformation efficiency of 2.4 x 10^3^ CFU/μg-DNA. This candidate gene was subsequently named *repP-5A*.

Although the complete sequence of the plasmid previously identified as IncP-10 is not available in public databases, a 2.7-kb DNA sequence from the archetype IncP-10 plasmid R91-5^2^ was used as the query. This sequence was previously identified as the replication region of R91-5 through incompatibility testing^3^. We identified 45 plasmids in the PLSDB with regions similar to this 2.7-kb fragment, including an unnamed plasmid from the *P. aeruginosa* strain PAB546 (GenBank accession number NZ_MN433456.1), which we tentatively named pPAB546. While no RIP gene was initially annotated in this plasmid, a manual search similar to that used for IncP-5 led to the identification of a putative RIP similar to Rep19A (RMSD = 5.4) (Figure S6B), encoded by a gene near the conserved 2.7-kb region (Figure S2). The transformation of mini-pPAB546 into *P. aeruginosa* yielded a frequency of 2.4 × 10^3^ CFU/μg-DNA (Figure S2).

The IncP-11 plasmid RP1-1 possessed a putative CDS classified as rep_cluster_2025 by the MOB-suite package^4^, located at nucleotides 54,711-54,953 of the RP1-1 sequence (GenBank accession number LC700336). A nearby *oriV* region featured four 12-bp repeats and an AT-rich region, although no DnaA box was identified (Figure S2). Initial transformation attempts with a pUC-GW-AMP construct harbouring these regions failed to yield transformants (<8.0 × 10 CFU/μg-DNA). However, a downstream CDS, showing 61% identity with the replication initiator protein RepA from a *P. stutzeri* plasmid-like contig, was cloned along with the *oriV* region. The constructed mini-RP1-1A successfully transformed *P. aeruginosa* at a frequency of 1.3 × 10² CFU/μg-DNA (Figure S2), leading to the designation of the CDS as *repP-11A*.

Similar to the IncP-2 plasmid, the IncP-12 plasmid R716 contained a putative RIP gene, *repP-12A*, encoding a trans-acting replication function (TrfA) family protein (pfam07042), which is analogous to the RIPs found in IncP-1 plasmids. Downstream of *repP-12A*, we identified eight repeats of a 14-bp sequence (5’-CGTCGTCCGACGCA-3’), one putative DnaA box (5’-TTATCCATA-3’), and an AT-rich region (Figure S2). A mini-R716A construct containing *repP-12A* and *oriV* was successfully introduced into *P. aeruginosa* via electroporation, with a transformation frequency of 1.1 × 10³ CFU/μg-DNA.

For PInc-17, two *repA–oriV* regions had previously been predicted in pQBR103 but had not been experimentally examined. We therefore constructed mini-replicons containing each candidate replication region and assessed their ability to replicate in *P. fluorescens* SBW25. Both constructs yielded transformants, supporting the replication competence of both predicted *repA–oriV* regions. For the *repP-17A–oriV* construct, *repP-17A* was expressed from a vector-derived *lac* promoter. Transformants were obtained in only one of three independent trials, at a transformation frequency of approximately 1.9 × 10¹ CFU/μg-DNA, which was close to the detection limit of the assay. The presence of the mini-plasmid in the transformants was confirmed by PCR.

Although both *repA–oriV* regions supported replication, we designated *repP-17A–oriV* as the representative replication region of PInc-17 based on its distribution among related plasmids. Plasmids carrying both replication regions are currently known, whereas among plasmids carrying only one of the two regions, only those carrying *repP-17A–oriV* have so far been identified. We therefore used *repP-17A* as the reference RIP for PInc-17 classification.

In addition, we determined the RIP and *oriV* region of the Inc-18 (pSTY-like) plasmid, pPT23-C1_1 (accession number: AP040308). The predicted *oriV* region was located immediately upstream of the RIP gene and comprised multiple direct repeats and an AT-rich region, consistent with iteron-type replication origins (Figure S2). A mini-pPT23-C1_1 containing *repP-18A* and *oriV* was successfully introduced into *P. putida* via electroporation, with a transformation frequency of 1.1 × 10³ CFU/μg-DNA.

**pMG26 as an integrative and conjugative element (ICE)**

The plasmid pMG26, previously identified as a multidrug resistance plasmid and classified as IncP-13^5,6^, did not yield a circular contig during assembly, even after removing reads mapping to the PAO1808 chromosome. Instead, a 99-kb DNA region was detected within the chromosome of its host, PAO1808, which was not conserved in the hosts of other plasmids. This observation led to the hypothesis that pMG26 might function as an ICE, similar to the reclassification of FP8 as ICEPae1161^7^.

To investigate this hypothesis, PCR analyses were performed to determine whether an intermediate circular DNA form of the 99-kb DNA region could be generated in the host PAO1808. As shown in Figure S14, PCR products were obtained using three different primer sets, suggesting that pMG26 could indeed form a circular intermediate within its host. Notably, 49-bp direct repeats (5’-ATGGTGGGTCGTGTAGGATTCGAACCTACGACCAATTGGTTAAAAGCCA-3’) were identified at both ends of the integrated DNA region. The predicted circular form of pMG26 was 99,171 bp in length and was integrated into the *tRNA*_Lys_ locus situated between the *lepA* and *clpB* region in the *P. aeruginosa* PAO1 chromosome (5,086,973 nt, [GenBank: CP114113](https://www.ncbi.nlm.nih.gov/nuccore/NC_002516)). Notably, the entire DNA region of pMG26 exhibited a high degree of similarity with that of pKLC102^8^ (Figure S15), a member of the PAPI-1 family of ICEs, which are abundant in *P. aeruginosa**^8^*.

Further analysis revealed that the 99-kb DNA region of pMG26 encoded a MOB_H_ type relaxase and MPF_G_ family type IV secretion system (T4SS) (Table 1). To test its conjugative transferability, both filter mating and liquid mating assays were performed between *P. aeruginosa* PAO1808 (donor) carrying pMG26 and *P. putida* KT2440 (recipient). The element was successfully transferred from the donor to the recipient strain, with transfer frequencies of 1.4-1.9 × 10^-4^ per donor (filter mating) and 0.28-3.4 × 10^-6^ (liquid mating) per donor (Table S9). Further analyses using MOB-suite indicated that rep_cluster_322 was present in both pMG26 and pKLC102. The putative *oriV* region of pKLC102^8^ was also identified in pMG26. Despite these similarities, no transformants were detected when a mini-pMG26A containing the rep_cluster_322 CDS and putative *oriV* region of pMG26 was introduced into *P. aeruginosa* (below the detection limit, <2.0 × 10 CFU/μg-DNA). This result was not unexpected because the autonomous replication of ICE is typically mediated by rolling-circle replication involving their relaxases^8^. These results strongly suggest that pMG26 functions as an ICE.

**Conjugation assays**

Transferability of five plasmids (Rms139, Rms163, Rsu2, RP1-1, and R716) was experimentally tested by mating assays on solid and liquid conditions. Filter mating assays were performed as described previously^9^. In brief, donor and recipient strains, *P. aeruginosa* PAO1808 with a plasmid and *P.* *aeruginosa* PAO1R or *P*. *putida* KT2440, were cultured in LB with the appropriate antibiotic(s) (preculture). One millilitre of the above culture in LB medium was harvested and suspended in fresh LB medium. Approximately 10^8^ colony forming units/mL (CFU/mL) of the donor and recipient suspended in 150 µL LB were mixed and then dropped on 0.2 µm pore-size filters (cellulose acetate, Advantec) on an LB agar plate for 24 h at 37°C. The mixture on the filter was resuspended in PBS and then spread onto LB plates with appropriate antimicrobials to select transconjugants (i.e., recipient cells with the plasmid). Liquid mating assays were performed as follows. The precultured donor cells were resuspended in 500 µL LB, mixed with 1 mL recipient cells for 2 h (200 rpm), and statically incubated for 3 h at 37°C. The cells were then harvested, resuspended in PBS, and spread onto LB plates with the appropriate antibiotics to select the transconjugants. As a result, all plasmids were successfully transferred from *P. aeruginosa* PAO1808 to *P. aeruginosa* PAO1R in filter mating assays, confirming their self-transmissibility (Table S9). No transconjugants of Rms163 were detected in the liquid mating assay (Table S9). Based on the MOB-suite analysis, the T4SS family and MOB type of Rms139 (PInc-2), Rsu2 (PInc-9) and RP1-1 (PInc-11) could be predicted, whereas Rms163 (PInc-5) and R716 (PInc-12) harbored Dot/Icm secretion systems, which differ from the traditional type IV secretion systems (T4SS)^10^. These different conjugation systems and their features possibly reflect environmental conditions, indicating that each plasmid may have a preference for certain environmental conditions that make its transfer more efficient.

**Pangenome analyses of PInc group plasmids**

In *Pseudomonas* species (e.g., *Pseudomonas aeruginosa*) and Enterobacterales (e.g., *Escherichia coli*), incompatibility (Inc) tests have been performed, and some Inc groups defined in *Pseudomonas* correspond to those in Enterobacterales; for example, IncP-1, IncP-3, IncP-4, and IncP-6 in *Pseudomonas* correspond to IncP, A/C, IncQ, and IncG in Enterobacterales, respectively. Here, we describe pangenome analyses of *Pseudomonas* plasmids, highlighting core genes conserved across all members of each Inc group, as well as accessory genes variably present among group members.

***PInc-1 (=IncP-1=IncP)*** In our previous study, 14 subgroups within PInc-1, denoted as PInc-1αβγδεζηθικλμορ, were proposed based on incompatibility tests^11^. This study introduces a new subgroup, PInc-1π, consisting of 41 plasmids, including pEP5289 (Table S1). Although previously identified as an PInc-1 plasmid from *Neisseria**^12^*, its incompatibility with other PInc-1 plasmids remains untested. *Neisseria* is the main host of these plasmids, potentially distinguishing them from others in the PInc-1 group. A total of 25 core genes were identified across 321 plasmids (Table 1, S2, S3-1).

***PInc-2 (=IncP-2)*** Plasmids in the PInc-2 group are larger, with an average size of 440 kb. They contain 293 core genes (Table 1, S2, S3-2), which accounts for around 53% of the gene count (293/549.8) in 87 plasmids (Table S2). The megaplasmids pBT2436 and pRBL16, associated with ARGs^13,14^, are recognized within this group. Interestingly, one key gene, the Tus protein (Table S3-2), binds to replication terminus sites^15^. This gene is found in all PInc-2 plasmids and several large plasmids (113-527 kb), leading to its removal from the RIP reference sequences (<https://figshare.com/s/3e3ced8633136259f678>).

***PInc-3 (=IncP-3=IncA/C)*** The PInc-3 group, also known as IncA/C, was reclassified into IncA and IncC^16^. Despite conserved RIP genes (>98% identity), differences in iteron numbers may allow compatibility. PIRATE analysis revealed 48 core genes, but accessory genes exhibit diversity (Table S3-3). Most PInc-3 plasmids (96.7%) carry at least one ARG (Table 1, S2), making them important ARG vectors.

***PInc-4 (=IncP-4=IncQ)*** PInc-4 plasmids (IncQ group) have a wide DNA size range, from 1.7 to 412 kb (average 121 kb, Table 1, Figure 1A). They are defined by the *repBAC* genes for single-strand displacement replication^17^. Only two core genes (*repA* and *repC*=*repP-4*) were consistently identified, making up 1.5% of their gene count (Table 1, S2, S3-4). The small number of core genes may reflect the presence of multi-replicons; only 29.1% of the plasmids have a single replicon (Table S2).

***PInc-5 (=IncP-5)*** Only two PInc-5 plasmid, Rms163 and pPALA50, were identified (Table 1). ARGs in Rms163 were found within a Tn*3*-like transposon and a remnant class 1 integron (Figure S16), with 100% identity to segments of Tn*5393* and an IncP-9 plasmid from Japan^9^. The *repP-5A* gene in Rms163 shares 100% identity with pPALA50, a plasmid from a cystic fibrosis patient, although pPALA50 lacks ARGs (Figure S17)^18^.

***PInc-6 (=IncP-6=IncG)*** PInc-6 plasmids range from 12 to 497 kb (Figure 1A). Three core genes were identified, including *repP-6A*, *parC* (for plasmid partitioning)^19^, and a recombinase/transposase gene (Table S3-5). These core genes account for 5.7% of the gene content (Table S2). The archetype Rms149 is incompatible with IncU plasmids but shares r*epP-6A* and *parC* genes^20^, although their backbone genes differ^21^. This study treats PInc-6 (IncG) and IncU as separate groups.

***PInc-7 (=IncP-7)*** PInc-7 plasmids have 10 core genes (5.2% of the average of 191.9 genes per plasmid, Table S2), including *repA* (*repP-7A*) and *parWAB* genes involved in plasmid partitioning^22^ (Table S3-6). Some plasmids, like pWW53 and pND6-1, lack conjugative transfer genes, possibly explaining the small number of core genes^23^. Of the 23 plasmids, only four contain ARGs, while nine others carry catabolic genes.

***PInc-9 (=IncP-9)*** PInc-9 plasmids have 10 core genes (9.3% per 107.6 average gene numbers, Table S2), including *rep* (*repP-9A*) and genes for plasmid partition and single-strand DNA-binding proteins (*ssb*) (Table S3-7). They range from 38 to 483 kb, with smaller plasmids lacking conjugative transfer genes (Table S3-7). Larger plasmids, like pKF715A, contain *bph*-*sal* element^24^, highlighting the diversity within IncP-9 plasmids.

***PInc-10 (=IncP-10)*** All PInc-10 plasmids were found in *P. aeruginosa*, with over 50% carrying carbapenem resistance genes (Table 1). These plasmids range from 29,402 to 106,679 bp, with pangenome analysis identifying 209 gene families and 19 core genes (29.0% of average gene numbers) involved in plasmid replication and maintenance (Table S3-8).

***PInc-11 (=IncP-11)*** PInc-11 plasmids, also found in *P. aeruginosa*, share 37 core genes, including *repP-11A* and plasmid partition genes (Table S3-9). Several of these plasmids were found in clinical isolates, indicating their role in spreading ARGs. Importantly, pOXA-198 (GenBank accession number MG958650), previously proposed as an “IncP-11 plasmid”^25^, lacks any genes resembling the *repP-11A*, suggesting that it may not belong to the traditionally recognised IncP-11 group^26^.

***PInc-12 (=IncP-12)*** Nine PInc-12 plasmids, hosted by *P. aeruginosa*, share 166 core genes (165 PIRATE-defined core genes plus the manually verified *repP-12A* locus, 74.6% of their average gene number, Table S2, S3-10). While these plasmids show high genetic conservation, only R716 contains known ARGs.

***PInc-15 (=PromA)*** Forty-two PInc-15 plasmids were identified, sharing 30 core genes (50.3% of their average gene count). These genes are involved in replication and conjugative transfer (Table S3-11). Despite their conserved structure, many of these plasmids lack known accessory genes^27,28^.

***PInc-16 (pSN1216-29-like plasmids)*** Twenty-four PInc-16 plasmids range in size from 35 to 174 kb and share 17 core genes (28.9% of their gene count), including *repA* and genes for conjugative transfer (Table S3-12). Some plasmids, like pCTX-M15, are multi-replicons, reducing shared gene numbers. Six PInc-16 plasmids carry ARGs.

***PInc-17 (pQBR103-like plasmids)*** Similar to PInc-2, PInc-17 plasmids are large (515 kb) with 229 core genes (Table S3, S3-13). These plasmids are important for their association with ARGs, including carbapenem and tigecycline resistance^29,30^***.***

***PInc-18 (pSTY-like plasmids)*** PInc-18 plasmids show significant genetic diversity, with 1,073 gene families across 15 plasmids (Table S3-14). Some carry clinically relevant ARGs, making them targets for further research^30,31^.

**Phylogenetic analysis of RIPs**

To identify the evolutionary relationships of the RIPs of PInc groups, 2,397 nucleotide sequences of putative RIPs were detected from DFAST-core predicted ORFs of PInc plasmids (n = 2,353) included in the PLSDB and two additional plasmids (pOZ176 and RK2). Detection was based on nucleotide similarity and rep_cluster of MOB-typer v3.1.8. Nucleotide sequences of the putative RIPs were translated using EMBOSS transeq v6.6.0^32^ using the standard prokaryotic genetic code (table 11), and 23 sequences were excluded because they had stop codons in the middle, possibly due to pseudogenization or misassembly of the plasmid sequences. Consequently, 2,374 RIP sequences were obtained from 2,332 *Pseudomonas* plasmids. A non-redundant protein set at 100% sequence identity was extracted using MMseqs2 cluster v15-6f452 (with options “--cluster-mode 2 --cov-mode 0 --min-seq-id 1.0 -c 0.5”)^33^, and the longest sequence was extracted as a representative for each cluster. The clustering identified 251 non-redundant proteins. Sequence homology between these non-redundant sequences was investigated using BLASTp^34^. The resultant hits were visualised using CPAP v0.3.0 (https://github.com/yosuken/CPAP). To detect remote homology between PInc groups, we extended the sequence diversity by collecting homologous sequences through two steps of homology search processes as described below. The detailed scheme for homolog collection and phylogenetic tree reconstruction is shown in Figure S4.

First, we clustered the 251 non-redundant RIP sequences at 20% identity using MMseqs2^33^ cluster with the options “--cluster-mode 2 --cov-mode 0 --min-seq-id 0.2 -c 0.5”, which yielded 12 clusters. These clusters largely corresponded to the PInc groups, except that some PInc groups were merged into a single protein cluster (i.e., PInc-2/PInc-3/PInc-17 and PInc-10/PInc-11). We therefore designated these clusters as G-PInc groups, following the PInc numbering system (e.g., G-PInc-1, G-PInc-3, and G-PInc-4; Figure S4C). Sequence alignments for the 12 groups were generated using MAFFT^35^ v7.455 with the options “--genafpair --maxiterate 1000”, and corresponding HMM profiles were built using hmmbuild in HMMER^36^ v3.3.2. Homologous RIP sequences were then searched against translated ORFs from PLSDB plasmids predicted by OrfM^37^ (n = 20,331,985), using OrfM with the options “-c 11 -m 270”, which specify codon table 11 and a minimum ORF length of 270 bp. We used OrfM to maximize coverage of the possible ORF space. Hits were retained when the E-value was <1e-2 and the aligned HMM length exceeded 60% of the full HMM length. When a hit was split into multiple alignments, the aligned regions were merged if their order was concordant, using parse_hmmsearch.rb v0.1.0 with the options “--max-hmm-ovp-frc 1 --max-hmm-ovp-len 60 --max-ali-ovp-frc 1 --max-ali-ovp-len 60 -e 1e+6”, which allow overlaps of up to 60 amino acids. This search yielded 15,049 RIPs. In parallel, eight additional known RIPs outside the PInc groups were used as queries for diamond blastp searches against the same OrfM-predicted PLSDB ORFs, and significant hits were retained when the E-value was <1e-4 and the aligned region covered at least 50% of the query length. This search yielded 11,179 RIPs in eight groups according to the best-hit query. After pooling the original sequences, the hmmsearch-derived RIPs, and the diamond-derived RIPs, protein clustering was performed again using MMseqs2 cluster with the options “--cluster-mode 2 --cov-mode 0 --min-seq-id 0.9 -c 0.5” to generate a non-redundant protein set at 90% sequence identity. Representative sequences were selected for each cluster by prioritizing sequences used in the original profile alignments over newly retrieved sequences, and by prioritizing longer sequences when candidates originated from the same source. This procedure yielded 1,885 non-redundant sequences. The non-redundant sequences were subsequently classified according to the HMM hits detected by hmmsearch, except that G-PInc-12 was merged into G-PInc-1 because 95% of G-PInc-1 sequences and 99% of G-PInc-12 sequences belonged to clusters containing members of both groups in the clustering step above, indicating that the two groups were highly similar. Consequently, the 1,885 non-redundant sequences were classified into 11 groups (Figure S4C).

Second, sequence alignments for the 11 groups and the eight groups derived from the additional RIP queries were generated using MAFFT with the same options, and the corresponding 19 HMM profiles were built using hmmbuild. Homologous RIP sequences were then searched against UniParc protein sequences (downloaded from the FTP site in February 2024; n = 607,912,929) using hmmsearch with the option “-Z 1000000”. Significant hits were retained when the E-value was <1e-5 and the aligned HMM length exceeded 60% of the full HMM length, yielding 134,212 RIPs. These sequences were pooled with the original sequences used in the profile alignments (n = 1,885), giving 146,076 sequences in total. Potential contamination by unrelated proteins, including transcription factors, was removed by hmmsearch against the Pfam database; sequences were discarded when a non-replication-related Pfam hit had a smaller E-value than the best RIP-associated hit. This filtering step retained 141,276 RIPs. Protein clustering was then performed using MMseqs2 cluster with the options “--cluster-mode 2 --cov-mode 0 --min-seq-id 0.6 -c 0.5”, and representative sequences were selected as described above, yielding 9,788 representatives. After alignment of each group using MAFFT and removal of positions with >70% gaps and sequences with >50% gaps, 19 HMMs were constructed using hmmbuild. These HMMs were then used in hmmsearch against the PLSDB ORFs. Significant hits were retained when the E-value was <1e-5 and the aligned HMM length exceeded 60% of the full HMM length, and potential contamination by unrelated proteins was again removed by hmmsearch against the Pfam database using the same criterion as above, yielding 53,391 RIPs. Pooling these sequences with the representatives yielded 63,179 RIPs in 19 groups according to the best-hit HMM. These 19 groups were then merged into 14 homologous groups based on UniParc sequence overlap, and representative sequences were selected by 50% identity clustering within each group, yielding 5,488 representative RIPs in 14 groups (Figure S4A–C). Sequence alignments for the 14 groups were then recalculated using MAFFT with the same options. The resulting alignments were trimmed by removing positions with more than 70% gaps. Sequences containing gaps in more than 50% of the positions in the trimmed alignments were subsequently removed, leaving 5,345 representative RIPs in 14 groups. HMM profile databases were then built from the filtered alignments using hhmake in HH-suite v3.3.0^38^.

To investigate whether these groups exhibit homology with one another, a highly sensitive HMM–HMM comparison was performed using HHblits with the options “-n 2 -mact 0.2 -z 20 -b 20”. Based on these comparisons, G-PInc-6 showed no detectable homology to the other groups and was excluded from subsequent extraction of the shared homologous region. Homologous regions were therefore extracted from the alignments of the remaining 13 groups, containing 4,793 sequences in total. For the sWH supergroup, regions of G-PInc-9 homologous to dWH groups were first identified, their union was taken, and a margin of 5 amino acids was added on both sides, yielding positions 1–105 of G-PInc-9 as the reference homologous region. Corresponding homologous regions were then extracted from the remaining sWH groups. For the dWH supergroup, regions of G-PInc-1 homologous to sWH groups were identified in the same manner, yielding positions 141–256 of G-PInc-1 as the reference homologous region, and corresponding regions were then extracted from the remaining dWH groups (Figure S5B). Sequences were discarded when the extracted region contained more than 60% gaps in the alignment. In total, this procedure removed 10 sequences.

Subsequently, we realigned the extracted homologous regions of the remaining 13 groups using MAFFT with the same options. Sequences containing more than 60% gaps in the extracted homologous region were discarded, removing 10 sequences. The resulting alignments contained 4,783 representative sequences in 13 groups. After removing positions with more than 50% gaps, the final alignment comprised 150 positions and was used for maximum-likelihood phylogenetic reconstruction.

**Definition and validation of winged-helix (WH) domains in RIPs**

To establish a unified framework for identifying winged-helix (WH) domains in plasmid replication initiation proteins (RIPs), we systematically refined domain boundaries for both double-WH (dWH) and single-WH (sWH) architectures. For RIPs containing tandem WH domains (dWH type), previously reported WH1 (194-309 aa in CAJ85684, 116 amino acids) and WH2 (310-382 aa in CAJ85684, 73 amino acids) regions for the RIP of plasmid RK2/RP4^39^ were used as an initial reference. In the present study, we redefined these boundaries to improve structural and evolutionary consistency. Specifically, the WH1 region was shortened to 97 amino acids (194-290 aa), and a discrete linker segment was introduced between WH1 and WH2 to clearly separate the two domains. Notably, structural analysis of the F plasmid RIP protein, RepE with double-WH regions^40^, delineates the WH1-equivalent region as a structurally compact segment shorter than that previously defined in RK2/RP4^39^. This structural evidence further supports our refined definition of WH1 as a more compact and structurally coherent domain. Multiple sequence alignment across representative dWH-type RIPs confirmed the validity and conservation of these refined boundaries. In addition, experimentally resolved crystal structures of dWH-type RIPs, including RepA of plasmid pPS10 (PDB: 1HKQ), were examined to verify that the refined WH1 and WH2 regions correspond to the structurally defined winged-helix folds.

**Before**

**TrfA44_WH1 (116 aa: 194-309/382)**

**RADDDELVWQQVLEYAKRTPIGEPITFTFYELCQDLGWSINGRYYTKAEECLSRLQATAMGFTSDRVGHLESVSLLHRFRVLDRGKKTSRCQVLIDEEIVVLFAGDHYTKFIWEKY**

**TrfA44_WH2 (73 aa: 310-382/382)**

**RKLSPTARRMFDYFSSHREPYPLKLETFRLMCGSDSTRVKKWREQVGEACEELRGSGLVEHAWVNDDLVHCKR**

**After**

**TrfA44_WH1 (97 aa: 194-290/382)**

**RADDDELVWQQVLEYAKRTPIGEPITFTFYELCQDLGWSINGRYYTKAEECLSRLQATAMGFTSDRVGHLESVSLLHRFRVLDRGKKTSRCQVLIDE**

**TrfA44_WH2 (73 aa: 310-382/382)**

**RKLSPTARRMFDYFSSHREPYPLKLETFRLMCGSDSTRVKKWREQVGEACEELRGSGLVEHAWVNDDLVHCKR**

Next, for RIPs containing a single WH domain (sWH type), we used structural and functional data of the RIP of pSK41 reported previously^41^ as a reference. Based on these data, we refined the conserved WH core. Based on these data, we refined the WH domain to an 83-amino-acid core region (30-112 aa in AAC61944). Multiple sequence alignment across representative sWH-type RIPs supported the validity and conservation of this boundary definition. Furthermore, the crystal structure of the sWH-type RIP from plasmid pTZ2162 (PDB: 4PT7) was used to confirm that the defined region coincides with the structurally resolved WH fold.

**Before**

**pSK41_WH (107 aa: 12-118/319)**

**YKERFYQLPKVFFTNPNYKDLSNDAKIAYAILRDRLQLSIKNNWIDTEGNIYFIYTVADLEVILNCGNKKITKIKKELENVDLLIQKRQGLNKPNLLYLLKPAITKN**

**After**

**pSK41_WH (83 aa: 30-112/319)**

**KDLSNDAKIAYAILRDRLQLSIKNNWIDTEGNIYFIYTVADLEVILNCGNKKITKIKKELENVDLLIQKRQGLNKPNLLYLLK**

Using these standardized criteria, we annotated the WH domain regions in all RIPs from PInc-1 through PInc-18, except PInc-6, which belongs to the AEP superfamily and does not contain a WH domain. For dWH-type RIPs, WH1 and WH2 are indicated in magenta and cyan, respectively. For sWH-type RIPs, the single WH domain is shown in cyan. The gray regions indicate sequences identified as Pfam domains.

**WH**

**Pnc-1alpha_RP4　(WH1 97 aa: 194-290/382; WH2 73 aa: 310-382/382)**

**MNRTFDRKAYRQELIDAGFSAEDAETIASRTVMRAPRETFQSVGSMVQQATAKIERDSVQLAPPALPAPSAAVERSRRLEQEAAGLAKSMTIDTRGTMTTKKRKTAGEDLAKQVSEAKQAALLKHTKQQIKEMQLSLFDIAPWPDTMRAMPNDTARSALFTTRNKKIPREALQNKVIFHVNKDVKITYTGVELRADDDELVWQQVLEYAKRTPIGEPITFTFYELCQDLGWSINGRYYTKAEECLSRLQATAMGFTSDRVGHLESVSLLHRFRVLDRGKKTSRCQVLIDEEIVVLFAGDHYTKFIWEKYRKLSPTARRMFDYFSSHREPYPLKLETFRLMCGSDSTRVKKWREQVGEACEELRGSGLVEHAWVNDDLVHCKR**

**PInc-2_Rms139　(WH1 119 aa: 138-256/395; WH2 92 aa: 276-367/395)**

**MDVIESQNDLLTGLDCNSQSLSRKETGVVARLNSIRSKATGKRIQEPAASSTLPLFSAYQITLDDNDLGNLVSIYDALPRFAWSGKTVRSVAEMTIVKEGTINGEPFRVILKAVPMQKRLKVDGKLSKQVEDVGVFPGAREEFVEEALRKFSTQGNAKFNEAECSVQFTLYELQKELQSQKHTYTYAELREALEILSEAPLTIQTRTADGEVIDIKSTYLPFLAIRSRNKRSSAYVQPSPDDDPESSVLCKAVLHPLISRSIANGDYRLYQYTTSMGLTNGIARVLYRMLSFKWRNASPSHPYTFSLVDFLSNTARGLSNRMPEDFRAMNIALEQLVKEKAIQRYEHTKIAKSKGKGAKDYTYKLWPTNEFVSTIIKGHQAEARREIQLMAKRAR**

**PInc-3_pRA1 (WH1 106 aa: 111-216/366; WH2 90 aa: 236-325/366)**

**MDHQLESINGTIMSKRTKDKDLEKLDVIKDSPQMSLFEIIESPAKKDDYSNTIEIYDALPKYIWDQKREHEDLSNAVVTRQCSIRGQQFTVKVKPAIIEKDDGRTVLIYAGQREEILEDALRKLAVNGKGHIIEGKAGVMFTLYELQKELSKMGHGYNLTEIKEAIQVCRGATLECISDDGEAFISSSFFPMVGLTTRGEFRKKGGNARCYVQFNPLVNESIMNLSFRQYNYKIGMQIRSPLARYIYKRMSHYWTQASPDSPYTPSLISFLTQSPRELSPRMPENVRAMKLALEALIKQEVISDYDANQIKDGRRVIDVRYVIRPHENFVKQVMASNKRKQQTELRAIKHGTIDHDIIDERQSKGR**

**PInc-4_pRSF1010 (WH1 120 aa: 60-179/283; WH2 79 aa: 200-278/283)**

**MVKPKNKHSLSHVRHDPAHCLAPGLFRALKRGERKRSKLDVTYDYGDGKRIEFSGPEPLGADDLRILQGLVAMAGPNGLVLGPEPKTEGGRQLRLFLEPKWEAVTAECHVVKGSYRALAKEIGAEVDSGGALKHIQDCIERLWKVSIIAQNGRKRQGFRLLSEYASDEADGRLYVALNPLIAQAVMGGGQHVRISMDEVRALDSETARLLHQRLCGWIDPGKTGKASIDTLCGYVWPSEASGSTMRKRRQRVREALPELVALGWTVTEFAAGKYDITRPKAAG**

**PInc-5_Rms163 (WH 88 aa: 58-145/235)**

**MVKIVEFQERAAERLEGFRSNAKALPSLVYARYLLAPTSGVAKRIRGMDFALRKNRQHIMHAQHRATCIDVLNLLISRMDIRSRQCLYVNPRFGIRRNIYVPEIARLLKVCERTVTRALGSLERAGYLLRTATAKGMKMFLSLKLLRELNISLMYERLSNQLKGLDKKAQYEASNKGKKPSAPKQPGQGPSTPHTPAEQPVTNPGAPKERTEESLNIGNNFLAQLRGRRKRPPPG**

**PInc-7_pCAR1 (WH1 91 aa: 31-121/288; WH2 86 aa: 143-228/288)**

**MSNLPKPAKPEPKPLRVTKSNTLITASYRLTLNEQRLILAAISKLDPRRPMPKKVSVSAVDYSDIYGVQLRHAYEQMKVAADELYERDIKTFDGSGMERKRWVDRAKYLDGEGRVELSFTIHVMPYLTMLYSKVTSYDLRRVACLDSSHSFRLFEMLMQFRKTGWAYIEVESLRVALGLSDAYQRFNNLRQRVIDPAVAELKTKSNLDVSYELRREGRKVIAIKFTFCDLAQLPLNLELDPEDLPPLPEYDPAEESEWLITKPLGELAEAQEAGLLFAVSAEEGEQPD**

**PInc-9_pWW0 (WH 73 aa: 36-108/184)**

**MASDNNEIRAYAQPAQRGTWVQTERAGHEAWAALTAQAPRAAQLMHILVQHMDKQGALIISQATLAKLMDTSVANHKTRYRHSYKHNWIQTISVGGQRGGTLAYVVNSRIAWADKRDNLQFALFNARVLVSTEDQADLGDAKLKQLPTMADGDIQLPAGPGMDPPAQESLEGLLPDMPSIPHNS**

**PInc-10_pPAB546 (WH 94 aa: 76-169/286)**

**MGRTESVARPYVQRTNHGGNFCGHQPDAPRLDLIRPTTDKNRPGILRQLQERVRRYYRSPTVIPDLRNANRSRKGRQQRSERREACLLLLTSIIEHTDLVSLRCGVPTSGGFLSLTLDYLTQWTGLHPRRAERAMADLKRANLLTVSQPRQLNEDGSWRGLAAVKAVSKHLFAAFGLGRRLGYERDRASKRLAKKIAVRGGTLTGWARNVALLVGGLGKKPGRAFRSPASAGPTGVSHGMDSATYASARMQLLIDLMTRHPGMPRDDLYAEAERILQERLTKSIRA**

**PInc-11_RP1-1 (WH 94 aa: 71-164/279)**

**MSINTRTYVPRVNNGGNYCGHKPDAPRLGLVKPTTLKNRPKILQRLQEELRQYYYSPSRLPSLNAANRSKRQQRSERREACLLVLAAVLEYTDLTSLRCGVPSAEGFQSLTFQFLAEYIGIGMRRIERAVADIKFANILTVSQPRQLQEDGSYRGLAAIKSVNSLLFGAFGLLKWLKHERKRASERLAKKAKRQGGNLGQWSRSALAIGKMLVIRRRGPGDLPRIGPSHGGAGPPQPDAQTYGHMLNELILALKQEDPSRDGATCRKLAEAIITKKLAG**

**PInc-12_R716 (WH1 93 aa: 132-224/353; WH2 75 aa: 243-317/353)**

**MPLTGERKITELVYSSLCQCDNFAKIQKRQAPGHMSDKPSTSRLAERIRQVQEKNTRKGKSKPSTDVASSDAQSGTPLQLPFWPEATRGVPSGFLRSALFAGIQSKDRRFLKDEVIEAVGGVEIRFRGEQLDQTDLDVWEAVLHLARINQLSTKDRLRFNAHGMLKMLDRSTGGDQHDWLKTTLLRLTGAILDIKVGTQAYFGPMLEGGTRDEATDEYEIWVNPKIRALYEAGWTQINNDQRNALRRKPLAQWLHGWYSSHAAPYPISVETFYKLSGSTNKQMAGYKRLLMQAHDDLVAIGVLTSWEIKNRLVAVSKPQTPSQKRFLERKSQRADDGPEGGRQAEFIIEHQPE**

**PInc-15alpha_pSB102 (WH1 107 aa: 259-365/484; WH2 84 aa: 384-467/484)**

**MTKQQTTTGDRTRKKAAADVFDEIAYRDELVKIGIDLEAATLVASKTAQAHRDNAKARKQQQTTVGDIITNNNIPGRLREHSDEWKRHEPAAKAMAEERAKIAAARAKQIDIAGDFSIDLSIAPNFPLLARVQAEARARAAELAGRPAAKAKKPVKLFTTPDDKPAPSAAVARRRVETPIKAVYDYRYSELPERVRDLIEAHLAIEAEDAKSAGTLGFMTRALAIATLPHRKLAEDRFVRKNGDFTLTMLTAHPEGLPYGTLPRLLLTWVATEAVQKKERVLSLGNTLASYLNELGLHNTGGKRGDITRLKHAMTTLFSAIISCRYEGRDSWALQNVLLADKVEWWQPQDAAEAGAWQSRLQLSEPFFQECLDHPLPVDMRAMKVLRQSPLALDIYVWLTHRMSYLSKRTTIPWVSLSGQFGAGYAVNDQGLRDFKRAFLRELKNVVAIYPEAKLSESRNGLVLYPSPTHVLPDNSPKQPSLPF**

**PInc-16_pSN1216-29　(WH1 116 aa: 171-286/413; WH2 77 aa: 308-384/413)**

**MNGPKPISDIVGGNLLARLEQTHQRYREKVEAAALPGETFEQAEKRLREEGEREQRRKSNEVAQFQAEATLAAQLRDRSPIPRSPIPNTSSPSGQTKRNNPASMRRPPESDRQPDFFVPGLYDVATKDNRSLMDVALFRLSKRDKRAGEVIRYDLPDGFVEVKAGPDGMASIWDYDIVLMLISHLTESMNLFNAGRGAMPGKKFVPHASDIAKFCRRGDGGRQADELEAALDRLLGTTIKSVRETPSRNGKRVVRETEAESLIGPYRVVSRTDTGKVASVEIEAPNWIYREVTGGKQPDVLTVHPDYFLIEPGLGRFVYRLARRAAGKNTAKWAFKTIYERSGSAGTFKEFCRMMRRLIDANDLPEYDLREEDGQSGPLLLITRRTDAVELAADDEPEPDGDEQENGKNAPGV**

**PInc-17_pQBR103　(WH1 120 aa: 122-241/376; WH2 92 aa: 261-352/376)**

**MENVSASAGRPDNSIAAKLRNMQHRNMQRSAGPDLQGELFNPYEIRLDDADHSNLVSIYDALPRFSWASKTVRDMSQMSTQLEGSLNGQPFRVVMKAVSMEIPIKIKGKATGQTEWVGLYPSSREECVEEALRKFSTQPGRFNEVKCSVSFSLYDLRQELKSVGHTYTYDELRQALDILAKSSLTIQTRSSDGDVIDITSNYLPFLYLRSKNKKSKNYIDVKDISDQEKGDATLCMAVLHPLISRGIERGDYRLYHYSTSMKLTNSLAKTLNRELSMRWRNASPTHPYRFHLVEFLSNTARGLSKRMPEDYRAVNIALAELVQEGVLSEFKSTPIKKPKGKGAIDYVYEILPTKKFVGMMIDGHRRENNRQLALTA**

**PInc-18_pSTY　(WH1 98 aa: 11-108/431; WH2 91 aa: 129-219/431)**

**MRNVSAASDFTFLQRKLYNTLLQFAQQRPREEMVHEIPIKQVEDNIGHTTSNSRDYLKKVLVSMSQTQVEFDYKGESPGRKSEWGIANLIAEAYILEDGQTLRFSFPPDLKRRLLDPAIFNLIDLRMQYHFSSFSALTLHEITSRYLGSPMGETYRAHWSEWSVVLSGSATPHAEFRDFNKMLGRAIDQVNSIERRFRISPHVTKLNRKMDKLWFKLETLIQPGLDLGPSPELVSQDVSKRLKALSLSQKDIDELGMTHDEEYLLAQADYTEAQMRKEGANVASPAAYFKAAVANNYAKAPTQQKAAEPGRGKKPAASKTGEKSKPAQAPSQAPKAAPSNQMADLLEQWGAAQREAIRAQFMELSDEQKKELAEKYETELRKDDLAYSQYRSKGLNTMVINCLVAIQFQERFPETPSSETLLQFLLGGAKI**

Following phylogenetic classification into eight major clades (A–H), representative RIPs were selected to illustrate the relationship between WH domains and other conserved Pfam-defined regions, including RepL (PF05732), Bac_RepA_C (PF18008), IncFII_RepA (PF02387), RepA_N (PF06970), RepA_C (PF04796), RPA (PF10134), RepC (PF06504), Rep3_N (PF01051), and Rep3_C (PF21205). Structural modeling further supported the evolutionary correspondence of the defined WH regions across clades.

**WH**

**A_pSN2 (WH 67 aa: 52-118/158)**

**MKERYGTVYKGSQRLIDEESGEVIEVDKLYRKQTSGNFVKAYIVQLISMLDMIGGKKLKIVNYILDNVHLSNNTMIATVREIAEGTNTSTKTVNTTLKILEEGNIIKRRTGALMLNPELLMRGDDQKQKYLLLEFGNFEQEDDQKQENALSEYYSFKE**

**B_p140355 (WH 84 aa: 27-110/341)**

**MPKSTTAYRSVRKSHFTQISNDLLNDKTISLEAKGLLSIFLSNNDEWDLHMSEIIKRSKNGRDAHYTALKKLIKAGYIARLEFKQLSNFQFLHLEYIFSDNKEDVINGIKDAQKFANENEQIIVLTYKDIEGRKIESSIENTEKKPFTENPDTEENSENSPLTENPDTDNPNTENANTENQYNNNTNSNNTNINNTNSNNTNDMNDKKQNTEKNHSYHSNHFSNTHDKESLKYIELQELPELSKSYINNFSYEEVKSIKSVILKAKKSFNNKYDTFYMLEDIDEELLLVLKRFKGYLVKKQEKVANMEGYLMRSIIAELEEMHSTIMRRKNMENNPLSLFN**

**C_pOXA-48　(WH 90 aa: 122-211/355)**

**MQGLVNQKNPYLQLSDIESVEGLSPEFISWLESQSPKESPLQLLLDEGTKPVRKTRRRRGTHSTACLCPEPSWYRPDNFKKLPGQLGHAYNRLVRRDRKTGALSLRMRISRHPYFVQLREQAGRKRDFRPEREKLLDAIVPLLVSTVDRATHIDTINLSKMAWQLSEKDSEGNVIRKVTVPRVCRLLQHMMEFGLLALPEGVTWDPFNRKWFPKHVVLTERLWKMIGVDLDKLYAEQAEQVAAEAAGWITRDEHGQAEEISVKAARRRWYEKMMHATLVRRREAALKGKREKKLKALAEQPYDDRAYAMSVHLIRTLPKDELHALSPEQFTQRVRSHLYQLDLGLERESGPPDYH**

**D_pSK41　(WH 83 aa: 30-112/319)**

**MSKQFFTVEENYKERFYQLPKVFFTNPNYKDLSNDAKIAYAILRDRLQLSIKNNWIDTEGNIYFIYTVADLEVILNCGNKKITKIKKELENVDLLIQKRQGLNKPNLLYLLKPAITKNDIYEIDKAENEVEALQDKEVSKGHVQKCQKDTSRNVKRTRLEMSKGHTNDTDFIDTDFIDTESNDMNNMNDTNQHSNHSNHFSNIHDKESLKYIELQELPELIKSYINNFSYEEVKSIKSVILKAKKSFNNKYDTFYMLEDIDEELLLVLKRFKGYLVKKQEKVANMEGYLMRSIIAELEEMHSTIMRRKNMENNPLSLFN**

**E_pSN1216-29 (WH1 116 aa: 171-286/413; WH2 77 aa: 308-384/413)**

**MNGPKPISDIVGGNLLARLEQTHQRYREKVEAAALPGETFEQAEKRLREEGEREQRRKSNEVAQFQAEATLAAQLRDRSPIPRSPIPNTSSPSGQTKRNNPASMRRPPESDRQPDFFVPGLYDVATKDNRSLMDVALFRLSKRDKRAGEVIRYDLPDGFVEVKAGPDGMASIWDYDIVLMLISHLTESMNLFNAGRGAMPGKKFVPHASDIAKFCRRGDGGRQADELEAALDRLLGTTIKSVRETPSRNGKRVVRETEAESLIGPYRVVSRTDTGKVASVEIEAPNWIYREVTGGKQPDVLTVHPDYFLIEPGLGRFVYRLARRAAGKNTAKWAFKTIYERSGSAGTFKEFCRMMRRLIDANDLPEYDLREEDGQSGPLLLITRRTDAVELAADDEPEPDGDEQENGKNAPGV**

**F_pRSF1010 (WH1 120 aa: 60-179/283; WH2 79 aa: 200-278/283)**

**MVKPKNKHSLSHVRHDPAHCLAPGLFRALKRGERKRSKLDVTYDYGDGKRIEFSGPEPLGADDLRILQGLVAMAGPNGLVLGPEPKTEGGRQLRLFLEPKWEAVTAECHVVKGSYRALAKEIGAEVDSGGALKHIQDCIERLWKVSIIAQNGRKRQGFRLLSEYASDEADGRLYVALNPLIAQAVMGGGQHVRISMDEVRALDSETARLLHQRLCGWIDPGKTGKASIDTLCGYVWPSEASGSTMRKRRQRVREALPELVALGWTVTEFAAGKYDITRPKAAG**

**G_pSB102 (WH1 107 aa: 259-365/484; WH2 84 aa: 384-467/484)**

**MTKQQTTTGDRTRKKAAADVFDEIAYRDELVKIGIDLEAATLVASKTAQAHRDNAKARKQQQTTVGDIITNNNIPGRLREHSDEWKRHEPAAKAMAEERAKIAAARAKQIDIAGDFSIDLSIAPNFPLLARVQAEARARAAELAGRPAAKAKKPVKLFTTPDDKPAPSAAVARRRVETPIKAVYDYRYSELPERVRDLIEAHLAIEAEDAKSAGTLGFMTRALAIATLPHRKLAEDRFVRKNGDFTLTMLTAHPEGLPYGTLPRLLLTWVATEAVQKKERVLSLGNTLASYLNELGLHNTGGKRGDITRLKHAMTTLFSAIISCRYEGRDSWALQNVLLADKVEWWQPQDAAEAGAWQSRLQLSEPFFQECLDHPLPVDMRAMKVLRQSPLALDIYVWLTHRMSYLSKRTTIPWVSLSGQFGAGYAVNDQGLRDFKRAFLRELKNVVAIYPEAKLSESRNGLVLYPSPTHVLPDNSPKQPSLPF**

**H_pCAR1 (WH1 91 aa: 31-121/288; WH2 86 aa: 143-228/288)**

**MSNLPKPAKPEPKPLRVTKSNTLITASYRLTLNEQRLILAAISKLDPRRPMPKKVSVSAVDYSDIYGVQLRHAYEQMKVAADELYERDIKTFDGSGMERKRWVDRAKYLDGEGRVELSFTIHVMPYLTMLYSKVTSYDLRRVACLDSSHSFRLFEMLMQFRKTGWAYIEVESLRVALGLSDAYQRFNNLRQRVIDPAVAELKTKSNLDVSYELRREGRKVIAIKFTFCDLAQLPLNLELDPEDLPPLPEYDPAEESEWLITKPLGELAEAQEAGLLFAVSAEEGEQPD**

**Pfam
A_pSN2 (RepL 148 aa: 1-148)
MKERYGTVYKGSQRLIDEESGEVIEVDKLYRKQTSGNFVKAYIVQLISMLDMIGGKKLKIVNYILDNVHLSNNTMIATVREIAEGTNTSTKTVNTTLKILEEGNIIKRRTGALMLNPELLMRGDDQKQKYLLLEFGNFEQEDDQKQENALSEYYSFKE**

**B_p140355 (Bac_RepA_C 56 aa: 248-303)**

**MPKSTTAYRSVRKSHFTQISNDLLNDKTISLEAKGLLSIFLSNNDEWDLHMSEIIKRSKNGRDAHYTALKKLIKAGYIARLEFKQLSNFQFLHLEYIFSDNKEDVINGIKDAQKFANENEQIIVLTYKDIEGRKIESSIENTEKKPFTENPDTEENSENSPLTENPDTDNPNTENANTENQYNNNTNSNNTNINNTNSNNTNDMNDKKQNTEKNHSYHSNHFSNTHDKESLKYIELQELPELSKSYINNFSYEEVKSIKSVILKAKKSFNNKYDTFYMLEDIDEELLLVLKRFKGYLVKKQEKVANMEGYLMRSIIAELEEMHSTIMRRKNMENNPLSLFN**

**C_pOXA-48 (IncFII_repA 255 aa: 37-291)**

**MQGLVNQKNPYLQLSDIESVEGLSPEFISWLESQSPKESPLQLLLDEGTKPVRKTRRRRGTHSTACLCPEPSWYRPDNFKKLPGQLGHAYNRLVRRDRKTGALSLRMRISRHPYFVQLREQAGRKRDFRPEREKLLDAIVPLLVSTVDRATHIDTINLSKMAWQLSEKDSEGNVIRKVTVPRVCRLLQHMMEFGLLALPEGVTWDPFNRKWFPKHVVLTERLWKMIGVDLDKLYAEQAEQVAAEAAGWITRDEHGQAEEISVKAARRRWYEKMMHATLVRRREAALKGKREKKLKALAEQPYDDRAYAMSVHLIRTLPKDELHALSPEQFTQRVRSHLYQLDLGLERESGPPDYH**

**D_pSK41 (RepA_N 75 aa: 15-89; Bac_RepA_C 83 aa: 219-301)**

**MSKQFFTVEENYKERFYQLPKVFFTNPNYKDLSNDAKIAYAILRDRLQLSIKNNWIDTEGNIYFIYTVADLEVILNCGNKKITKIKKELENVDLLIQKRQGLNKPNLLYLLKPAITKNDIYEIDKAENEVEALQDKEVSKGHVQKCQKDTSRNVKRTRLEMSKGHTNDTDFIDTDFIDTESNDMNNMNDTNQHSNHSNHFSNIHDKESLKYIELQELPELIKSYINNFSYEEVKSIKSVILKAKKSFNNKYDTFYMLEDIDEELLLVLKRFKGYLVKKQEKVANMEGYLMRSIIAELEEMHSTIMRRKNMENNPLSLFN**

**E_pSN1216-29 (RPA 255 aa: 27-281)**

**MNGPKPISDIVGGNLLARLEQTHQRYREKVEAAALPGETFEQAEKRLREEGEREQRRKSNEVAQFQAEATLAAQLRDRSPIPRSPIPNTSSPSGQTKRNNPASMRRPPESDRQPDFFVPGLYDVATKDNRSLMDVALFRLSKRDKRAGEVIRYDLPDGFVEVKAGPDGMASIWDYDIVLMLISHLTESMNLFNAGRGAMPGKKFVPHASDIAKFCRRGDGGRQADELEAALDRLLGTTIKSVRETPSRNGKRVVRETEAESLIGPYRVVSRTDTGKVASVEIEAPNWIYREVTGGKQPDVLTVHPDYFLIEPGLGRFVYRLARRAAGKNTAKWAFKTIYERSGSAGTFKEFCRMMRRLIDANDLPEYDLREEDGQSGPLLLITRRTDAVELAADDEPEPDGDEQENGKNAPGV**

**F_pRSF1010 (RepC 255 aa: 7-261)**

**MVKPKNKHSLSHVRHDPAHCLAPGLFRALKRGERKRSKLDVTYDYGDGKRIEFSGPEPLGADDLRILQGLVAMAGPNGLVLGPEPKTEGGRQLRLFLEPKWEAVTAECHVVKGSYRALAKEIGAEVDSGGALKHIQDCIERLWKVSIIAQNGRKRQGFRLLSEYASDEADGRLYVALNPLIAQAVMGGGQHVRISMDEVRALDSETARLLHQRLCGWIDPGKTGKASIDTLCGYVWPSEASGSTMRKRRQRVREALPELVALGWTVTEFAAGKYDITRPKAAG**

**G_pSB102 (RepA_C 153 aa: 258-410)**

**MTKQQTTTGDRTRKKAAADVFDEIAYRDELVKIGIDLEAATLVASKTAQAHRDNAKARKQQQTTVGDIITNNNIPGRLREHSDEWKRHEPAAKAMAEERAKIAAARAKQIDIAGDFSIDLSIAPNFPLLARVQAEARARAAELAGRPAAKAKKPVKLFTTPDDKPAPSAAVARRRVETPIKAVYDYRYSELPERVRDLIEAHLAIEAEDAKSAGTLGFMTRALAIATLPHRKLAEDRFVRKNGDFTLTMLTAHPEGLPYGTLPRLLLTWVATEAVQKKERVLSLGNTLASYLNELGLHNTGGKRGDITRLKHAMTTLFSAIISCRYEGRDSWALQNVLLADKVEWWQPQDAAEAGAWQSRLQLSEPFFQECLDHPLPVDMRAMKVLRQSPLALDIYVWLTHRMSYLSKRTTIPWVSLSGQFGAGYAVNDQGLRDFKRAFLRELKNVVAIYPEAKLSESRNGLVLYPSPTHVLPDNSPKQPSLPF**

**H_pCAR1 (Rep3_N 142 aa: 17-158; Rep3_C 59 aa: 169-227)**

**MSNLPKPAKPEPKPLRVTKSNTLITASYRLTLNEQRLILAAISKLDPRRPMPKKVSVSAVDYSDIYGVQLRHAYEQMKVAADELYERDIKTFDGSGMERKRWVDRAKYLDGEGRVELSFTIHVMPYLTMLYSKVTSYDLRRVACLDSSHSFRLFEMLMQFRKTGWAYIEVESLRVALGLSDAYQRFNNLRQRVIDPAVAELKTKSNLDVSYELRREGRKVIAIKFTFCDLAQLPLNLELDPEDLPPLPEYDPAEESEWLITKPLGELAEAQEAGLLFAVSAEEGEQPD**

These analyses provide a consistent domain framework that underpins the comparative and structural interpretations presented in Figure 3 and Figures S6–S7.

**Supplemental Figure Legends**

**Figure S1. Methods to identify RIP genes of plasmids.**
Prediction of RIP genes were conducted by sequence-based search (BLAST, <https://blast.ncbi.nlm.nih.gov/Blast.cgi>) or protein structure-based search using AlphaFold 3 (<https://alphafoldserver.com/about>) and Foldseek (<https://search.foldseek.com/search>).

**Figure S2. Genetic structure of mini-replicon of PInc-2 (Rms139), PInc-5 (Rms163), PInc-10 (pPAB546), PInc-11 (RP1-1), PInc-12 (R716), Pinc-17 (pQBR103) and PInc-18 (pPT23-C1_1) plasmids.**
 (A) The mini-replicon of each plasmid was constructed by using DNA regions shown in a red arrow (replication initiation protein gene) with its promoter (gray box) and a gray solid line (*oriV*).
Putative iterons, palindromes, and AT-rich regions are shown by short red arrows, short yellow arrows, and blue boxes below the *oriV* region, respectively. Putative DnaA boxes are shown by black arrowheads in the *oriV* region. (B) Consensus sequences of putative iterons are shown. (C) Transformation efficiency of *Pseudomonas aeruginosa* PAO1, *P. putida* KT2440 or *P. fluorescens* SBW25 with the mini-replicons are shown.

**Figure S3. Homologous relationships between 251 non-redundant RIPs of 15 PInc groups.** %-identities of all-against-all BLASTp among the 251 non-redundant RIPs were visualised. The heatmap and the dendrograms were generated using R v3.6.1 and R packages dendextend and gplots via CPAP v0.3.0 (https://github.com/yosuken/CPAP) with a option (“--clust-method complete”) to perform clustering with complete-linkage method.

**Figure S4.The procedure of phylogenetic reconstruction of RIPs.**
The detailed procedure for (A) homolog collection, (B) phylogenetic reconstruction, (C) grouping of RIPs. The grouping started from 15 PInc groups, and the groups were sequentially merged and renamed. The occasions of group rearrangement are indicated by stars. The detailed path of group rearrangement is shown in (C).

**Figure S5. Remote homology identification between 14 orthologous groups of RIPs.**
(A) %-probability and identified regions of all-against-all HHblits between 14 orthologous groups of RIPs. Cells are filled with red if the %-probability is ≥80%, and filled with yellow if the %-probability is ≥50%. (B) Detailed procedures for extraction of homologous regions for each group of sWH and dWH RIPs based on the HHblits result.

**Figure S6. Predicted 3D structures of RIPs of PInc groups plasmids**
(A) Predicted structures of RIPs of PInc-1 (RepP-1A (=TrfA33) of RK2/RP4), PInc-2 (RepP-2A of Rms139), PInc-3 (RepP-3A of pRA1), PInc-4 (RepP-4A (=RepC) of pRSF1010), PInc-7 (RepP-7A of pCAR1), PInc-12 (RepP-12A of R716), PInc-15 (RepP-15A of pSN1104-11), PInc-16 (RepP-16A of pSN1216-29), PInc-17 (RepP-17A of pQBR103), and PInc-18 (RepP-18A of pSTY). The crystal structure (PDB: 2NRA) of Pi protein of IncX plasmid, R6K, interacted with iteron-containing dsDNA is shown as reference, of which structure is elucidated by X-ray crystallography. The WH domains of Pi are shown in pink and blue (WH1: 29-139 aa; WH2: 164-263 aa). (B) Predicted structures of RIPs of PInc-5 (RepP-5A of Rms163), PInc-9 (RepP-9A of pWW0), PInc-10 (RepP-10A of pPAB546), and PInc-11 (RepP-11A of RP1-1). The crystal structure (subunit B in PDB: 4PT7) of Rep19A proteins of *Staphylococcus* plasmid, pCA-347, is shown as reference, of which structure is elucidated by X-ray crystallography. The WH domain of Rep19A is shown in blue (WH: 29-111 aa).

**Figure S7. Predicted three-dimensional structures of RIPs and their oriV of plasmids in PInc groups.**

Only the WH domains extracted from the predicted structures of RIPs shown in Figure S6 are displayed. (A) The binding model of dWH with DNA was generated with reference to the crystal structure of the Pi protein of the IncX plasmid R6K (PDB: 2NRA). (B)

An identity matrix among dWH domains is shown. (C) The sWH dimer model was constructed with reference to the crystal structure of the Rep19A protein from the Staphylococcus plasmid pCA-347 (subunit B in PDB: 4PT7). The DNA-binding model was generated with reference to the crystal structure of the Pi protein of the IncX plasmid R6K (PDB: 2NRA). (D) An identity matrix among sWH domains is shown.

**Figure S8. Multiple sequence alignment used for phylogenetic tree reconstruction.**

Multiple sequence alignment of the conserved WH region used for phylogenetic reconstruction. To visualize sequence features flanking the conserved WH region, each sequence was extended by up to 300 amino acids on both the N- and C-terminal sides.

**Figure S9. Subclade-level summary of structural, compositional, taxonomic, ecological, and functional features across the WH RIP phylogeny.**

Subclade-level distributions and compositions of plasmid- and RIP-associated features across the 42 WH RIP subclades. From left to right, panels show the number of associated plasmids in PLSDB and IMG/PR, RIP amino-acid length, plasmid length, RIP GC content, plasmid GC content, RIP isoelectric point (pI), plasmid proteome pI, Pfam annotation profiles, taxonomic composition in PLSDB and IMG/PR, environmental composition in IMG/PR, MOB and MPF classes, antimicrobial resistance (AMR) gene counts, defense system counts, and resistant metal counts. Boxplots summarize distributions of continuous variables for clade-associated RIPs within each subclade, whereas stacked bars show the proportional composition of categorical features among associated plasmids or RIPs. Plasmids were counted as unique entries within each subclade.

**Figure S10.** **Detection coverage and assignment multiplicity in the PLSDB dataset.**
(A) Overlap between plasmids with a PlasmidFinder assignment and those with a WHRepDB and/or PseudomonasRepDB assignment. Circle and overlapping areas are proportional to counts. Plasmids without a hit in any of the three databases are listed separately below the circles. Percentages use all 59,895 plasmids as the denominator and are displayed to one decimal place. (B) Detection coverage of PlasmidFinder, WHRepDB, PseudomonasRepDB, and the combined detection by WHRepDB and/or PseudomonasRepDB. Each annotation gives the detected count divided by all 59,895 plasmids, followed by the percentage displayed to one decimal place. (C) Plasmids with multiple distinct assignments, evaluated separately within each database at four levels: PlasmidFinder types, WHRepDB numbered subclades, WHRepDB clades A–H, and PseudomonasRepDB types. Distinct assignments are counted once per plasmid within each level. WHRepDB subclades represent finer classifications within clades A–H; for example, H13 and H22 are distinct subclades belonging to the same clade H. Percentages use the corresponding database-positive population as the denominator. The four metrics are not additive across databases. Multiple reported assignments do not necessarily indicate multiple independent replication loci.

**Figure S11.** **Plasmid-level co-occurrence of PlasmidFinder types and WHRepDB subclades.**
The heatmap shows 69 PlasmidFinder types detected in at least 100 plasmids positive for both PlasmidFinder and WHRepDB, together with two additional types, rep1 and IncP6, yielding 71 rows arranged in two side-by-side blocks. The complete analysis included 23,322 unique plasmids positive for both databases and 226 PlasmidFinder types. Each cell indicates the number of plasmids carrying the corresponding PlasmidFinder type and WHRepDB subclade or unnumbered category. Cell colours represent the percentage of plasmids relative to the corresponding PlasmidFinder row denominator (Plasmids (%)). The row denominators, labelled "WHRepDB hits (n)", indicate the number of distinct plasmids carrying the specified PlasmidFinder type and at least one WHRepDB hit, rather than the number of sequence alignments. Plasmids with multiple WHRepDB assignments may contribute to multiple columns; therefore, row percentages may sum to more than 100%. The horizontal axis, labelled "WHRepDB subclade", also includes unnumbered categories: B and H indicate clade-only assignments, whereas SC1 and SC2 correspond to SC_CP031270 and SC_pKK5, respectively. The complete crosswalk atlas and accompanying tables include all 226 PlasmidFinder types. Plasmid-level co-occurrence does not necessarily indicate detection of the same replication locus by both databases.

**Figure S12.** **WHRepDB assignment outcomes in plasmids with a single PlasmidFinder type.**
The heatmaps summarise 18,493 plasmids with a single distinct PlasmidFinder type, encompassing all 113 types represented by at least 20 plasmids. Row labels indicate the number of plasmids (n) for each type. Each plasmid is assigned to one of four mutually exclusive categories: a single numbered ‘WHRepDB subclade’, ‘No WHRepDB’, Multiple subclades’, or ‘N/A’. ‘No WHRepDB’ indicates the absence of a WHRepDB hit, whereas ‘Multiple subclades’ indicates hits assigned to two or more distinct numbered WHRepDB subclades. ‘N/A’ indicates at least one WHRepDB hit without a numbered subclade assignment and takes priority over Multiple subclades when both conditions are met. Repeated assignments to the same subclade are counted only once. Cell colours represent the percentage of plasmids in each category relative to the corresponding row total (n). Percentages are displayed to one decimal place, except for 100%, which is shown without decimals; nonzero values below 0.1% are indicated as <0.1%. Of the 18,493 plasmids, 12,006 were assigned to a single numbered WHRepDB subclade, 5,645 had no WHRepDB hit, 839 were assigned to multiple numbered subclades, and three had at least one unnumbered assignment. These categories describe the distribution of reported assignments and do not represent validated detection sensitivity or demonstrate equivalence of replication loci. Complete counts and percentages are provided in the accompanying data workbook.

**Figure S13.** **Direct matches between reference database entries and WHRepDB.**
(A) Reference entries with at least one reported WHRepDB match. Bar lengths indicate the percentage of source reference entries with a WHRepDB hit. End labels show the number of matched entries divided by the total number of input entries, followed by the percentage: PlasmidFinder, 193/488 (39.5%); AcinetobacterPlasmidTyping, 457/555 (82.3%); and PseudomonasRepDB, 345/404 (85.4%). (B) WHRepDB clade composition among matched reference entries from these three databases, divided into 14 reference groups. The nine PlasmidFinder rows correspond to Enterobacteriales, Inc18, NT_Rep, Rep_trans, Rep1, Rep2, Rep3, RepA_N, and RepL. AcinetobacterPlasmidTyping is divided into Rep1, Rep3, and PriCT, whereas PseudomonasRepDB is divided into Other and PInc-6. For each row, the displayed fraction indicates the number of WHRepDB-positive reference entries divided by the total number of input entries. Heatmap cells show the number and percentage of matched entries assigned to each WHRepDB clade, using the number of WHRepDB-positive entries in the corresponding row as the denominator. Percentages are displayed to one decimal place, except for 100%, which is shown without decimals. Cells with no matches are indicated by a dash (–). N/A denotes matched entries that could not be assigned to any of the WHRepDB clades A–H; no such entries were observed in the three databases shown. Identical query–WHRepDB label combinations were counted once. Matches were reported at a minimum of 90% nucleotide identity and 60% overlap. These results represent reference-sequence matches rather than plasmid-level detection coverage or experimentally validated equivalence of replication loci.

**Figure S14. Identification of ICEpMG26 (previously assigned as IncP-13 plasmid) in the chromosome of *Pseudomonas aeruginosa*.**
(A) The illustration of how the primer sets (blue arrows) for detecting circular forms of the ICEpMG26 were designed. The 49-bp direct repeats are shown in orange. (B) Results of agarose gel electrophoresis for the PCR products to detect the circular form of ICEpMG26.

**Figure S15. Genome comparison of ICEpMG26 and pKLC102.**
Complete sequences of ICEpMG26 and pKLC102 were aligned. CDSs and their predicted functions (green for conjugation, yellow for maintenance, blue for genes related to mobile genetic elements, pink for antimicrobial resistance genes and magenta for other genes). Homologous regions are indicated by frame areas.

**Figure S16. A genetic map of Rms163, a PInc-5 plasmid.**
Innermost rings represent the GC contents (black) and GC skew (purple and green). The *oriV* candidate regions predicted by GC skew are shown by red circles. The CDSs of Rms163 are shown by arcs in the outer rings of the GC skew plot.

**Figure S17. Comparisons of genomes of PInc-5 plasmids.**
Comparisons of the whole genetic structure of PInc-5 plasmids, Rms163 and pPALA50 (CP111035). CDSs and their predicted functions (red for replication, green for conjugation, yellow for maintenance in PInc-5 backbone, blue for genes related to mobile genetic elements and magenta for other genes). Homologous regions are indicated by frame areas.

**References**

1.	Shintani, M., Suzuki, H., Nojiri, H. & Suzuki, M. Precise classification of antimicrobial resistance-associated IncP-2 megaplasmids for molecular epidemiological studies on *Pseudomonas* species. *J. Antimicrob. Chemother.* **77**, 1203–1205 (2022).

2.	Davies, S. & Krishnapillai, V. DNA sequence analysis of the replication region of the *Pseudomonas aeruginosa* plasmid R91-5. *J. Genet.* **69**, 101–112 (1990).

3.	Cain, D. & Holloway, B. W. Prime plasmids derived from the IncP-10 plasmid R91-5 in *Pseudomonas putida*. *FEMS Microbiol. Lett.* **24**, 97–101 (1984).

4.	Robertson, J. & Nash, J. H. E. MOB-suite: software tools for clustering, reconstruction and typing of plasmids from draft assemblies. *Microb. Genom.* **4**, e000206 (2018).

5.	Jacoby, G. A. Plasmid determined resistance to carbenicillin and gentamicin in *Pseudomonas aeruginosa*. In *Plasmids and Transposons: Environmental Effects and Maintenance Mechanisms* (eds Stuttard, C. & Rozee, K. R.) 83–96 (Academic Press, 1980).

6.	Bradley, D. E. Specification of the conjugative pili and surface mating systems of *Pseudomonas* plasmids. *J. Gen. Microbiol.* **129**, 2545–2556 (1983).

7.	Kawalek, A. et al. Genome sequence of *Pseudomonas aeruginosa* PAO1161, a PAO1 derivative with the ICEPae1161 integrative and conjugative element. *BMC Genomics* **21**, 14 (2020).

8.	Klockgether, J., Reva, O., Larbig, K. & Tümmler, B. Sequence analysis of the mobile genome island pKLC102 of *Pseudomonas aeruginosa* C. *J. Bacteriol.* **186**, 518–534 (2004).

9.	Shintani, M., Yoshida, T., Habe, H., Omori, T. & Nojiri, H. Large plasmid pCAR2 and class II transposon Tn4676 are functional mobile genetic elements to distribute the carbazole/dioxin-degradative car gene cluster in different bacteria. *Appl. Microbiol. Biotechnol.* **67**, 370–382 (2005).

10.	Kitao, T., Kubori, T. & Nagai, H. Recent advances in structural studies of the *Legionella pneumophila* Dot/Icm type IV secretion system. *Microbiol. Immunol.* **66**, 67–74 (2022).

11.	Hayakawa, M. et al. Hitherto-unnoticed self-transmissible plasmids widely distributed among different environments in Japan. *Appl. Environ. Microbiol.* **88**, e01114-22 (2022).

12.	Pachulec, E. & van der Does, C. Conjugative plasmids of *Neisseria gonorrhoeae*. *PLoS ONE* **5**, e9962 (2010).

13.	Cazares, A. et al. A megaplasmid family driving dissemination of multidrug resistance in *Pseudomonas*. *Nat. Commun.* **11**, 1370 (2020).

14.	Jiang, X. et al. Plasmids of novel incompatibility group IncpRBL16 from *Pseudomonas* species. *J. Antimicrob. Chemother.* **75**, 2093–2100 (2020).

15.	Hidaka, M., Kobayashi, T., Takenaka, S., Takeya, H. & Horiuchi, T. Purification of a DNA replication terminus (ter) site-binding protein in *Escherichia coli* and identification of the structural gene. *J. Biol. Chem.* **264**, 21031–21037 (1989).

16.	Ambrose, S. J., Harmer, C. J. & Hall, R. M. Compatibility and entry exclusion of IncA and IncC plasmids revisited: IncA and IncC plasmids are compatible. *Plasmid* **96–97**, 7–12 (2018).

17.	Loftie-Eaton, W. & Rawlings, D. E. Diversity, biology and evolution of IncQ-family plasmids. *Plasmid* **67**, 15–34 (2012).

18.	Robinson, L. A., Collins, A. C. Z., Murphy, R. A., Davies, J. C. & Allsopp, L. P. Diversity and prevalence of type VI secretion system effectors in clinical *Pseudomonas aeruginosa* isolates. *Front. Microbiol.* **13**, 1042505 (2023).

19.	Haines, A. S., Cheung, M. & Thomas, C. M. Evidence that IncG (IncP-6) and IncU plasmids form a single incompatibility group. *Plasmid* **55**, 210–215 (2006).

20.	Kulińska, A., Czeredys, M., Hayes, F. & Jagura-Burdzy, G. Genomic and functional characterization of the modular broad-host-range RA3 plasmid, the archetype of the IncU group. *Appl. Environ. Microbiol.* **74**, 4119–4132 (2008).

21.	Rhodes, G. et al. Complete nucleotide sequence of the conjugative tetracycline resistance plasmid pFBAOT6, a member of a group of IncU plasmids with global ubiquity. *Appl. Environ. Microbiol.* **70**, 7497–7510 (2004).

22.	Shintani, M. et al. Characterization of the replication, maintenance, and transfer features of the IncP-7 plasmid pCAR1, which carries genes involved in carbazole and dioxin degradation. *Appl. Environ. Microbiol.* **72**, 3206–3216 (2006).

23.	Yano, H. et al. Complete sequence determination combined with analysis of transposition/site-specific recombination events to explain genetic organization of IncP-7 TOL plasmid pWW53 and related mobile genetic elements. *J. Mol. Biol.* **369**, 11–26 (2007).

24.	Suenaga, H. et al. Insights into the genomic plasticity of *Pseudomonas putida* KF715, a strain with unique biphenyl-utilizing activity and genome instability properties. *Environ. Microbiol. Rep.* **9**, 589–598 (2017).

25.	Bonnin, R. A. et al. Molecular characterization of OXA-198 carbapenemase-producing *Pseudomonas aeruginosa* clinical isolates. *Antimicrob. Agents Chemother.* **62**, e02496-17 (2018).

26.	Shintani, M., Suzuki, H., Nojiri, H. & Suzuki, M. Reconsideration of the previously classified incompatibility groups of plasmids, IncP-1 and IncP-11. *Environ. Microbiol.* **25**, 1071–1076 (2023).

27.	Yanagiya, K. et al. Novel self-transmissible and broad-host-range plasmids exogenously captured from anaerobic granules or cow manure. *Front. Microbiol.* **9**, 2602 (2018).

28.	Tokuda, M. et al. Transconjugant range of PromA plasmids in microbial communities is predicted by sequence similarity with the bacterial host chromosome. *Microb. Genom.* **9**, 001043 (2023).

29.	Botelho, J. et al. Combining sequencing approaches to fully resolve a carbapenemase-encoding megaplasmid in a *Pseudomonas shirazica* clinical strain. *Emerg. Microbes Infect.* **8**, 1186–1194 (2019).

30.	Zhu, J. et al. Identification of TMexCD-TOprJ-producing carbapenem-resistant Gram-negative bacteria from hospital sewage. *Drug Resist. Updat.* **70**, 100989 (2023).

31.	Chen, F. et al. VIM-encoding IncpSTY plasmids and chromosome-borne integrative and mobilizable elements (IMEs) and integrative and conjugative elements (ICEs) in *Pseudomonas*. *Ann. Clin. Microbiol. Antimicrob.* **21**, 10 (2022).

32.	Rice, P., Longden, I. & Bleasby, A. EMBOSS: the European Molecular Biology Open Software Suite. *Trends Genet.* **16**, 276–277 (2000).

33.	Steinegger, M. & Söding, J. MMseqs2 enables sensitive protein sequence searching for the analysis of massive data sets. *Nat. Biotechnol.* **35**, 1026–1028 (2017).

34.	Altschul, S. F. et al. Gapped BLAST and PSI-BLAST: a new generation of protein database search programs. *Nucleic Acids Res.* **25**, 3389–3402 (1997).

35.	Katoh, K. & Standley, D. M. MAFFT multiple sequence alignment software version 7: improvements in performance and usability. *Mol. Biol. Evol.* **30**, 772–780 (2013).

36.	Eddy, S. R. Accelerated profile HMM searches. *PLoS Comput. Biol.* **7**, e1002195 (2011).

37.	Woodcroft, B. J., Boyd, J. A. & Tyson, G. W. OrfM: a fast open reading frame predictor for metagenomic data. *Bioinformatics* **32**, 2702–2703 (2016).

38.	Steinegger, M. et al. HH-suite3 for fast remote homology detection and deep protein annotation. *BMC Bioinformatics* **20**, 473 (2019).

39.	Wegrzyn, K. et al. Defining a novel domain that provides an essential contribution to site-specific interaction of Rep protein with DNA. *Nucleic Acids Res.* **49**, 3394–3408 (2021).

40.	Komori, H. et al. Crystal structure of a prokaryotic replication initiator protein bound to DNA at 2.6 Å resolution. *EMBO J.* **18**, 4597–4607 (1999).

41.	Schumacher, M. A. et al. Mechanism of staphylococcal multiresistance plasmid replication origin assembly by the RepA protein. *Proc. Natl Acad. Sci. USA* **111**, 9121–9126 (2014).

42.	Stover, C. K. et al. Complete genome sequence of *Pseudomonas aeruginosa* PAO1, an opportunistic pathogen. *Nature* **406**, 959–964 (2000).

43.	Royle, P. L., Matsumoto, H. & Holloway, B. W. Genetic circularity of the *Pseudomonas aeruginosa* PAO chromosome. *J. Bacteriol.* **145**, 145–155 (1981).

44.	Rainey, P. B. & Bailey, M. J. Physical and genetic map of the *Pseudomonas fluorescens* SBW25 chromosome. *Mol. Microbiol.* **19**, 521–533 (1996).

45.	Bagdasarian, M. et al. Specific-purpose plasmid cloning vectors. II. Broad host range, high copy number, RSF1010-derived vectors, and a host-vector system for gene cloning in *Pseudomonas*. *Gene* **16**, 237–247 (1981).

46.	Nelson, K. E. et al. Complete genome sequence and comparative analysis of the metabolically versatile *Pseudomonas putida* KT2440. *Environ. Microbiol.* **4**, 799–808 (2002).

47.	Sawada, Y., Yaginuma, S., Tai, M., Iyobe, S. & Mitsuhashi, S. Plasmid-mediated penicillin beta-lactamases in *Pseudomonas aeruginosa*. *Antimicrob. Agents Chemother.* **9**, 55–60 (1976).

48.	Sagai, H. et al. Classification of R plasmids by incompatibility in *Pseudomonas aeruginosa*. *Antimicrob. Agents Chemother.* **10**, 573–578 (1976).

49.	Summers, A. O. & Jacoby, G. A. Plasmid-determined resistance to boron and chromium compounds in *Pseudomonas aeruginosa*. *Antimicrob. Agents Chemother.* **13**, 637–640 (1978).

50.	Kawakami, Y., Mikoshiba, F., Nagasaki, S., Matsumoto, H. & Tazaki, T. Prevalence of *Pseudomonas aeruginosa* strains possessing R factor in a hospital. *J. Antibiot.* **25**, 607–609 (1972).

51.	Hedges, R. W. & Jacob, A. E. Mobilization of plasmid-borne drug resistance determinants for transfer from *Pseudomonas aeruginosa* to *Escherichia coli*. *Mol. Gen. Genet.* **140**, 69–79 (1975).

52.	Ingram, L., Sykes, R. B., Grinsted, J., Saunders, J. R. & Richmond, M. H. A transmissible resistance element from a strain of *Pseudomonas aeruginosa* containing no detectable extrachromosomal DNA. *J. Gen. Microbiol.* **72**, 269–279 (1972).

53.	Bryan, L. E., Semaka, S. D., van den Elzen, H. M., Kinnear, J. E. & Whitehouse, R. L. Characteristics of R931 and other *Pseudomonas aeruginosa* R factors. *Antimicrob. Agents Chemother.* **3**, 625–637 (1973).

54.	Yanisch-Perron, C., Vieira, J. & Messing, J. Improved M13 phage cloning vectors and host strains: nucleotide sequences of the M13mp18 and pUC19 vectors. *Gene* **33**, 103–119 (1985).

55.	Kovach, M. E. et al. Four new derivatives of the broad-host-range cloning vector pBBR1MCS, carrying different antibiotic-resistance cassettes. *Gene* **166**, 175–176 (1995).
